# Discovery of a Small-Molecule mTORC1 Pathway Activator that Enhances Adoptive T-Cell Therapy via Immuno-metabolic Reprogramming of CD8^+^ T Cells

**DOI:** 10.64898/2026.09.11.750909

**Authors:** Dipika Sarkar, Puspendu Ghosh, Sunny Goon, Ishita Sarkar, Himadri Sekhar Sarkar, Deborpita Sarkar, Anupam Goutam, Shaun Mahanti, Sandip Paul, Shilpak Chatterjee, Arindam Talukdar

**Affiliations:** Department of Organic and Medicinal Chemistry, CSIR-Indian Institute of Chemical Biology, 4 Raja S. C. Mullick Road, Kolkata 700032, WB (India); Department of Cancer Biology and Inflammatory Disorders, CSIR-Indian Institute of Chemical Biology, CN6, Sector V, Salt Lake, Kolkata 700091, WB (India); Institute for Bioinformatics and Medical Informatics, University of Tübingen, Sand 14, 72076 Tübingen, Germany; 3 International Max Planck Research School “From Molecules to Organisms”, Max Planck Institute for Biology Tübingen, Max-Planck-Ring 5, 72076, Tübingen, Germany; Center for Health Science and Technology, JIS Institute of Advanced Studies and Research, JIS University, Kolkata, India; Academy of Scientific and Innovative Research (AcSIR), Ghaziabad, 201002, India

## Abstract

Adoptive cell therapy (ACT) has transformed cancer immunotherapy; however, its clinical efficacy remains limited by progressive T cell dysfunction and inadequate metabolic fitness acquired during ex vivo expansion and within the tumor microenvironment. Here, we report the discovery of AS-3, a first-in-class small-molecule activator that enhances the therapeutic competence of CD8⁺ T cells through RAPTOR-dependent mTORC1 activation. Identified by phenotypic screening of an in-house library of pharmacologically relevant scaffolds and subsequent medicinal chemistry optimization, AS-3 markedly increased effector cytokine production while preserving T cell viability. Mechanistically, AS-3 enhanced phosphorylation of mTORC1 downstream effectors, promoted glycolytic and mitochondrial metabolism, and sustained pathway activity under rapamycin-mediated inhibition. Transcriptomic profiling of activated human CD8⁺ T cells revealed coordinated enrichment of mTORC1 signalling, oxidative phosphorylation, glycolysis, proliferative programs, and cytotoxic effector gene networks, consistent with comprehensive immunometabolic reprogramming. Genetic silencing of RAPTOR attenuated both mTORC1 signalling and cytokine induction, establishing pathway dependency. Ex vivo conditioning with AS-3 significantly improved antitumor efficacy in two independent adoptive T-cell therapy models, accompanied by enhanced persistence, reduced exhaustion, and superior effector function. Together, these findings establish pharmacological activation of the RAPTOR-mTORC1 axis as a strategy to generate metabolically resilient T cells and provide a clinically translatable approach to improving the durability and efficacy of next-generation ACT.

## Introduction

Cancer immunotherapy has transformed the treatment landscape across a broad spectrum of malignancies (1, 2). Among these approaches, adoptive T cell therapy (ACT), which involves the ex vivo programming and subsequent infusion of tumor epitope-reactive autologous T cells into tumor-bearing hosts, has emerged as a particularly promising strategy for patients with advanced disease (3). In recent years, chimeric antigen receptor (CAR) T cell therapy, utilizing autologous or allogeneic T cells engineered to express CAR constructs, has achieved regulatory approval and demonstrated remarkable clinical success, particularly in relapsed or refractory (R/R) hematologic malignancies (4, 5). However, despite these advances, a substantial proportion of patients fail to achieve durable responses or experience only transient clinical benefit. This limitation is largely attributed to the rapid onset of T cell exhaustion, which compromises CAR T cell persistence and effector function (5, 6). Therefore, strategies that sustain effector differentiation while preventing or delaying T cell exhaustion are critical to improving response durability in ACT.

The limited persistence and efficacy of adoptively transferred T cells arise from a complex interplay of intrinsic and extrinsic factors that collectively drive T cell dysfunction. Within the tumor microenvironment (TME), chronic antigen stimulation, metabolic stress, and immunosuppressive cytokines promote the progressive differentiation of T cells into an exhausted state (7–9). This dysfunctional phenotype is characterized by sustained expression of multiple co-inhibitory receptors, including PD-1, TIM-3, LAG-3, and CTLA-4, along with diminished proliferative capacity and impaired effector function (7). While extrinsic signals within the TME are key drivers of this process, accumulating evidence highlights a critical role for T cell-intrinsic programs in both the initiation and maintenance of exhaustion (7, 8, 10–12). These intrinsic mechanisms reinforce the stability and persistence of the dysfunctional state. Importantly, this evolving understanding presents a strategic opportunity in ACT, as the ex vivo activation and expansion phase offers a controlled window to reprogram T cell-intrinsic pathways before infusion (13). In this context, ex vivo modulation strategies using clinically approved agents, including monoclonal antibodies and small molecules, have emerged as an attractive approach to generate ACT products with enhanced functionality and improved resistance to exhaustion (14). Several signalling and metabolic pathways, such as PI3K-AKT, Wnt/β-catenin, BTK, IDH2, HK2, etc, are currently being targeted during ex vivo expansion to enhance therapeutic efficacy; however, an optimized and clinically standardized approach remains to be established (15–19).

Emerging evidence indicates that metabolic fitness is a central determinant of ACT efficacy, enabling T cells to adapt to the hostile conditions of the TME (20, 21). While aerobic glycolysis supports immediate effector functions, robust mitochondrial fitness is essential for long-term persistence and sustained anti-tumor activity (22–24). Notably, tumor-reactive T cells optimized for mitochondrial metabolism exhibit superior persistence and enhanced tumor clearance (12, 23). Therefore, the development of small-molecule interventions that simultaneously enhance effector function and metabolic fitness represents a compelling strategy to improve ACT outcomes. Incorporating such approaches during the ex vivo manufacturing phase may yield more durable and therapeutically effective T cell products for cancer immunotherapy.

In the present study, to discover small molecules capable of enhancing T-cell fitness in a physiologically relevant manner, we screened an in-house library of pharmacologically active scaffolds without pre-imposed target bias. This phenotypic screening strategy enables the identification of a hit compound that modulates complex pathways critical for effective T-cell responses.

## Results

### Identification of small molecules capable of inducing the functional and metabolic fitness of CD8⁺ T cells

Given that T cells with enhanced effector function and metabolic activity elicit durable anti-tumor responses, we screened for small molecules capable of augmenting these properties during ex vivo activation of T cells prior to infusion into tumor-bearing hosts. To this end, we performed a focused screen of an in-house library enriched for pharmacophores previously associated with immunomodulatory activity (25–29) using primary CD8⁺ T cells isolated from C57BL/6 mice and activated in the presence or absence of the indicated compounds. Effector function was evaluated by measuring intracellular IFN-γ and TNF-α as primary readouts because they represent canonical outputs of activated CD8⁺ T cells and are closely linked to productive TCR signalling, co-stimulation, and downstream metabolic engagement (Figure 1A). Importantly, these cytokines are not only markers of activation but also key mediators of cytotoxic function, making them particularly well-suited for early-stage functional screening (20, 30).

**Figure 1.**
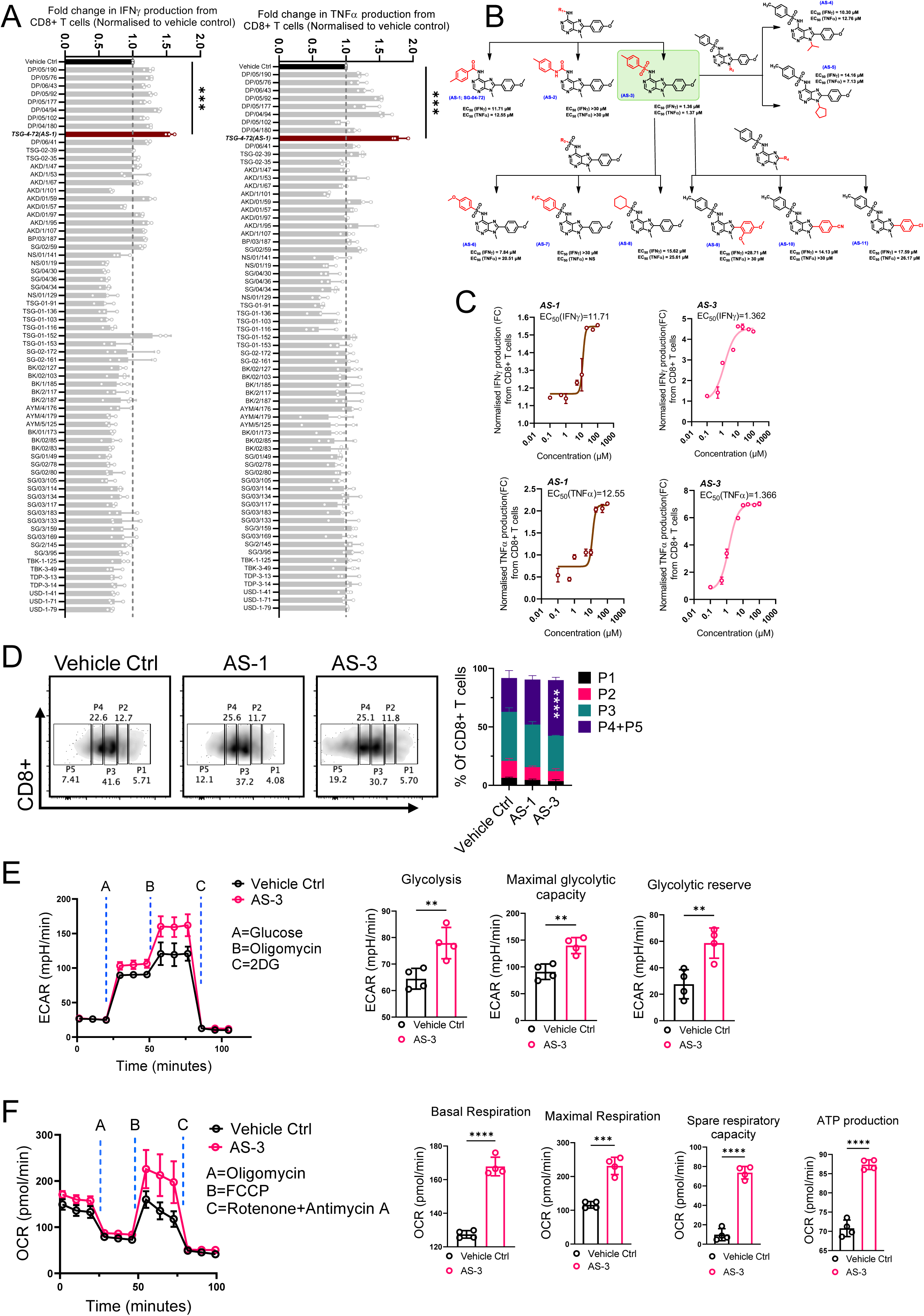
SAR-guided identification of AS-3 as a modulator of CD8⁺ T cell function and metabolic features. (A) High-throughput screening displaying fold changes in IFN-*γ* and TNF-*α* production by CD8+ T cells relative to vehicle control across a library of small-molecule compounds. Data are representative of three independent experiments. (B) Schematic visualization for structure-activity relationship (SAR) development showing chemical structures and EC_50_values for IFN-*γ* and TNF-*α* modulation across key analogs (AS-1 to AS-11). NS = non-significant. (C) Dose-response curves illustrating normalized IFN-*γ* and TNF-*α* production (fold change) in CD8+ T cells treated with compounds AS-1 and AS-3, with indicated EC_50_ values for each cytokine. (D) Flow cytometry analysis showing CTV dilution and quantification of CD8+ T cell proliferation following treatment with vehicle control, AS-1, or AS-3. Data are cumulative of four independent experiments. (E) Glycolytic function assessed by real-time extracellular acidification rate (ECAR) measurements following sequential addition of glucose, oligomycin, and 2-deoxy-D-glucose (2-DG). Bar graphs display quantified glycolysis, glycolytic reserve, and maximal glycolytic capacity for vehicle control and AS-3-treated CD8+ T cells. Data are cumulative of four independent experiments. (F) Mitochondrial respiration measured by real-time oxygen consumption rate (OCR) profiles following sequential treatment with oligomycin, FCCP, and rotenone plus antimycin A. Adjacent bar graphs quantify basal respiration, maximal respiration, spare respiratory capacity, and ATP-linked respiration in the respective group from four independent experiments . Data are shown as mean ± SEM. ns, not significant; *p < 0.05, **p < 0.01, ***p < 0.001, ****p < 0.0001.

This screening strategy led to the identification of AS-1 (TSG-04-72) as an initial hit that enhanced cytokine production in activated CD8^+^ T cells (Figure 1A). Although AS-1 demonstrated measurable activity, its modest potency (EC_50_: 11.71 μM for IFNγ and 12.55 μM for TNFα) indicated scope for structural optimization. AS-1 comprises an adenine scaffold bearing an amide substituent at the C6 position, which was initially targeted for SAR exploration (Figure 1B). Replacement of the amide with a urea moiety was poorly tolerated, as AS-2 showed minimal activity. In contrast, conversion to sulfonamide analogues improved potency. Among these, the *p*-tolyl sulfonamide derivative AS-3 (TSG-02-56) showed the strongest activity, with EC₅₀ values of 1.36 μM for IFNγ and 1.37 μM for TNFα in activated human CD8⁺ T cells (Figure 1C and Figure S1A), representing an approximately 9-fold improvement in potency over AS-1.

We next investigated the tolerance of the scaffold to substitution at N9. Introduction of either an isopropyl group (AS-4; EC_50_ = 10.30 μM for IFNγ and 12.76 μM for TNFα) or a cyclopentyl group (AS-5; EC_50_ = 14.16 μM for IFNγ and 7.13 μM for TNFα) resulted in reduced potency relative to AS-3, indicating that increased steric bulk at N9 did not provide additional benefit. Modification of the distal aryl group of the sulfonamide similarly revealed a relatively narrow SAR. Replacement of the *p*-methyl substituent with methoxy (AS-6) reduced activity (EC_50_ = 7.84 μM for IFNγ and 20.51 μM for TNFα), whereas the *p*-trifluoromethyl analogue AS-7 was poorly tolerated. Replacement of the aryl sulfonamide with an aliphatic sulfonamide (AS-8) was also unfavorable, yielding EC_50_ values of 15.62 μM for IFNγ and 25.61 μM for TNFα. Together, these modifications established the *p*-tolyl sulfonamide of AS-3 as the preferred substitution pattern within this series. Finally, we examined the tolerance of the adenine core to modification at C8. Introduction of either a cyano group (AS-10) or chloro substituent (AS-11) substantially reduced activity, with EC_50_ values of 14.13 and 17.59 μM for IFNγ and >30 and 26.17 μM for TNFα, respectively. The dimethoxy-substituted analogue AS-9 was even less active, exhibiting an EC_50_ of 28.71 μM for IFNγ and >30 μM for TNFα. Together, these results identified AS-3 as the most active analogue in the series and the compound selected for further mechanistic characterization (Figure 1B).

Notably, AS-3 showed minimal cytotoxicity in CD8^+^ T cells derived from human PBMC even at higher concentrations up to 50 µM (Figure S1B). Beyond cytokine enhancement, AS-2 treatment significantly increased the proliferative capacity of CD8^+^ T cells, compared to both control and AS-1 (Figure 1D). Together, this improved cytokine profile and proliferative potential suggest an efficient engagement of activation-associated signalling pathways in AS-3-treated T cells.

Given that the effector cytokine production and proliferative potential of T cells are intrinsically coupled to metabolic reprogramming, we next sought to determine whether AS-3 influences the metabolic fitness of CD8^+^ T cells. Accordingly, we assessed its effects on both glycolytic and mitochondrial metabolic programs as they collectively support the energetic and biosynthetic demands required for sustained effector function. Consistent with an activated phenotype, human CD8^+^ T cells treated with AS-3 displayed significantly increased uptake of the fluorescent glucose analogue 2-NBDG, indicating enhanced glucose utilization (Figure S1C). This observation was further supported by extracellular acidification rate (ECAR) measurements, which revealed a clear elevation in basal glycolysis (Figure 1E). Upon oligomycin treatment, which restricts mitochondrial ATP production and forces reliance on glycolysis, AS-3-treated cells exhibited a higher maximal glycolytic capacity. The increase in glycolytic reserve, together with the sensitivity to 2-deoxyglucose (2-DG), confirms that AS-3 promotes a glycolysis-dependent metabolic program with improved metabolic flexibility under stress conditions.

In addition to augmenting glycolytic activity, treatment with AS-3 induced a pronounced remodelling of mitochondrial architecture and function in CD8^+^ T cells, further reinforcing its role as a metabolic modulator. Cells exposed to AS-3 displayed a clear increase in mitochondrial mass, as evidenced by elevated MitoTracker Green fluorescence, alongside a concomitant rise in mitochondrial membrane potential measured by MitoTracker Deep Red staining (Figure S1D-E). These observations indicate not only an increase in mitochondrial content but also an improvement in mitochondrial activity. Functional interrogation using flux analysis revealed that these structural changes translated into enhanced OXPHOS (Figure 1F). AS-3-treated T cells exhibited a significant increase in basal oxygen consumption rate (OCR_Bas_), reflecting heightened mitochondrial activity under steady-state conditions. Upon uncoupling with FCCP, these cells achieved a substantially higher maximal respiratory capacity (OCR_Max_), indicating an improved ability of the electron transport chain to operate at peak efficiency when energy demand increases. Importantly, the spare respiratory capacity was also elevated, suggesting that AS-3 equips T cells with the metabolic reserve required to withstand bioenergetic stress, a feature closely associated with long-lasting memory T cells (31). Furthermore, ATP-linked respiration was enhanced following oligomycin treatment, indicating that the observed increase in oxygen consumption is functionally coupled to ATP production rather than uncoupled respiration. This enhancement underscores the ability of AS-3 to support efficient energy generation, which is essential for maintaining effector responses during prolonged activation. Taken together, these findings demonstrate that AS-3 orchestrates a coordinated metabolic program characterized by simultaneous enhancement of glycolytic flux and mitochondrial respiration. Such dual metabolic reinforcement establishes a bioenergetically robust and adaptable T cell state, capable of sustaining prolonged activation and function.

### Transcriptional remodelling of CD8⁺ T cells upon AS-3 treatment

To investigate the molecular basis underlying the enhanced functional and metabolic phenotype induced by AS-3, we performed RNA sequencing on activated human CD8⁺ T cells cultured in the presence or absence of AS-3. Principal component analysis revealed a clear separation between control and AS-3-treated samples, with PC1 explaining 98% of the total variance, indicating a robust and consistent transcriptional response to AS-3 treatment (Figure 2A). Gene set enrichment analysis (GSEA) demonstrated significant enrichment of HALLMARK_MTORC1_SIGNALING in AS-3-treated cells, indicating activation of the canonical mTORC1 transcriptional program (Figure. 2B-C). In addition, metabolic pathways associated with mitochondrial respiration, including HALLMARK_OXIDATIVE_PHOSPHORYLATION and respiratory electron transport, were enriched in the AS-3-treated condition, consistent with increased mitochondrial activity observed in our earlier metabolic flux assays. In parallel, pathways associated with cellular growth and proliferation, including MYC targets, E2F targets, and the G2M checkpoint, were enriched upon AS-3 treatment, suggesting increased biosynthetic activity and proliferative capacity. Notably, apoptotic signalling programs were comparatively reduced in AS-3-treated cells, consistent with a transcriptional state favoring survival and sustained activation.

**Figure 2.**
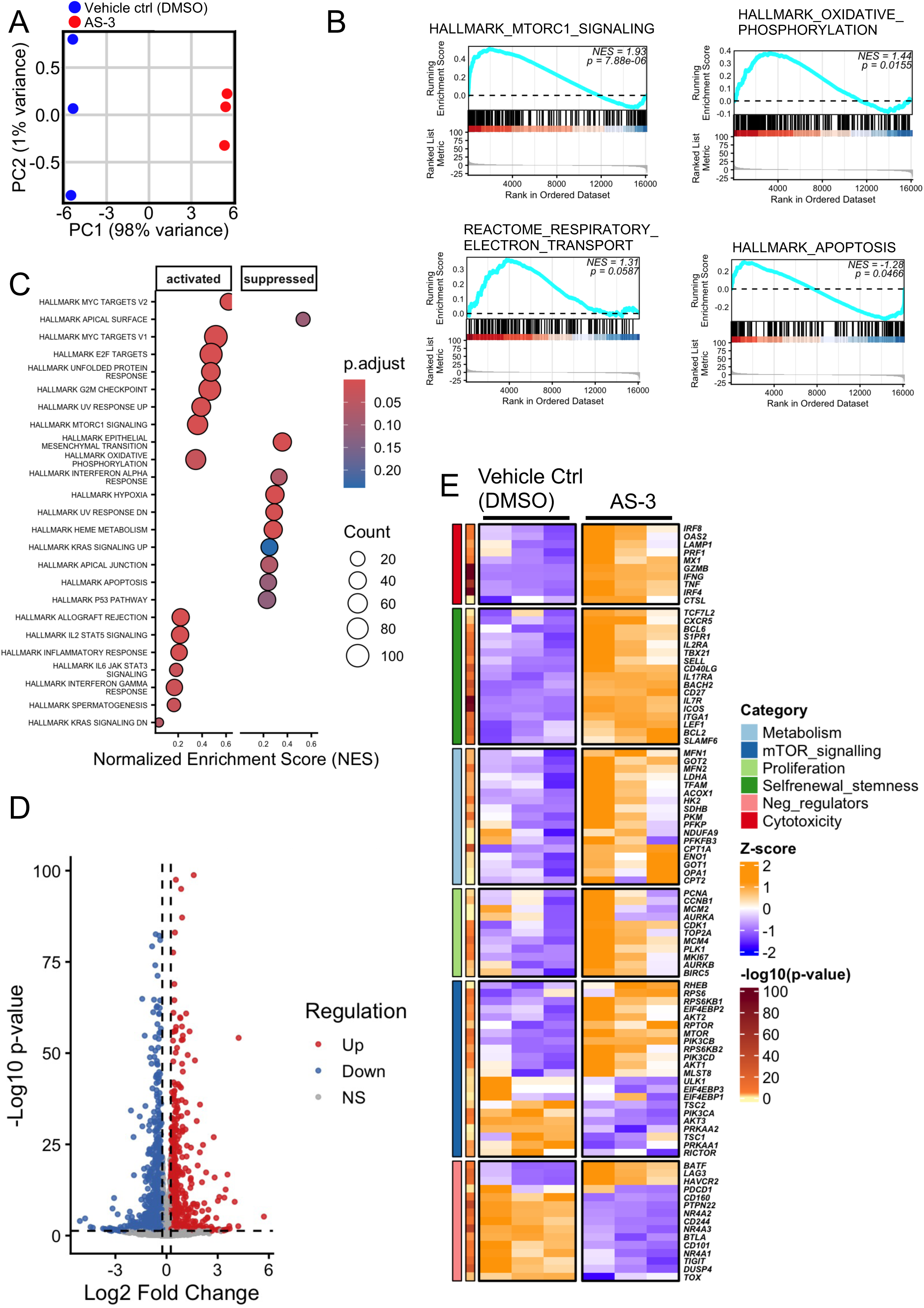
AS-3 induces transcriptional reprogramming of CD8⁺ T cells toward enhanced metabolic and functional fitness. (A) Principal component analysis (PCA) of RNA-seq profiles from activated human CD8⁺ T cells cultured with or without AS-3. (B) Representative GSEA enrichment plots showing enrichment of mTORC1 signaling ,oxidative phosphorylation, respiratory electron transport and hallmark apoptosis in AS-3 treated CD8⁺ T cells. (C) Bubble plot showing the top 25 enriched Hallmark pathways identified by GSEA in AS-3 treated versus control CD8⁺ T cells. Pathways are separated into positively (activated) and negatively (supressed) enriched gene sets based on the normalized enrichment score (NES). Bubble size represents the number of genes contributing to each pathway, and color indicates statistical significance (adjusted p-value). (D) Volcano plot showing differentially expressed genes between control and AS-3 treated CD8⁺ T cells. Red, upregulated; blue, downregulated; grey, not significant (NS). (E) Heatmap depicting expression of curated CD8⁺ T-cell gene modules related to cytotoxicity, metabolism, proliferation, self-renewal, and mTOR signaling in control and AS-3 treated cells (row Z-score).

Differential gene expression analysis further revealed broad transcriptional remodeling following AS-3 treatment, with 1053 genes significantly upregulated (log₂FC > 0.3, p < 0.05) and 1218 genes significantly downregulated (log₂FC < −0.3, p < 0.05) (Figure. 2D). To further define the functional programs associated with this transcriptional shift, we examined curated CD8⁺ T-cell gene modules encompassing cytotoxicity, metabolism, proliferation, self-renewal/stemness, negative regulatory pathways, and mTOR signaling (Figure 2E).

AS-3-treated CD8⁺ T cells displayed marked induction of effector and cytotoxicity-associated genes, including *IFNG, TNF, PRF1, GZMB*, and *MX1*, together with transcriptional regulators such as *IRF8* and *IRF4*, indicating acquisition of a highly activated effector state (32). Increased expression of *OAS2* and *LAMP1* further suggested enhanced inflammatory and antiviral-response programs (33). In parallel, genes associated with T-cell activation, trafficking, and memory-like differentiation, including *IL7R, SELL, BCL6, SLAMF6, IL2RA,* and *LEF1* were elevated, suggesting that AS-3 promotes an activated yet persistent effector phenotype rather than terminal dysfunction (32, 34).

AS-3 treatment also induced extensive metabolic rewiring characterized by coordinated upregulation of glycolytic, mitochondrial, and fatty acid metabolic programs. Genes associated with glycolysis and anabolic metabolism, including *HK2, LDHA, PFKFB3,* and *ENO1*, were significantly increased, consistent with enhanced glucose utilization required to sustain rapid proliferation and effector function. In parallel, AS-3-treated cells exhibited elevated expression of genes linked to mitochondrial fitness and oxidative metabolism, including *TFAM, MFN1, CPT1A, CPT2, GOT1, GOT2,* and *ACOX1*, indicating enhanced mitochondrial biogenesis, respiratory capacity, and metabolic flexibility. Enrichment of oxidative phosphorylation and respiratory electron transport pathways further supported the establishment of a highly bioenergetic state in AS-3-treated CD8⁺ T cells. Given that mitochondrial respiratory capacity, fatty acid oxidation, and mitochondrial fusion are critical determinants of memory formation, persistence, and resistance to exhaustion in CD8⁺ T cells, the increased expression of mitochondrial regulatory genes such as *CPT1A, MFN2,* and *TFAM* suggests that AS-3 promotes a metabolically resilient state capable of sustaining long-term effector function. Consistent with the enrichment of proliferative gene programs observed by GSEA, AS-3-treated cells showed elevated expression of multiple cell-cycle regulators and DNA replication-associated genes, including *MKI67, PCNA, CDK1, TOP2A, PLK1,* and *AURKB,* indicating enhanced proliferative fitness and cell-cycle progression. In parallel, the mTOR signalling module demonstrated coordinated upregulation of canonical pathway components and downstream effectors, including *RHEB, MTOR, RPTOR, RPS6, PIK3CD,* and *ULK1*, further supporting sustained activation of anabolic mTORC1 signalling in AS-3-treated cells.

In contrast, several genes associated with inhibitory or dysfunctional T-cell states, including *PDCD1, NR4A1, TIGIT, and TOX* were comparatively reduced following AS-3 treatment, suggesting that AS-3 limits the establishment of exhaustion-associated transcriptional programs despite promoting strong effector differentiation (35, 36). These data define a coordinated transcriptional program in AS-3-treated CD8⁺ T cells characterized by enhanced mTORC1 signalling, metabolic fitness, proliferation, and cytotoxic differentiation while simultaneously restraining inhibitory and apoptotic programs, thereby supporting the development of a metabolically optimized and functionally persistent effector state.

### AS-3 engages mTORC1 signalling in CD8⁺ T cells

To determine whether the functional and metabolic effects of AS-3 were associated with direct engagement of the mTOR signalling pathway, we examined key nodes within the pathway in primary CD8^+^ T cells. Human CD8⁺

T cells isolated from PBMC were activated with AS-3 (10 μM), followed by immunoblot analysis of key mTOR pathway components. Immunoblot analysis showed that treatment with AS-3 increased phosphorylation of mTORC1, S6, and 4E-BP1 relative to control, indicating activation of the canonical mTORC1 downstream signalling axis. In contrast, phosphorylation of Akt was minimally affected under the same conditions (Figure 3A). A dose-response analysis of S6 phosphorylation further demonstrated a low EC_50_ of 0.095 μM, supporting the high potency of AS-3 in activating this pathway. These findings provide an important mechanistic anchor for the earlier observations and are consistent with the transcriptional signatures. Phosphorylation of S6 and 4E-BP1 is a defining output of mTORC1 activation and reflects enhanced translational capacity, biosynthetic readiness, and metabolic engagement. These signalling changes are therefore fully consistent with the previously observed rise in IFNγ and TNFα production, as effector cytokine output depends not only on transcriptional activation but also on metabolic support (37, 38). The limited effect on Akt suggests that AS-3 does not broadly amplify the upstream PI3K-Akt pathway, but instead acts in a manner that preferentially strengthens signalling at, or proximal to, the mTORC1 branch.

**Figure 3.**
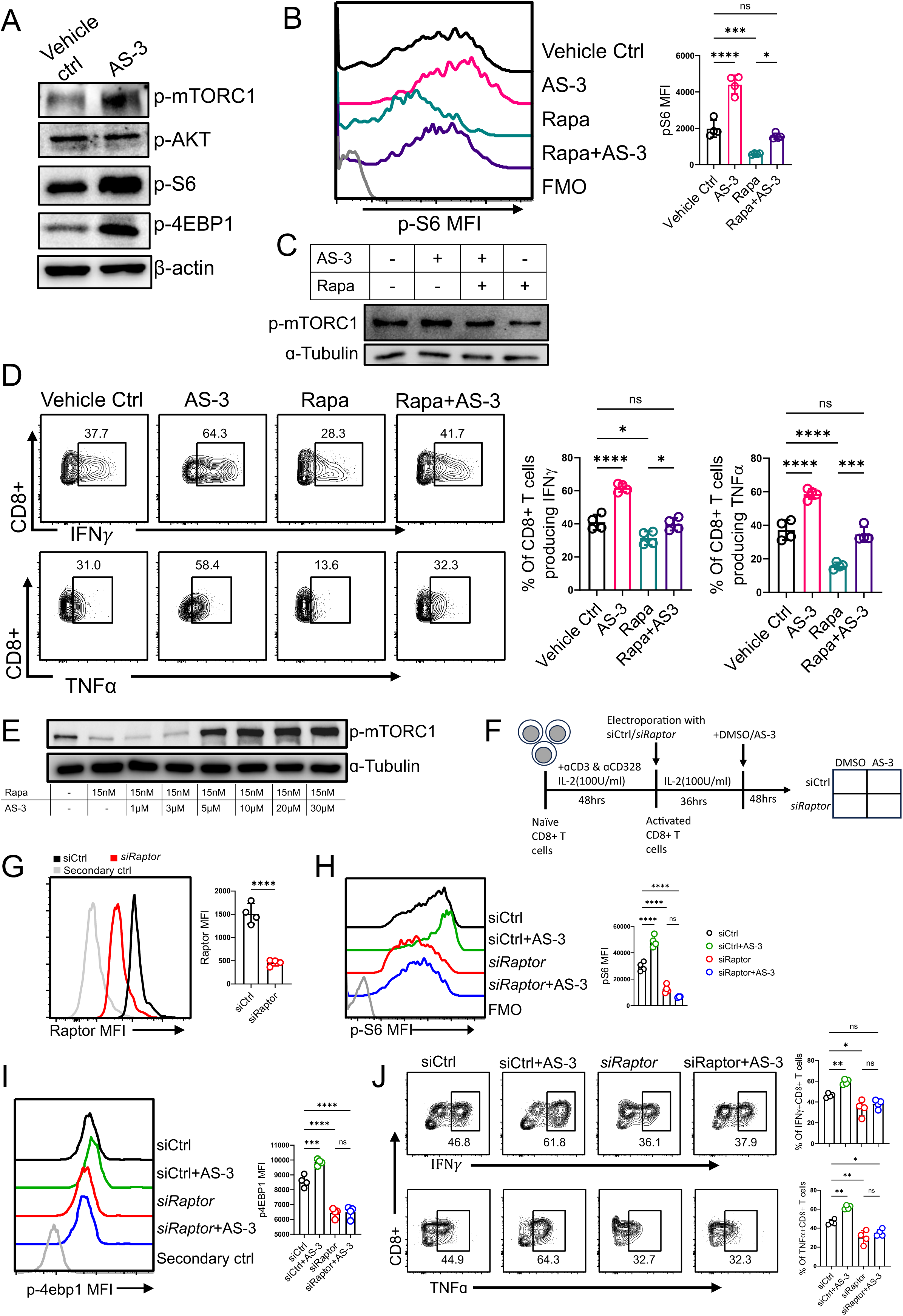
AS-3 enhances CD8⁺ T cell function through activation of the mTORC1 signaling pathway. (A) Immunoblot analysis of phosphorylated mTORC1, AKT, S6, and 4EBP1 in activated primary human CD8⁺ T cells treated with vehicle control or AS-3. (B) Representative flow cytometry histograms and quantification of phosphorylated S6 (pS6) mean fluorescence intensity (MFI) in activated CD8⁺ T cells treated with vehicle control, AS-3, rapamycin (15 nM), or rapamycin plus AS-3. Adjacent bar plot represents cumulative data from four independent experiments. (C) Immunoblot analysis of phosphorylated mTORC1 (Ser2448) in activated CD8⁺ T cells treated with AS-3 and/or rapamycin (15 nM). (D) Representative flow cytometry plots and quantification of IFNγ and TNFα producing CD8⁺ T cells following treatment with vehicle control, AS-3, rapamycin, or rapamycin plus AS-3. Numbers indicate the percentage of cytokine-positive cells within the CD8⁺ T-cell population. Adjacent bar plot represents cumulative data from four independent experiments. (E) Dose-response immunoblot analysis of phosphorylated mTORC1 (Ser2448) in activated CD8⁺ T cells treated with rapamycin (15 nM) together with increasing concentrations of AS-3 (1–30 μM). (F) Experimental design. Naïve CD8⁺ T cells from human PBMC were activated with anti-CD3 and anti-CD28 antibodies in the presence of IL-2 for 48 h, followed by electroporation with either control siRNA (siCtrl) or RAPTOR-targeting siRNA (*siRaptor*) using the Lonza 4D-Nucleofector system. After 36 h, cells were treated with vehicle control (DMSO) or AS-3 for 48 h prior to analysis by flow cytometry. (G) Representative histograms and quantification of RAPTOR expression showing knockdown efficiency. (H–I) Representative histograms and quantification of pS6 (H) and p4EBP1 (I) expression in the indicated groups. (J) Representative intracellular cytokine staining plots and quantification of IFNγ⁺ and TNFα⁺ CD8⁺ T cells. Adjacent bar plots (G-J) represents cumulative data from four independent biological replicates. Data are shown as mean ± SEM. ns, not significant; *p < 0.05, **p < 0.01, ***p < 0.001, ****p < 0.0001.

An important feature of AS-3 is its ability to preserve mTORC1 activity. This is mechanistically significant because mTORC1 signalling is central to the coupling of nutrient sensing, translational control, and metabolic fitness to T-cell effector function (39, 40). Activated CD8⁺ T cells were therefore treated in the presence of rapamycin (15 nM), a canonical mTORC1 inhibitor. Under these conditions, AS-3-treated cells retained higher levels of pS6 compared to rapamycin-treated controls, indicating a partial preservation of mTORC1 activity (Figure 3B). Notably, increased phosphorylation of mTOR was also observed in the presence of rapamycin, accompanied by elevated production of effector cytokines, suggesting that key functional outputs of mTORC1 signalling remain active despite inhibitory pressure (Figure 3C-D). Dose-response analysis further revealed a graded restoration of pathway activity, with phosphorylation levels increasing in a concentration-dependent manner and peaking at ∼5 µM, while remaining detectable even in the continued presence of rapamycin (Figure 3E). These observations extend the earlier finding that AS-3 activates canonical mTORC1 outputs under basal conditions, including phosphorylation of mTOR, S6, and 4E-BP1 with minimal modulation of Akt.

To genetically define the requirement for mTORC1 in mediating the activity of AS-3, Raptor, an obligate scaffolding component of the mTORC1 complex, was selectively inhibited in activated primary human CD8⁺ T cells using siRNA. CD8⁺ T cells isolated from human PBMCs were activated with anti-CD3/CD28 antibodies in the presence of IL-2, electroporated with either control siRNA (siCtrl) or RAPTOR-targeting siRNA (siRaptor), and subsequently treated with AS-3 (Figure 3F). Efficient Raptor silencing was confirmed by assessing intracellular Raptor levels, which were markedly reduced in the siRaptor group compared with the siCtrl group (Figure 3G). Importantly, *RAPTOR* knockdown substantially attenuated mTORC1 signalling, as evidenced by reduced phosphorylation of the canonical downstream effectors S6 and 4E-BP1 (Figure 3H and 3I). Interestingly, AS-3 treatment robustly enhanced mTORC1 signalling in siCtrl-transfected cells; however, this response was completely abrogated following *RAPTOR* knockdown, indicating that the ability of AS-3 to enhance downstream mTORC1 signalling is dependent on an intact Raptor-containing mTORC1 complex. Consistent with the loss of mTORC1 activity, Raptor knockdown significantly impaired the production of the effector cytokines IFNγ and TNFα, and AS-3 failed to restore cytokine expression under these conditions (Figure 3J). These findings demonstrate that the immunostimulatory activity of AS-3 requires an intact Raptor-containing mTORC1 complex. While the compound partially preserves mTORC1-associated signalling under pharmacological inhibition by rapamycin, disruption of the complex through Raptor depletion abolishes its ability to activate downstream signalling and promote CD8⁺ T-cell effector function.

Collectively, these findings support a model in which AS-3 stabilizes a mTORC1-centered immune-metabolic state, thereby strengthening its potential as an ex vivo conditioning strategy for improving adoptive T-cell therapies.

### AS-3 enhances antitumor efficacy of adoptively transferred CD8⁺ T cells

Given the observed enhancement of mTORC1 signaling, together with the upregulation of metabolic pathways associated with the generation and maintenance of durable T cell responses, we next investigated whether AS-3 treatment could translate into improved therapeutic efficacy in vivo. Because sustained mTOR activation has been implicated in tumor progression and metastasis, we first sought to exclude the possibility that residual AS-3 might be carried over with the adoptively transferred T cells and subsequently released within the tumor microenvironment, where it could potentially exert unintended effects on tumor cells. To address this concern, CD8+ T cells were exposed ex vivo to AS-3 (10 µM) for 72 h, followed by extensive washing, cell lysis, and extraction with cold methanol. LC-MS analysis failed to detect measurable levels of intracellular AS-3 in the treated T cells, indicating that the compound does not appreciably accumulate within T cells under the conditions used. These findings therefore suggest that adoptively transferred AS-3-treated T cells are unlikely to introduce biologically relevant amounts of residual AS-3 into recipient mice, minimizing the possibility of direct pharmacological activation of mTOR signaling in tumor cells following cell transfer.

Having established that AS-3 does not appreciably accumulate within T cells following ex vivo treatment, we next investigated whether AS-3 conditioning could enhance the therapeutic efficacy of adoptively transferred T cells in vivo. We employed an adoptive cell transfer (ACT) model using OT1 CD8⁺ T cells. OT1 T cells expressing a transgenic T cell receptor specific for the SIINFEKL peptide were activated ex vivo for 3 days in the presence of AS-3 or vehicle control and subsequently transferred into C57BL/6 mice with subcutaneously established EL4-OVA thymoma tumors (Figure 4A). Adoptive transfer of AS-3-treated OT1 T cells (0.5 × 10⁶ cells per mouse) resulted in significantly improved tumor control compared to mice receiving vehicle-treated cells (Figure 4B). This enhanced antitumor response was accompanied by a marked increase in overall survival of the tumor-bearing mice, demonstrating a clear therapeutic advantage conferred by AS-3 treatment. To investigate the underlying cellular basis, we quantified tumor-epitope reactive CD8⁺Vβ5.1⁺ OT1 T cells in tumor-bearing hosts. Mice receiving AS-3-treated T cells exhibited a higher frequency of circulating and tumor-infiltrating CD8⁺Vβ5.1⁺ OT1 T cells, indicating improved persistence in vivo (Figure 4C).

**Figure 4.**
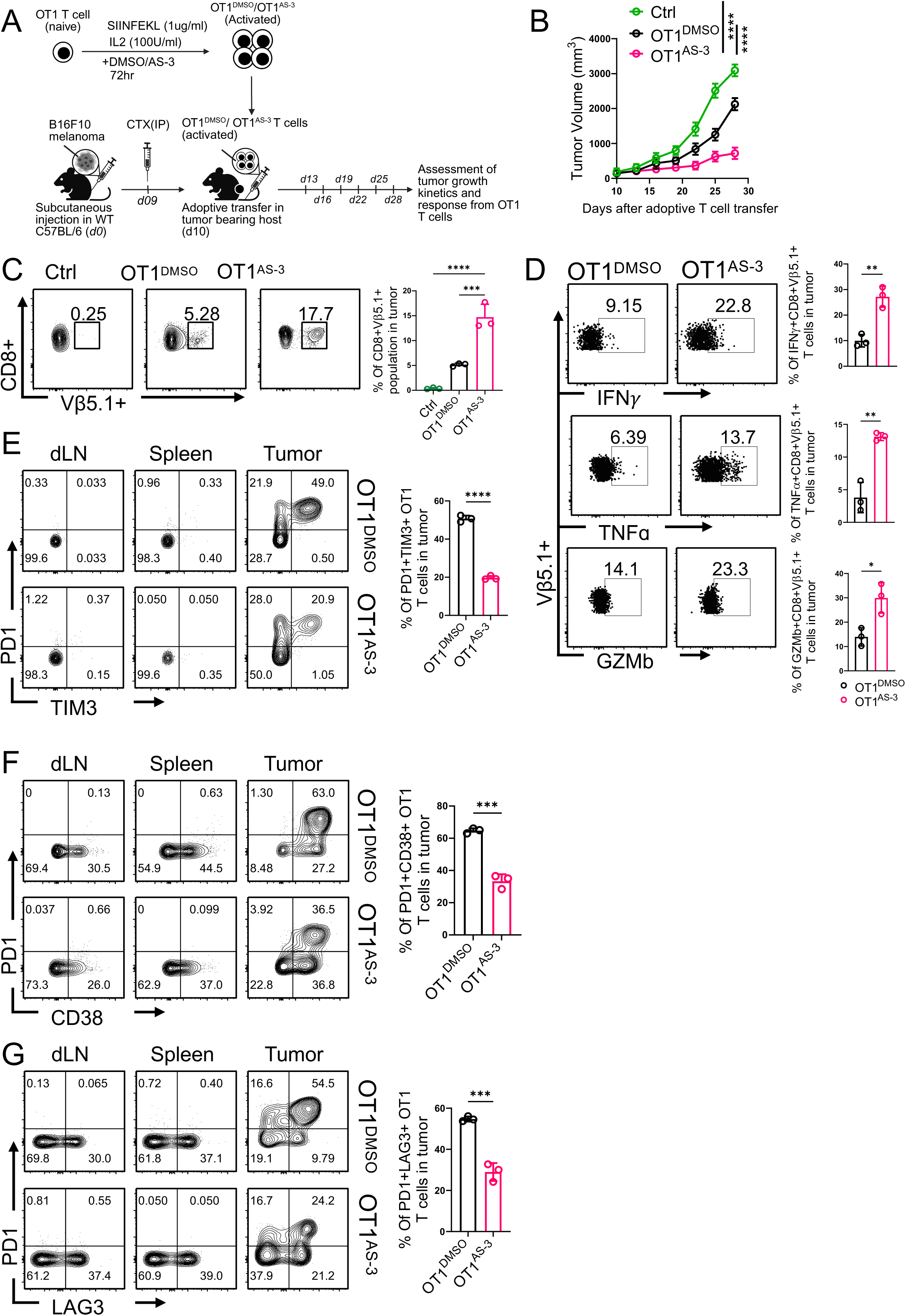
AS-3 preconditioning improves tumor control and limits exhaustion of adoptively transferred OT-1 T cells. (A) Experimental design illustrating ex vivo activation of OT-1 CD8+ T cells in the presence of vehicle control (OT1^DMSO^) or AS-3 (OT1^AS-3^) for 72h prior to adoptive transfer into C57BL/6 mice bearing established EL4-OVA tumors at day 7 post-inoculation. (B) Tumor growth kinetics measured by tumor volume over time (days post-adoptive transfer) in non-transferred control (Ctrl), OT1^DMSO^ and OT1^AS-3^ treatment cohorts. Data in figure demonstrate the mean tumor volume at each time point. (C) Flow cytometric evaluation of EL4-OVA–specific OT-1 T cells (vβ5.1^+^CD8^+^ T cells) at the tumor site after 17 days following T-cell transfer. Adjacent bar diagram representing the cumulative data of the percentage of tumor specific OT-1 T cells (*n* = 03 mice/group). (D) Representative flow cytometry plots and percentage quantification of tumor-infiltrating OT-1 T cells producing effector cytokines: IFNγ⁺, TNFα⁺ and GZMb. (E-G) Representative flow cytometry plots assessing T cell exhaustion via co-expression of (E) PD1 and TIM3, (F) PD1 and CD38, and (G) PD1 and LAG3 on OT1 T cells across the dLN, spleen, and tumor. Adjacent bar graphs quantify these respective double-positive populations within the tumor. Data are shown as mean ± SEM. ns, not significant; *p < 0.05, **p < 0.01, ***p < 0.001, ****p < 0.0001.

The differentiation of tumor-reactive T cells to terminally exhausted populations, characterized by sustained expression of inhibitory receptors and a hierarchical loss of effector cytokines, poses a major challenge for antitumor immunity (41, 42). Functionally, tumor-infiltrating OT1 T cells from the AS-3 group displayed significantly enhanced production of effector cytokines, including IFN-γ, TNF-α, and granzyme B, consistent with a sustained cytotoxic phenotype (Figure 4D). Importantly, OT-1 T cells ex vivo activated with AS-3 exhibited a marked reduction in the canonical markers of terminally exhausted T cells, including a reduction in the frequency of PD1^+^TIM3^+^, PD1^+^LAG3^+^, and PD1^+^CD38^+^ T cells, when derived from the tumor site compared to OT-1 T cells ex vivo activated with vehicle control. These data reveal that AS-3 enhances the persistence, functional capacity, and antitumor efficacy of adoptively transferred CD8⁺ T cells while limiting the development of terminal exhaustion.

### AS-3 promotes durable CD8⁺ T cell memory and sustained antitumor function in vivo

To determine whether AS-3 influences long-term T cell persistence and memory differentiation, we performed a “T cell parking” experiment in Rag1⁻/⁻ mice using Pmel CD8⁺ T cells specific for the gp100 antigen expressed by B16F10 melanoma. Pmel T cells were treated ex vivo with either vehicle control (DMSO) or AS-3 and activated for three days with their cognate peptide before adoptive transfer and “parking” into lymphopenic Rag1⁻/⁻ hosts in the absence of tumor engraftment (Figure 5A). This system enabled the assessment of homeostatic proliferation and memory formation independent of antigen-driven exhaustion. Analysis of peripheral blood 21 days post-transfer revealed that AS-3-treated Pmel T cells persisted at significantly higher frequencies compared with control cells (Figure 5B). Phenotypic characterization further demonstrated an increased proportion of effector memory (CD44⁺CD62L⁻) subsets in the AS-3 group (Figure 5C). To evaluate recall capacity, the same recipient mice were subsequently challenged with subcutaneous B16F10 melanoma implantation. Remarkably, within five days post-challenge, AS-3-treated Pmel cells underwent robust clonal expansion in response to antigen re-encounter, significantly exceeding the proliferative capacity of control-treated cells. Furthermore, these cells transitioned more efficiently into effector memory (CD44⁺CD62L⁻) subsets, suggesting superior preparedness to mount secondary immune responses (Figure 5D-E).

**Figure 5.**
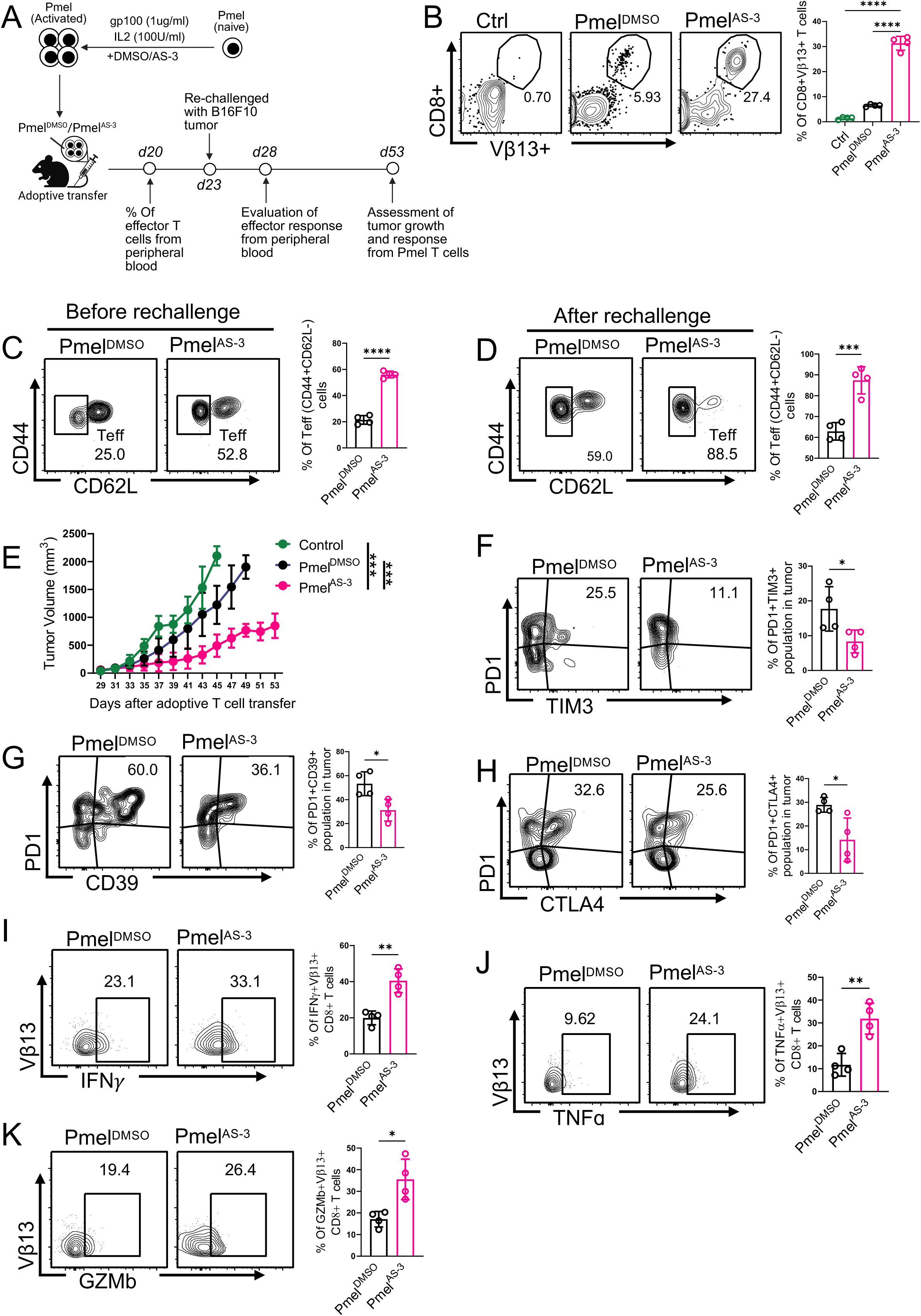
AS-3 preconditioning promotes durable antitumor immunity and persistence of Pmel CD8⁺ T cells. (**A**) Experimental schematic detailing ex-vivo Pmel CD8+ T cell priming with gp100, IL-2, ± AS-3 and adoptive transfer in RAG1-/-mice. Timeline highlights key experimental checkpoints: pre-rechallenge peripheral blood T cell evaluation (d20), B16F10 melanoma rechallenge (d23), post-rechallenge recall analysis (d 28), and assessment of tumor growth kinetics. (B) Flow cytometric analysis of gp100-specific Pmel T cells (vβ13⁺CD8⁺) in peripheral blood after tumor rechallenge following adoptive T-cell transfer. Adjacent bar plot representing the cumulative data of the percentage of vβ13⁺CD8⁺ T cells from respective group (*n* = 04/group). (C-D) Flow cytometry plots depicting circulating effector T cell (T_eff_, CD44^+^CD62L^−^) percentage (C) before rechallenge (day 20) and (E) after rechallenge (day 28) with tumor. Adjacent bar graph represent cumulative percentage of T_eff_ cells from respective groups. (E) Growth kinetics of rechallenged B16F10 tumor in Rag1⁻/⁻ mice receiving vehicle (Pmel^DMSO^) or AS-3 (Pmel^AS-3^) treated Pmel T cells. (F-H) Analysis and quantification of PD1⁺TIM3⁺ (F), PD1⁺CD39⁺ (G), and PD1⁺CTLA4⁺ (H) populations among intratumoral CD8^+^Vβ13⁺ Pmel T cells. (I-K) Quantification of intracellular cytokine production IFN-γ (I), TNF-α (J), and GZMb (K) from intratumoral Pmel T cells following ex vivo restimulation. Adjacent bar plots represent the cumulative data of the percentage of vβ13⁺CD8⁺ T cells positive for individual cytokines. Data are shown as mean ± SEM. ns, not significant; *p < 0.05, **p < 0.01, ***p < 0.001, ****p < 0.0001.

Consistent with these findings, tumor growth monitoring demonstrated that Rag1⁻/⁻ mice receiving AS-3-treated Pmel T cells mounted a more potent antitumor response than mice receiving vehicle-treated Pmel T cells (Figure 5F). Analysis of intratumoral Vβ13⁺ T cells, representing the transgenic TCR expressed by Pmel cells, revealed that AS-3-treated Pmel T cells displayed reduced features of terminal exhaustion, including lower frequencies of PD1⁺Tim3⁺, PD1⁺CD39⁺, and PD1⁺CTLA4⁺ populations, compared with vehicle-treated controls (Figure 5G-I). Importantly, AS-3-treated intratumoral Pmel T cells exhibited superior effector function, producing higher levels of IFN-γ, TNF-α, and granzyme B upon in vitro re-stimulation (Figure 5J-L). This enhanced cytokine and cytolytic molecule production reflects both preserved immune competence and heightened cytotoxic potential.

These findings demonstrate that transient ex vivo treatment of CD8⁺ T cells with AS-3 establishes a durable memory-associated program that promotes long-term persistence, robust recall expansion, and sustained effector functionality upon tumor antigen re-exposure. Importantly, AS-3 enables transferred T cells to resist terminal exhaustion while maintaining potent antitumor activity within the immunosuppressive tumor microenvironment, highlighting its therapeutic potential for improving adoptive T cell immunotherapy.

## Discussion

The therapeutic durability of ACT, including both TCR-T and CAR-T cell therapies, critically hinges on the sustained persistence and functional competence of transferred T cells in the tumor-bearing host (3, 6, 43). Increasing evidence suggests that prolonged T-cell survival and effector functionality are tightly linked to their metabolic programming, which orchestrates the epigenetic and transcriptional landscapes required for sustained anti-tumor activity (44). Consequently, metabolic reprogramming of T cells has emerged as a promising strategy to generate functionally superior ACT products (20, 21, 45). The present study identifies AS-3 as a potent small-molecule modulator that enhances the functional and metabolic fitness of CD8⁺ T cells through a RAPTOR-dependent mTORC1 signalling program. Identified through phenotypic screening and medicinal chemistry optimization, AS-3 enhanced effector function, mitochondrial fitness, and proliferative capacity during ex vivo T-cell activation. Importantly, transient ex vivo exposure of T cells to AS-3 was sufficient to generate durable anti-tumor responses in multiple ACT models, highlighting that metabolic conditioning during the manufacturing phase can profoundly influence subsequent T-cell fate and therapeutic efficacy in vivo.

A major obstacle limiting the long-term success of TCR-T and CAR-T therapy is the progressive acquisition of T cell dysfunction and terminal exhaustion within the tumor microenvironment (TME) (3, 43). Although chronic antigen stimulation and suppressive extrinsic signals are well-established contributors to this process, accumulating evidence indicates that exhaustion is also reinforced by intrinsic metabolic insufficiency and impaired bioenergetic adaptability (11). In this context, strategies capable of simultaneously enhancing metabolic resilience and preserving effector competence are particularly attractive. Our findings demonstrate that AS-3 induces a coordinated metabolic program characterized by enhanced glycolysis together with increased mitochondrial respiration and spare respiratory capacity. This dual metabolic reinforcement is especially significant because efficient glycolytic flux supports immediate effector activity, whereas mitochondrial fitness and respiratory reserve are essential for persistence, memory formation, and sustained anti-tumor immunity (22, 23, 31). It is noteworthy that although mTORC1 activity has been reported to inversely regulate T-cell memory differentiation, earlier studies from our group and others have demonstrated that mTORC1-dependent glycolysis in CD8⁺ T cells promotes the generation of effector memory cells in the context of both viral infection and cancer (20, 46, 47). Consistent with these observations, we argue that AS-3-mediated enhancement of glycolysis during ex vivo activation of CD8⁺ T cells may channel pyruvate toward acetyl-CoA generation, thereby sustaining mitochondrial fitness and facilitating histone acetylation-driven epigenetic remodelling (20). Such metabolic-epigenetic coupling may promote the metabolic adaptability of AS-3-treated T cells within the TME and facilitate their durable anti-tumor response.

The transcriptional remodelling induced by AS-3 further supports the establishment of a metabolically optimized and functionally persistent T cell state. RNA-sequencing analysis revealed enrichment of mTORC1 signalling, oxidative phosphorylation, MYC-driven anabolic pathways, and cell-cycle progression programs, accompanied by suppression of exhaustion-associated transcriptional modules. Notably, AS-3-treated T cells simultaneously expressed genes associated with cytotoxicity and effector function together with markers linked to memory-like persistence and self-renewal. This balanced transcriptional state is particularly important because excessive terminal differentiation often compromises durability of ACT products despite strong initial effector function (3, 48). In contrast, AS-3 appears to promote a hybrid state in which activated CD8⁺ T cells maintain robust cytotoxic potential while retaining metabolic flexibility and persistence-associated features.

Mechanistically, our data identify mTORC1 signalling as a central pathway engaged by AS-3. Activation of mTORC1 downstream effectors, including S6 and 4E-BP1, together with the minimal effect on Akt phosphorylation, suggests that AS-3 preferentially reinforces signalling at or proximal to the mTORC1 branch rather than broadly amplifying upstream PI3K-Akt signalling (49). This distinction may have important therapeutic implications. While sustained hyperactivation of PI3K-Akt pathways can drive terminal differentiation and exhaustion, controlled enhancement of mTORC1 activity may instead optimize biosynthetic and metabolic programs necessary for productive T-cell responses (12, 40, 50). Consistent with this notion, AS-3-treated T cells retained elevated mTORC1 activity even under rapamycin-mediated inhibitory conditions, suggesting that AS-3 stabilizes anabolic signalling programs required for maintaining T-cell functionality under metabolic stress.

The RAPTOR knockdown study provides direct mechanistic evidence that AS-3 exerts its immunostimulatory activity predominantly through the mTORC1 signalling axis. Although previous biochemical and transcriptomic analyses indicated robust activation of mTORC1-associated programs, these observations remained largely correlative. Genetic disruption of RAPTOR, an obligate scaffolding component of mTORC1, enabled us to directly interrogate pathway dependency. Loss of RAPTOR markedly reduced AS-3-induced phosphorylation of S6 and 4EBP1, two established downstream effectors of mTORC1 that regulate protein translation and cellular biosynthesis. These findings strongly suggest that AS-3 requires an intact RAPTOR-mTORC1 complex to propagate downstream signalling. Importantly, the attenuation of IFNγ and TNFα production following RAPTOR depletion demonstrates that enhanced effector differentiation induced by AS-3 is mechanistically linked to mTORC1 activation rather than arising from nonspecific T-cell stimulation. These findings integrate seamlessly with the transcriptomic data, which revealed significant enrichment of HALLMARK_MTORC1_SIGNALING, MYC targets, oxidative phosphorylation, glycolysis, and proliferative gene signatures in AS-3-treated T cells. The observed improvements in mitochondrial fitness and metabolic adaptability, and the RAPTOR knockdown studies establish mTORC1 activation as a critical determinant of the effector program induced by AS-3. These findings indicate that AS-3 promotes coordinated immunometabolic reprogramming through the RAPTOR-mTORC1 axis, thereby providing a compelling rationale for assessing its ability to enhance antitumor efficacy in adoptive cell therapy models in vivo.

The in vivo findings further establish the translational potential of AS-3 as an ex vivo conditioning strategy for ACT. OT1 T cells activated in the presence of AS-3 displayed superior tumor control, enhanced persistence, and reduced acquisition of terminal exhaustion markers following transfer into tumor-bearing hosts. Importantly, these effects were not restricted to acute effector responses. In the Pmel “T-cell parking” model, transient exposure to AS-3 promoted long-term persistence, improved recall expansion upon antigen re-encounter, and enhanced generation of effector-memory populations. These observations are particularly relevant because durable clinical responses following ACT are strongly associated with the persistence and recall capacity of transferred T cells. The ability of AS-3 to sustain both immediate effector function and long-term memory-like fitness therefore represents a highly desirable therapeutic attribute.

Several studies have attempted to improve ACT efficacy through pharmacologic modulation of T-cell signalling and metabolism during ex vivo expansion, including targeting PI3K, Wnt/β-catenin, BTK, and mitochondrial pathways (14, 15, 19). While many of these approaches promote memory-like differentiation, they often do so at the expense of immediate effector function or proliferative expansion. In contrast, AS-3 appears to preserve a favorable balance between activation, metabolic fitness, and persistence. The simultaneous enhancement of glycolytic activity, mitochondrial respiration, cytokine production, proliferation, and memory-associated programs suggests that AS-3 induces a broadly optimized immune-metabolic state rather than selectively enforcing a single differentiation trajectory.

Despite these encouraging findings, several important questions remain. Although our data strongly support engagement of mTORC1 signalling, the direct molecular target of AS-3 has not yet been identified. Future biochemical and structural studies will therefore be required to define the precise mechanism through which AS-3 modulates mTORC1 activity. In addition, while the present study demonstrates efficacy in murine ACT models and primary human CD8⁺ T cells, further validation using human CAR-T platforms will be important for translational development.

In summary, this study identifies AS-3 as a first-in-class small-molecule immune-metabolic modulator that enhances the therapeutic fitness of CD8⁺ T cells through reinforcement of mTORC1-driven metabolic and functional programs. By simultaneously promoting effector differentiation, mitochondrial fitness, persistence, and resistance to terminal exhaustion, AS-3 substantially improves the efficacy of adoptive T-cell therapy in vivo. . These findings establish transient ex vivo metabolic conditioning with AS-3 as a promising and clinically translatable strategy for generating superior ACT products with enhanced durability and anti-tumor function.

## Materials and methods

### Cell lines

EL4-OVA thymoma and B16F10 melanoma cells were maintained under standard culture conditions and routinely tested to confirm the absence of mycoplasma contamination. Cells were passaged a minimum of four times in complete culture medium prior to in-vivo tumor implantation experiments.

### Isolation and activation of human CD8⁺ T cells

Peripheral blood mononuclear cells (PBMCs) were isolated from de-identified healthy donor buffy coats following institutional ethical approval. PBMCs were separated by Ficoll density-gradient centrifugation, and CD8⁺ T cells were purified by negative magnetic selection. Purified CD8⁺ T cells were cultured in RPMI-1640 supplemented with 10% fetal bovine serum (FBS), 100 U/mL penicillin, 100 μg/mL streptomycin, and recombinant human IL-2 (100 U/mL). Cells were activated using plate-bound anti-CD3 (5 μg/mL) and anti-CD28 (2 μg/mL) antibodies in the presence of vehicle control (DMSO), AS-1, or AS-3 at the indicated concentrations for 72 h. For mTORC1 inhibition studies, activated CD8⁺ T cells were treated with rapamycin (15 nM) alone or in combination with AS-3. For dose-response experiments, increasing concentrations of AS-3 were added in the presence of rapamycin, followed by analysis of mTOR pathway activation and cytokine production.

### Cell proliferation and viability assays

Cell proliferation was assessed using CellTrace Violet (CTV; Thermo Fisher Scientific). Purified CD8⁺ T cells were labeled with 5 μM CTV for 20 min at 37°C prior to activation according to the manufacturer’s instructions. Labeled cells were subsequently cultured under the indicated experimental conditions, and proliferation was evaluated by flow cytometric analysis of dye dilution after 72 h. Cell viability was determined using LIVE/DEAD™ Fixable Yellow Dead Cell Stain (Thermo Fisher Scientific; 1:1000 dilution).

### Flow cytometry

Cells were stained with fluorochrome-conjugated antibodies in FACS buffer (PBS containing 0.5% BSA and 0.1% sodium azide) for 30 min at 4°C. Cell viability was assessed using LIVE/DEAD™ Fixable Yellow Dead Cell Stain (1:1000 dilution). For intracellular cytokine staining, cells were stimulated with PMA (100 ng/mL) and ionomycin (1 μg/mL) in the presence of GolgiPlug™ (1 μL/mL) for 4 h at 37°C, followed by surface staining, fixation/permeabilization using the Cytofix/Cytoperm kit (BD Biosciences), and intracellular staining for IFN-γ, TNF-α, Granzyme B. For glucose uptake analysis, activated CD8⁺ T cells were washed with PBS and incubated with 2-NBDG (100 μM) in glucose-free RPMI-1640 medium for 30 min at 37°C. For assessment of mitochondrial mass and membrane potential, cells were washed with PBS and stained with MitoTracker Green FM (100 nM) or MitoTracker Deep Red FM (100 nM), respectively, in HBSS for 30 min at 37°C. Following staining, cells were washed and analyzed by flow cytometry. For phospho-flow analysis, cells were fixed with BD Phosflow™ Fix Buffer I and permeabilized with BD Phosflow™ Perm Buffer III according to standard phosphoprotein staining procedures, followed by intracellular staining for phospho-S6. Samples were acquired on a BD LSRFortessa flow cytometer (BD Biosciences) and analyzed using FlowJo software.

### Extracellular flux analysis

Cellular metabolic activity was assessed using an Agilent Seahorse XFe24 Extracellular Flux Analyzer (Agilent Technologies). Activated human CD8⁺ T cells were harvested, washed, and resuspended in Seahorse XF Base Medium supplemented with 1 mM sodium pyruvate and adjusted to pH 7.4. Cells (3 × 10⁵ cells/well) were seeded onto Cell-Tak-coated XF24 cell culture microplates and centrifuged at 200 × g for 1 min to facilitate cell attachment. Plates were incubated for 30 min at 37°C in a non-CO₂ incubator before analysis. For mitochondrial stress tests, oxygen consumption rate (OCR) was measured under basal conditions and following sequential injections of oligomycin (10 μM), FCCP (10 μM), and rotenone/antimycin A (1 μM each). For glycolytic stress tests, extracellular acidification rate (ECAR) was measured following sequential injections of glucose (10 mM), oligomycin (10 μM), and 2-deoxy-D-glucose (2-DG; 100 mM). Measurements were performed using standard Seahorse assay cycles consisting of 3 min mixing, 2 min waiting, and 3 min measurement periods. Data were analyzed using Wave software (Agilent Technologies).

### Immunoblot analysis

Activated CD8⁺ T cells were lysed in RIPA buffer supplemented with protease and phosphatase inhibitors. Protein concentrations were determined using the Bradford assay. Equal amounts of protein were resolved by SDS-PAGE and transferred onto PVDF membranes. Membranes were blocked with 5% BSA and incubated overnight at 4°C with primary antibodies. After incubation with HRP-conjugated secondary antibodies, protein bands were visualized using enhanced chemiluminescence and quantified using ImageJ software.

### Electroporation-Mediated Gene Silencing in CD8+ T Cells

Activated human CD8+ T cells (48 hours post *α*CD3/*α*CD28 stimulation in 100 U/mL IL-2) were harvested, washed with PBS, and pelleted to remove residual culture medium. Cells were resuspended in 4D-Nucleofector Solution mixed with Supplement (Lonza) at a standard concentration according to the manufacturer’s protocol for primary T cells. Target siRNA against *Raptor* (*siRaptor*) or non-targeting control siRNA (*siCtrl*) was added to the cell suspension. Transfection was executed using the 4D-Nucleofector System (Lonza) with a provided primary cell-optimized pulse program. Immediately following electroporation, pre-warmed culture medium was added to the nucleofection cuvettes, and transfected cells were transferred into culture plates containing media supplemented with 100 U/mL IL-2. Transfected cells were incubated for 36 hours post-electroporation prior to downstream treatment with DMSO or AS-3 for 48 hours.

### RNA sequencing and bioinformatic analysis

Primary human CD8⁺ T cells activated in the presence of vehicle control or AS-3 were harvested for transcriptomic analysis. Total RNA was extracted using the RNeasy Mini Kit (Qiagen), and RNA integrity was confirmed prior to library preparation. Poly(A)-selected, strand-specific libraries were generated using the KAPA RNA HyperPrep workflow and sequenced on an Illumina NovaSeq platform to generate paired-end 150 bp reads. Raw sequencing reads were subjected to quality control and adapter trimming before alignment to the human reference genome (GRCh38) using STAR. Gene-level counts were generated using featureCounts and imported into R for downstream analyses. Differential expression analysis was performed using DESeq2. Genes with an adjusted P value < 0.05 were considered significantly differentially expressed. Principal component analysis (PCA), volcano plot, and heatmap were generated using R. Gene set enrichment analysis (GSEA) was performed using Hallmark and Reactome gene sets to identify pathways significantly enriched following AS-3 treatment.

### OT1 adoptive cell transfer model

Splenic OT1 CD8⁺ T cells were isolated from OT1 transgenic mice and activated ex vivo with SIINFEKL peptide (1ug/ml) in the presence of vehicle control (DMSO) or AS-3 for 72 h. C57BL/6 mice bearing established subcutaneous EL4-OVA tumors received 1 × 10⁶ activated OT1 cells by intravenous injection. Tumor growth was monitored longitudinally using caliper measurements. At experimental endpoints, peripheral blood, spleens, draining lymph nodes, and tumors were harvested for evaluation of transferred Vβ5.1⁺ OT1 T cells, cytokine production, and exhaustion-associated markers.

### T-cell parking and recall response model

Pmel CD8⁺ T cells were activated ex vivo with gp100 peptide (1ug/ml) in the presence of vehicle control (DMSO) or AS-3 for 72 h and intravenously transferred into Rag1−/− recipient mice(1.5 × 10⁶ activated Pmel CD8⁺ T cells per mouse). Twenty-one days after transfer, peripheral blood was collected to assess T-cell persistence and effector phenotype. Recipient mice were then challenged with subcutaneous B16F10 melanoma cells. Recall expansion, effector differentiation, exhaustion-associated phenotypes, cytokine production, and antitumor responses were evaluated by flow cytometry and tumor growth monitoring.

### Statistical analysis

Data are presented as mean ± SEM unless otherwise indicated. Statistical analyses were performed using GraphPad Prism and R software. Comparisons between two groups were performed using unpaired two-tailed Student’s t-test, whereas multiple-group analyses were conducted using one-way or two-way ANOVA with appropriate post hoc correction. Differences were considered statistically significant at P < 0.05.

## Author Contribution

A.T. and S.C. supervised and coordinated different aspects of the project and oversaw the progress of each component. A.T. and S.C. acquired the funding for the project from CSIR-FBR070304 and CSIR National Laboratory Scheme ULIP (MLP2620). D.S., P.G., and S.G. contributed equally to the study as they made indispensable but complementary contributions. A.T., S.C., D.S., and S.G. conceptualised the idea of identifying mTOR pathway modulators through screening and subsequent application. P.G. mainly took the lead and was assisted by D.S, H.S.S. and Deborpita S. in standardizing the in vitro screening platforms and validation of mTOR pathway activator. P.G and S.C. analyzed the in vitro validation data. D.S and S.G standardized the synthetic schemes and performed the synthesis/characterization of new compounds. P.G performed the in vivo animal model efficacy experiments. D.S. and P.G thank UGC for the fellowship. A.T, D.S and S.C. wrote the manuscript. The manuscript was finally corrected by all the authors.

