## Supplementary material for "Discovery of a Small-Molecule mTORC1 Pathway Activator that Enhances Adoptive T-Cell Therapy via Immuno-metabolic Reprogramming of CD8^+^ T Cells": Sarkar et al_Supporting Information.pdf

4.5.15 Synthesis of N-(8-(4-methoxyphenyl)-9-methyl-9H-purin-6-yl)-4-methylbenzenesulfonamide (AS3) 16
4.5.16 Synthesis of N-(9-isopropyl-8-(4-methoxyphenyl)-9H-purin-6-yl)-4-methylbenzenesulfonamide (AS4) 16
4.5.17 Synthesis of N-(9-cyclopentyl-8-(4-methoxyphenyl)-9H-purin-6-yl)-4-methylbenzenesulfonamide (AS5) ......................................................................................... 16
4.5.18 Synthesis of 4-methoxy-N-(8-(4-methoxyphenyl)-9-methyl-9H-purin-6-yl)benzenesulfonamide (AS6) 17
4.5.19 Synthesis of N-(8-(4-methoxyphenyl)-9-methyl-9H-purin-6-yl)-4-(trifluoromethyl)benzene sulfonamide (AS7) ......................................................................................................... 17
4.5.20 Synthesis of N-(8-(4-methoxyphenyl)-9-methyl-9H-purin-6-yl)cyclohexanesulfonamide (AS8) 18
4.5.21 Synthesis of N-(8-(2,4-dimethoxyphenyl)-9-methyl-9H-purin-6-yl)-4-methylbenzenesulfonamide (AS9) 18

4.5.22 Synthesis of N-(8-(4-cyanophenyl)-9-methyl-9H-purin-6-yl)-4-methylbenzenesulfonamide (AS10) 18

4.5.23 Synthesis of N-(8-(4-chlorophenyl)-9-methyl-9H-purin-6-yl)-4-methylbenzenesulfonamide (AS11) 19

5. NMR SPECTROSCOPIC DATA 20

5.1 $^{1}H$ NMR of compound 2a (400 MHz, DMSO-$d_6$): 20

5.2 $^{13}C$ NMR of compound 2a (400 MHz, DMSO-$d_6$): 20

5.3 $^{1}H$ NMR of compound 2b (400 MHz, DMSO-$d_6$): 21

5.4 $^{13}C$ NMR of compound 2b (400 MHz, DMSO-$d_6$): 21

5.5 $^{1}H$ NMR of compound 2c (400 MHz, DMSO-$d_6$): 22

5.6 $^{13}C$ NMR of compound 2c (101 MHz, DMSO-$d_6$): 22

5.7 $^{1}H$ NMR of compound 3a (600 MHz, DMSO-$d_6$): 23

5.8 $^{13}C$ NMR of compound 3a (101 MHz, DMSO-$d_6$): 23

5.9 $^{1}H$ NMR of compound 3b (400 MHz, DMSO-$d_6$): 24

5.10 $^{13}C$ NMR of compound 3b (101 MHz, DMSO-$d_6$): 24

5.11 $^{1}H$ NMR of compound 3c (600 MHz, Chloroform-*d*): 25

5.12 $^{13}C$ NMR of compound 3c (101 MHz, Chloroform-*d*): 25

5.13 $^{1}H$ NMR of compound 4a (400 MHz, DMSO-$d_6$): 26

5.14 $^{13}C$ NMR of compound 4a (101 MHz, DMSO-$d_6$): 26

5.15 $^{1}H$ NMR of compound 4b (400 MHz, Chloroform-*d*): 27

5.16 $^{13}C$ NMR of compound 4b (101 MHz, Chloroform-*d*): 27

5.17 $^{1}H$ NMR of compound 4c (400 MHz, Chloroform-*d*): 28

5.18 $^{13}C$ NMR of compound 4c (101 MHz, Chloroform-*d*): 28

5.19 $^{1}H$ NMR of compound 4d (400 MHz, Chloroform-*d*): 29

5.20 $^{13}C$ NMR of compound 4d (101 MHz, Chloroform-*d*): 29

5.21 $^{1}H$ NMR of compound 4f (400 MHz, DMSO-$d_6$): 30

5.22 $^{13}C$ NMR of compound 4f (101 MHz, DMSO-$d_6$): 30

5.23 $^{1}H$ NMR of compound AS1 (400 MHz, Chloroform-*d*): 31

5.24 $^{13}C$ NMR of compound AS1 (101 MHz, Chloroform-*d*): 31

5.25 $^{1}H$ NMR of compound AS2 (400 MHz, Chloroform-*d*): 32

5.26 $^{13}C$ NMR of compound AS2 (101 MHz, Chloroform-*d*): 32

5.27 $^{1}H$ NMR of compound AS3 (400 MHz, Chloroform-*d*): 33

5.28 $^{13}C$ NMR of compound AS3 (101 MHz, Chloroform-*d*): 33

5.29 $^{1}H$ NMR of compound AS4 (400 MHz, Chloroform-*d*): 34

5.30 $^{13}C$ NMR of compound AS4 (101 MHz, Chloroform-*d*): 34

5.31 $^{1}H$ NMR of compound AS5 (400 MHz, Chloroform-*d*): 35

### 1. Figure S1. Functional and metabolic effects of AS-3 in CD8$^+$ T cells

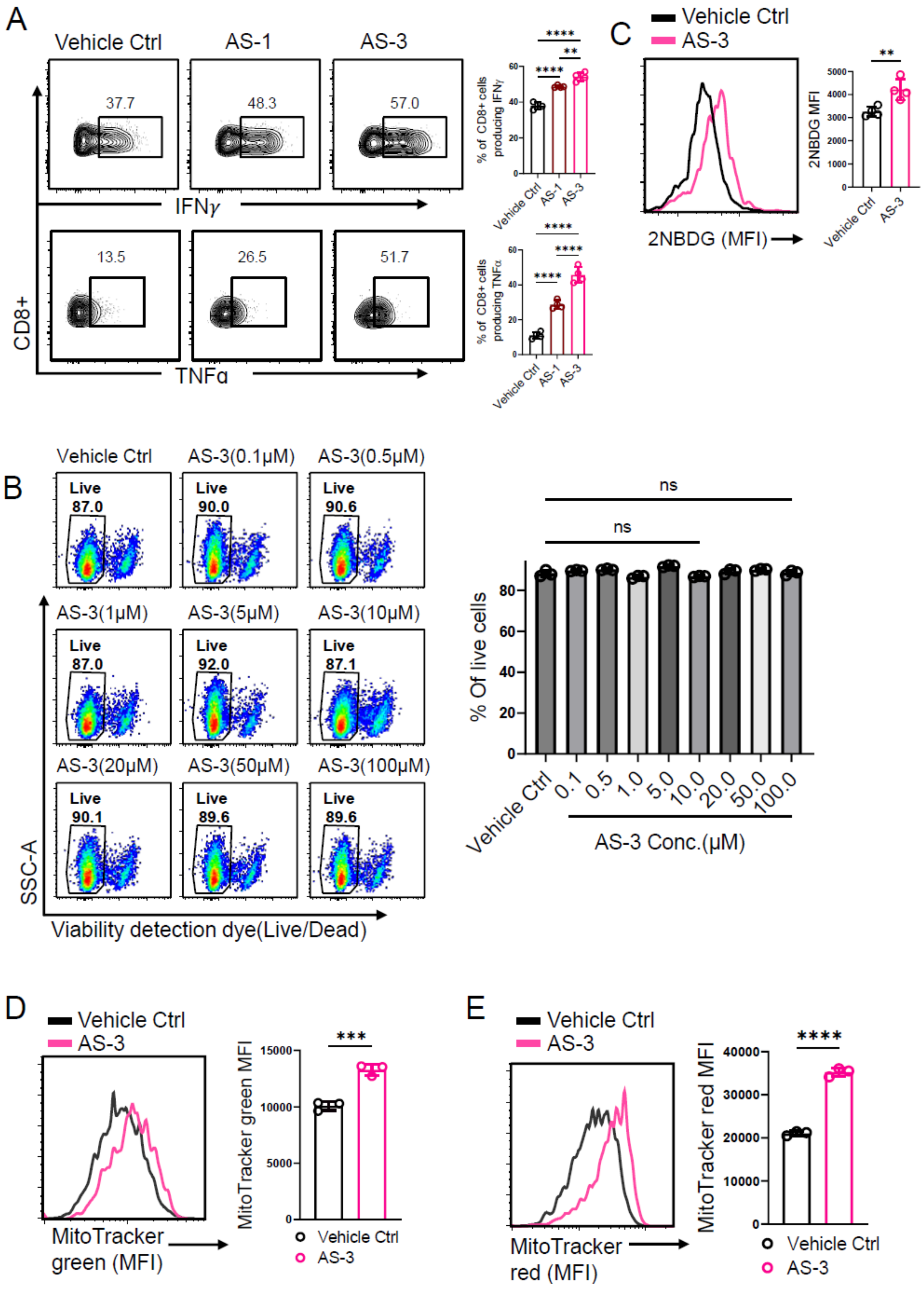

**Figure S1. AS-3 enhances CD8$^+$ T cell effector function and metabolic fitness without compromising cell viability.** (A) Flow cytometric analysis of intracellular IFN-γ and TNF-α production from human CD8$^+$ T cells following activation in the presence of vehicle control, AS-1, or AS-3. (B) Viability assessment of activated human CD8$^+$ T cells treated with increasing concentrations of AS-3, measured by Live/Dead staining and quantified as percentage of live cells. (D–F) Metabolic profiling of activated human CD8$^+$ T cells treated with vehicle control or AS-3. Representative histograms and corresponding quantification of 2-NBDG uptake (D), mitochondrial mass measured by MitoTracker Green fluorescence (E), and mitochondrial membrane potential measured by MitoTracker Deep Red fluorescence (F). Bar graphs depict cumulative data of mean fluorescence intensity (MFI) from three independent experiment. Data are shown as mean ± SEM. ns, not significant; *$p < 0.05$, **$p < 0.01$, ***$p < 0.001$, ****$p < 0.0001$.

#### 2. Figure S2. MASS SPECTROMETRIC EVALUATION OF CELLULAR UPTAKE OF AS-3

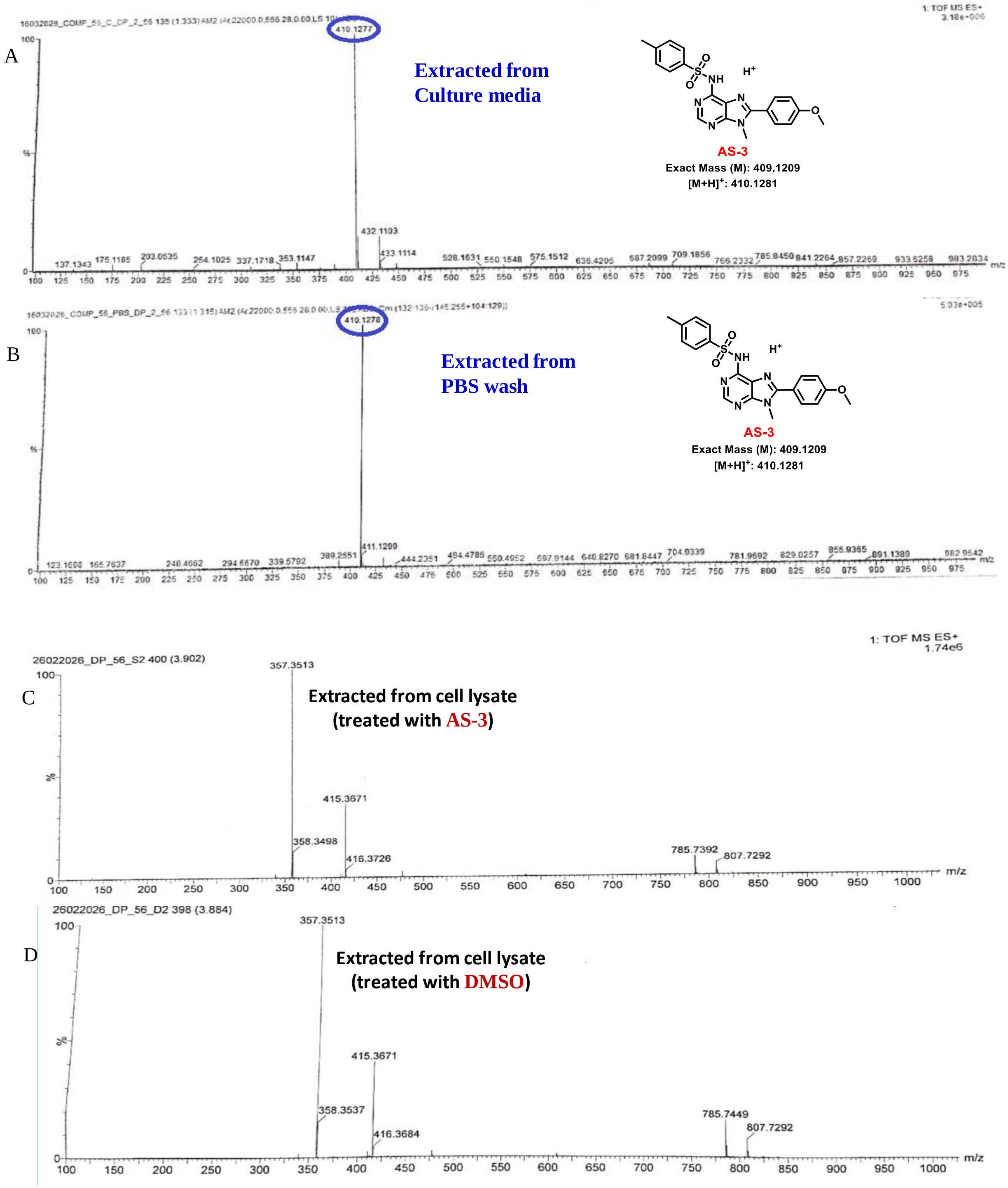

**Figure S2. HRMS analysis of the cellular uptake of AS-3.** Representative high-resolution mass spectra acquired using electrospray ionization (ESI-HRMS) of extracts obtained from (A) culture medium, (B) PBS wash, (C) cell lysate from AS-3 treated cells, and (D) cell lysate from DMSOtreated control cells. Following treatment with AS-3, extracellular medium and PBS wash fractions were collected and extracted, while cell lysates from AS-3 and DMSO treated cells were processed in parallel. The AS-3 treated and control cell lysates exhibited comparable mass spectral profiles, with no detectable ion corresponding to AS-3 in the treated cell lysate under the experimental conditions, suggesting negligible intracellular accumulation within the detection limit of the ESI-HRMS analysis.

### 3. CHEMISTRY

Our designed adenine core-based compounds were synthesized by following schemes which describes the synthetic protocols and strategies for the synthesis of required derivatives with modifications at C6-$NH_2$ position. The synthesis was initiated from commercially available adenine (**1**). N9 methylation of adenine was performed by reacting adenine (**1**) with methyl iodide (MeI) (1.5 eqv) in presence of THF as solvent and tetrabutylammonium fluoride (TBAF) 1M in THF (1 eqv) as base at room temperature for 12-16 hrs to obtain compound **2a**. Compound **2b-2c** was synthesized by reacting  Bromination of compound **2a-2c** was done by using N-Bromosuccinimide (1.5 eqv) in a dry $CHCl_3$ at room temperature (25-30 °C) for 12-16 hrs to obtain compound **3a-3c**. The C-C cross-coupling reactions (Suzuki-Miyaura coupling) on **3a-3c** was performed by palladium catalyst; $Pd(PPh_3)_4$ (0.1 eqv) in dioxane/$H_2O$ (9:1 solvent ratio v/v) at 120 °C for 12-16 hrs in presence of $K_2CO_3$ (2 eqv) and requisite phenylboronic acid (1.2 eqv) to obtain **4a-4f**. Next, **AS:1-10** were prepared by reacting compound **4a-4f** with required substituted benzoyl chloride (1.2 eqv) or phenyl isocyanate in dry THF as the solvent and LiHMDS (2 eqv) as a base at room temperature (25-30 °C) for 2 hrs while sulphonamides are prepared by reacting compound **4a-4f**  with different requisite substituted benzene sulfonyl chlorides (1.2 eqv) in dry DMF as the solvent and sodium hydride (NaH) (1.5 eqv) as a base at room temperature (25-30 °C) for 1 hr (Scheme).

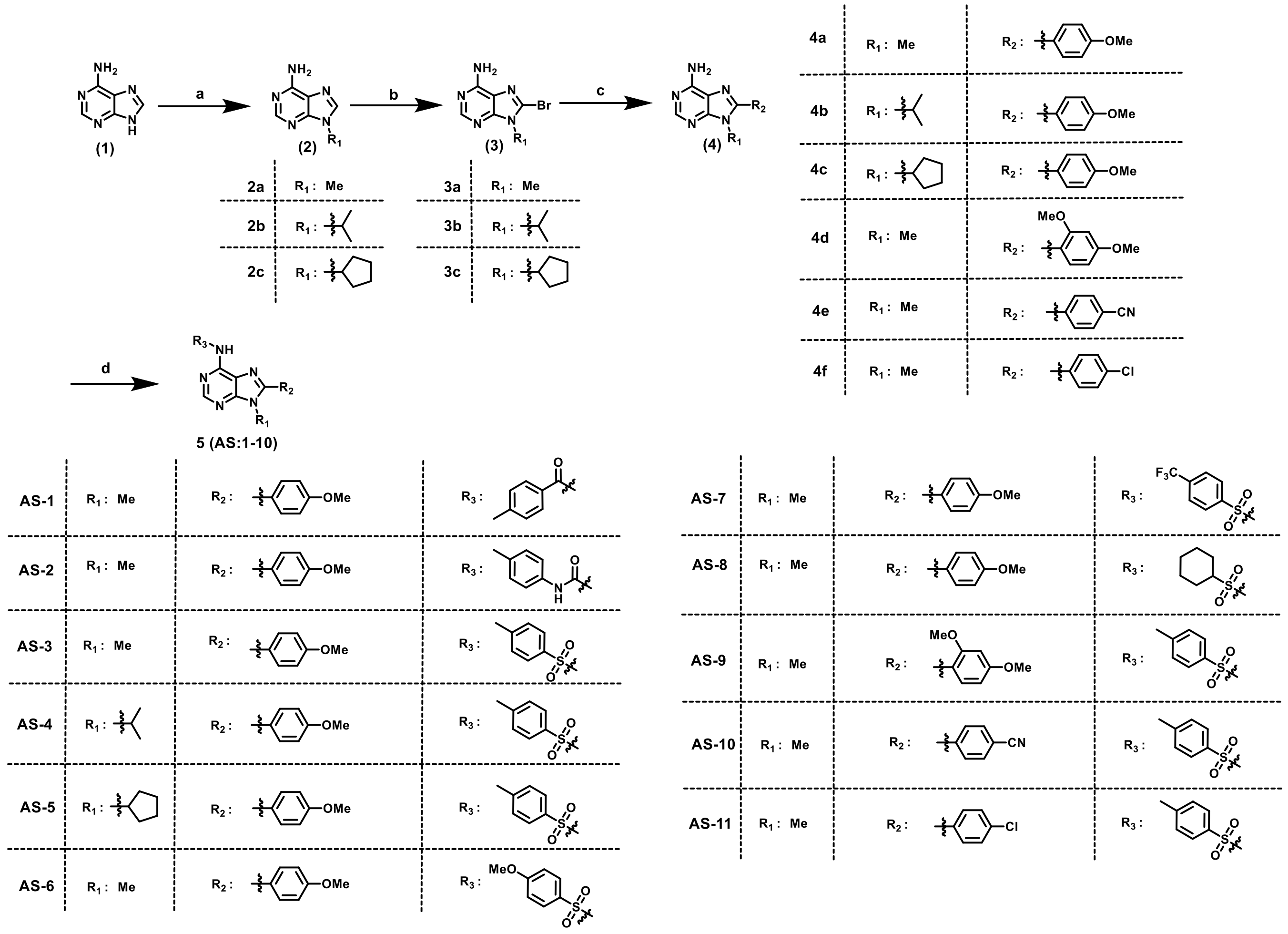

[a]**Reagents and conditions:** (a) $CH_3I$, TBAF, dry THF, room temperature, 12-16 hrs, 87% or $R_1$-I, $K_2CO_3$, dry DMF, room temperature, 12 hrs, (75-78%); (b) NBS, dry $CHCl_3$, 4-5 hrs, (85-95%); (c) $R_2$-$B(OH)_2$, $Pd(PPh_3)_4$, dioxane/ $H_2O$ (9:1), $K_2CO_3$, 120 °C, 12-18 hrs, (62-72%); (d) 4-methylbenzoyl chloride, LiHMDS, dry THF, room temperature, 2 hrs, (68%); 4-methylphenyl isocyanate, LiHMDS, dry THF, room temperature, 2 hrs, (72%); $R_3$-$SO_2Cl$, NaH, dry DMF, room temperature, 1 hr, (68-84%).

### 4. CHEMICAL SYNTHESIS AND METHODS

#### 4.1 General methods

All starting materials, reagents, and solvents were purchased from commercial suppliers and used without further purification. Dry solvents were either commercially purchased or dried using a standard protocol. Reactions that had a sensitivity toward moisture or oxygen were carried out under a dry nitrogen or argon atmosphere. All the TLC experiments were performed on silica gel plates (Merck silica gel 60, F254). The spots were visualized under UV light (λ = 254 and 365 nm) or by using the appropriate stain. Compounds were purified using a Teledyne ISCO Combi Flash $R_f$ system using a 230−400 mesh size silica gel. $^{1}H$ NMR was recorded at 300 MHz (Bruker-DPX), 400 MHz (JEOL), and 600 MHz (Bruker AVANCE) frequencies, and $^{13}C$ NMR spectra were recorded at 75 MHz (Bruker-DPX), 100 MHz (JEOL), and 150 MHz (Bruker AVANCE) frequencies in $CDCl_3$ or $CD_3OD$ or *DMSO-$d_6$* using tetramethylsilane as the internal standard. The following abbreviations were used to explain multiplicities: s = singlet, d = doublet, t = triplet, q = quartet, m = multiplet, brs. = broad singlet. The coupling constant, *J*, was reported in the Hertz unit (Hz). High-resolution mass spectroscopy, HRMS (m/z), was carried out using ESI (Q-Tof Micro mass spectrometer), and ESI (LTQ Orbitrap XL mass spectrometer).

#### 4.2 General Procedure A: Bromination

Compound **2a-2c** (1 eqv) was taken in a dry $CHCl_3$ and N-Bromosuccinimide (1.5 eqv) was added in reaction mixture and stirred at room temperature for 4-5 hours. Progress of the reaction was thoroughly monitored through TLC, and after completion of the reaction mixture was washed with satd. solution of $Na_2S_2O_3$ and extracted with EtOAc. The organic layer was washed with brine solution, dried over $Na_2SO_4$ and was evaporated under vacuum to obtain the crude. The product was then purified by column chromatography using 230−400 mesh size silica gel and EtOAC/Pet ether or MeOH/$CHCl_3$ as eluents to get the pure compounds with a yield of 85-95%.

#### 4.3 General Procedure B: Suzuki Coupling

Compound **3a-3c** (1 eqv), appropriate aromatic boronicacid (1.2 eqv), $K_2CO_3$ (2 eqv) were taken in a pressure tube and dissolved in (9:1) mixture of Dioxane & water. Solution was purged with Argon gas for 30 minutes. Then catalyst $Pd(PPh_3)_4$ (0.1 eqv) was added and the reaction mixture was stirred at 100˚C -110 ˚C for for 12-16 hours. The progress of the reaction was monitored by checking TLC. After reaction completion, the reaction mixture was filtered through a celite bed and washed with 10% MeOH in $CHCl_3$ (V/V); the organic layer was washed with brine solution and dried over $Na_2SO_4$. The solvent was evaporated under vacuum to obtain the crude, which was then purified in flash chromatography using 230-400 mesh size silica gel and EtOAc in Pet ether and MeOH in $CHCl_3$ as eluent to afford pure compound with 62-72% yield.

#### 4.4 General Procedure C: Sulphonamide formation reaction

Compound **4** (1 eqv) was dissolved in dry DMF and then NaH (2 eqv) was added portion wise at ice-cold condition keeping nitrogen atmosphere and the reaction mixture was allowed to stir at room temperature for 15 minutes. Then required sulfonyl chlorides (1.2 eqv) were added portion wise at ice-cold condition and then stirred at room temperature (25-30 °C) for another 1 hr. The progress of the reaction was monitored by checking TLC. After completion, reaction mass was washed with water and extracted with ethyl acetate. The organic layer was washed with brine solution, dried over $Na_2SO_4$ and was evaporated under vacuum to obtain the crude. If needed column chromatography was performed to get the pure product.

#### 4.5 Synthesis and characterization

##### 4.5.1 Synthesis of methyl 9-methyl-9H-purin-6-amine (2a):

Adenine **1** (5gm, 37.03 mmole) was taken in dry THF and TBAF (37.03mL, 37.03mmole) was added as a base at ice-cold condition. Then the reaction mixture was stirred in room temperature for 30 mins keeping nitrogen atmosphere. Next, MeI (1.8mL, 55.54 mmole) was added dropwise at ice-cold condition and the reaction mixture was allowed to stir at room temperature for 12-16 hours. Progress of the reaction was thoroughly monitored through TLC, and after completion of the reaction solvent was evaporated in reduced pressure and the product was purified by column chromatography using 230−400 mesh size silica gel and 5% MeOH/$CHCl_3$ as eluent to get compound **2a** as off white solid (yield 87%). $^1$H NMR (400 MHz, DMSO-$d_6$) δ in ppm 8.10 (s, 1H), 8.02 (s, 1H), 7.06 (s, 2H), 3.67(s, 3H). $^{13}$C NMR (101 MHz, DMSO-$d_6$) δ in ppm 156.40, 152.89, 150.41, 141.89, 119.19, 29.82. HRMS (ESI) *m*/*z* (M + H)$^+$ calculated for $C_6H_8N_5$: 150.0780; found: 150.0774.

##### 4.5.2 Synthesis of 9-isopropyl-9H-purin-6-amine (2b):

Adenine **1** (1.5 g, 11.1 mmol), $K_2CO_3$ (1.8 g, 13.3 mmol), and 2-iodopropane (1.1 mL, 11.1 mmol) were dissolved in 5 mL of dry DMF. The reaction mixture was stirred at room temperature for 12-16 hours. Progress of the reaction was thoroughly monitored through TLC. Upon completion, the reaction mixture was partitioned between EtOAc (50 ml) and ice-water (50 ml). The organic layer was isolated and washed with saturated brine (50 mL) for 2 times and evaporated under reduced pressure. The resulting crude mixture purified by flash column chromatography (Silica gel, mesh size 100-200) eluting with 3% MeOH/ $CHCl_3$ to get compound **2b** (yield 75%) as white solid. $^1$H NMR (400 MHz, DMSO-$d_6$) δ in ppm 8.18 (s, 1H), 8.09 (s, 1H), 7.13 (s, 2H), 4.68 (sep, $J$ = 6.8 Hz, 1H), 1.47 (d, $J$ = 6.8 Hz, 6H). $^{13}$C NMR (101 MHz, DMSO-$d_6$) δ in ppm 156.50, 152.65, 149.57, 139.30, 119.64, 46.96, 22.65. HRMS (ESI) *m*/*z* (M + H)$^+$ calculated for $C_8H_{12}N_5$: 178.1093; found: 178.1084.

##### 4.5.3 Synthesis of 9-cyclopentyl-9H-purin-6-amine (2c)

Adenine **1** (1.5 g, 11.1 mmol), $K_2CO_3$ (1.8 g, 13.3 mmol), and Bromocyclopentane (1.2 mL, 11.1 mmol) were dissolved in 5 mL of dry DMF. The reaction mixture was stirred at room temperature for 12-16 hours. Progress of the reaction was thoroughly monitored through TLC. Upon completion, the reaction mixture was partitioned between EtOAc (50 ml) and ice-water (50 ml). The organic layer was isolated and washed with saturated brine (50 mL) for 2 times and evaporated under reduced pressure. The resulting crude mixture purified by flash column chromatography (Silica gel, mesh size 100-200) eluting with 2 % MeOH/ $CHCl_3$ to get compound **2c** (yield 78%) as white solid. $^1$H NMR (400 MHz, DMSO-$d_6$) δ 8.15 (s, 1H), 8.09 (s, 1H), 7.12 (s, 2H), 4.79 (p, $J$ = 7.6 Hz, 1H), 2.13-2.05 (m, 2H), 1.99-1.91 (m, 2H), 1.86-1.78 (m, 2H), 1.68-1.61 (m, 2H). $^{13}$C NMR (101 MHz, DMSO-$d_6$) δ 156.51, 152.67, 149.94, 139.83, 119.71, 55.86, 32.43, 24.07. HRMS (ESI) *m/z* $(M + H)^+$ calculated for $C_{10}H_{14}N_5$: 204.1249; found: 204.1240.

###### 4.5.4 Synthesis of 8-bromo-9-methyl-9H-purin-6-amine (3a)

Compound **2a** (1g, 6.70 mmol) was taken in a dry $CHCl_3$ (10 mL) and N-Bromosuccinimide (1.79g, 10.06 mmol) was added in reaction mixture and the reaction was performed by the **general procedure A**. Column chromatography (Eluent 30% EtOAc in Pet ether) was performed for purification and compound **3a** was appeared as light yellow coloured solid (yield 86%). $^1$H NMR (400 MHz, DMSO-$d_6$) δ in ppm 8.08 (s, 1H), 7.30 (s, 2H), 3.60 (s, 3H). $^{13}$C NMR (101 MHz, DMSO-$d_6$) δ in ppm 155.19, 153.24, 151.46, 127.59, 119.47, 30.65. HRMS (ESI) *m/z* $(M + H)^+$ calculated for $C_6H_7BrN_5$: 227.9885; found: 227.9883.

###### 4.5.5 Synthesis of 8-bromo-9-isopropyl-9H-purin-6-amine (3b)

Compound **2b** (1g, 5.64 mmol) was taken in a dry $CHCl_3$ (10 mL) and N-Bromosuccinimide (1.50g, 8.46 mmol) was added in reaction mixture and the reaction was performed by the **general procedure A**. Column chromatography (Eluent 50% EtOAc in Pet ether) was performed for purification and compound **3b** was appeared as light yellow coloured solid (yield 90%).$^1$H NMR (400 MHz, DMSO-$d_6$) δ 8.07 (s, 1H), 7.29 (s, 2H), 4.75 (sep, $J$ = 6.8 Hz, 1H), 1.56 (d, $J$ = 6.8 Hz, 6H). $^{13}$C NMR (101 MHz, DMSO-$d_6$) δ 155.40, 152.73, 151.05, 126.19, 120.15, 50.49, 21.09. HRMS (ESI) *m/z* $(M + H)^+$ calculated for $C_8H_{11}BrN_5$: 256.0198; found: 256.0187.

###### 4.5.6 Synthesis of 8-bromo-9-cyclopentyl-9H-purin-6-amine (3c)

Compound **2c** (1g, 4.92 mmol) was taken in a dry $CHCl_3$ (10 mL) and N-Bromosuccinimide (1.31g, 7.38 mmol) was added in reaction mixture and the reaction was performed by the **general procedure A**. Column chromatography (Eluent 35% EtOAc in Pet ether) was performed for purification and compound **3c** was appeared as light yellow coloured solid (yield 92%). $^{1}$H NMR (600 MHz, Chloroform-d) δ 8.26 (s, 1H), 5.89 (s, 2H), 4.91 (p, J = 8.6 Hz, 1H), 2.47-2.39 (m, 2H), 2.11-2.03 (m, 4H), 1.75-1.67 (m, 2H). $^{13}$C NMR (101 MHz, Chloroform-*d*) δ 154.40, 152.40, 151.23, 127.59, 120.59, 58.75, 30.54, 24.81.HRMS (ESI) *m*/*z* $(M + H)^+$ calculated for $C_{10}H_{13}BrN_5$: 282.0354; found: 282.0344.

##### 4.5.7 Synthesis of 8-(4-methoxyphenyl)-9-methyl-9H-purin-6-amine (4a)

Compound **3a** (1g, 4.40 mmol), 4-methoxyphenylboronic acid (0.803g, 5.28 mmol), $K_2CO_3$ (1.216g, 8.80 mmol) were taken in a pressure tube and dissolved in (9:1) mixture of Dioxane & water (15 mL). Solution was purged with Argon gas for 30 minutes. Then catalyst $Pd(PPh_3)_4$ (0.508g, 0.44 mmol) was added and the reaction was performed according to the **general procedure B**. Pure compound was isolated by column chromatography (Silica gel, mesh size 230-400) eluting (3% MeOH/ $CHCl_3$, V/V) to afford **4a** as yellow solid (yield 70%). $^{1}$H NMR (400 MHz, DMSO-*d*$_6$) δ in ppm 8.12 (s, 1H), 7.78-7.74 (m, 2H), 7.21 (s, 2H), 7.08-7.05 (m, 2H), 3.79 (s, 3H), 3.73 (s, 3H). $^{13}$C NMR (101 MHz, DMSO-*d*$_6$) δ in ppm 160.88, 155.86, 152.49, 151.84, 150.41, 130.96, 122.61, 119.00, 114.65, 55.86, 30.88. HRMS (ESI) *m*/*z* $(M + H)^+$ calculated for $C_{13}H_{14}N_5O$: 256.1198; found: 256.1205.

##### 4.5.8 Synthesis of 9-isopropyl-8-(4-methoxyphenyl)-9H-purin-6-amine (4b)

Compound **3b** (1g, 3.92 mmol), 4-methoxyphenylboronic acid (0.715g, 4.70 mmol), $K_2CO_3$ (1.083g, 7.84 mmol) were taken in a pressure tube and dissolved in (9:1) mixture of Dioxane & water (15 mL). Solution was purged with Argon gas for 30 minutes. Then catalyst $Pd(PPh_3)_4$ (0.453g, 0.39 mmol) was added and the reaction was performed according to the **general procedure B**. Pure compound was isolated by column chromatography (Silica gel, mesh size 230-400) eluting (1% MeOH/ $CHCl_3$, V/V) to afford **4b** as yellow solid (yield 68%).$^{1}$H NMR (400 MHz, Chloroform-*d*) δ 8.32 (s, 1H), 7.54 – 7.52 (m, 2H), 7.05 – 7.02 (m, 2H), 5.82 (s, 2H), 4.68 (sep, *J* = 6.9 Hz, 1H), 3.88 (s, 3H), 1.69 (d, *J* = 6.8 Hz, 6H). $^{13}$C NMR (101 MHz, Chloroform-*d*) δ 161.10, 154.98, 151.57, 151.36, 130.83, 122.53, 119.90, 114.44, 55.53, 49.59, 21.37. HRMS (ESI) *m*/*z* $(M + H)^+$ calculated for $C_{15}H_{18}N_5O$: 284.1511; found: 284.1502.

##### 4.5.9 Synthesis of 9-cyclopentyl-8-(4-methoxyphenyl)-9H-purin-6-amine (4c)

Compound **3c** (1g, 3.55 mmol), 4-methoxyphenylboronic acid (0.648g, 4.27 mmol), $K_2CO_3$ (0.981g, 7.1 mmol) were taken in a pressure tube and dissolved in (9:1) mixture of Dioxane & water (15 mL). Solution was purged with Argon gas for 30 minutes. Then catalyst $Pd(PPh_3)_4$ (0.404g, 0.35 mmol) was added and the reaction was performed according to the **general procedure B**. Pure compound was isolated by column chromatography (Silica gel, mesh size 230-400) eluting (90% EtOAc/ Pet ether,

V/V) to afford **4c** as yellow solid (yield 66%). $^{1}$H NMR (400 MHz, Chloroform-*d*) δ 8.32 (s, 1H), 7.56 – 7.54 (m, 2H), 7.05 – 7.03 (m, 2H), 5.79 (s, 2H), 4.71 (p, *J* = 8.7 Hz, 1H), 3.88 (s, 3H), 2.60 – 2.51 (m, 2H), 2.12 – 2.05 (m, 2H), 2.02 – 1.94 (m, 2H), 1.67 – 1.58 (m, 2H). 13C NMR (101 MHz, CHLOROFORM-D) δ 161.08, 154.98, 152.10, 151.56, 151.19, 130.86, 122.61, 119.98, 114.43, 58.05, 55.53, 30.96, 24.79. HRMS (ESI) *m/z* $(M + H)^+$ calculated for $C_{17}H_{20}N_5O$: 310.1668; found: 310.1665. Melting Point: 70 °C.

###### 4.5.10 Synthesis of 9-cyclopentyl-8-(4-methoxyphenyl)-9H-purin-6-amine (4d)

Compound **3a** (1g, 4.40 mmol), 2,4-dimethoxyphenylboronic acid (0.960g, 5.28 mmol), $K_2CO_3$ (1.216g, 8.80 mmol) were taken in a pressure tube and dissolved in (9:1) mixture of Dioxane & water (15 mL). Solution was purged with Argon gas for 30 minutes. Then catalyst $Pd(PPh_3)_4$ (0.508g, 0.44 mmol) was added and the reaction was performed according to the **general procedure B**. Pure compound was isolated by column chromatography (Silica gel, mesh size 230-400) eluting (2 % MeOH/ $CHCl_3$, V/V) to afford **4d** as yellow solid (yield 72%). $^{1}$H NMR (400 MHz, $CDCl_3$) δ in ppm 8.34 (s, 1H), 7.41 (d, J= 8.4Hz, 1H), 6.61 (dd, J= 8.4 Hz, 2.4 Hz, 1H), 6.55 (d, J= 2.4 Hz, 1H), 6.03 (s, 2H), 3.86 (s, 3H), 3.79 (s, 3H), 3.61 (s, 3H). $^{13}$C NMR (101 MHz, $CDCl_3$) δ in ppm 163.09, 158.91, 154.95, 152.36, 151.51, 150.17, 132.92, 119.34, 111.49, 105.33, 98.79, 55.65, 29.78. HRMS (ESI) *m/z* $(M + H)^+$ calculated for $C_{14}H_{16}N_5O_2$: 286.1304; found: 286.1315.

###### 4.5.11 Synthesis of 4-(6-amino-9-methyl-9H-purin-8-yl)benzonitrile (4e)

Compound **3a** (1g, 4.40 mmol), 4-cyanophenylboronic acid (0.775g, 5.28 mmol), $K_2CO_3$ (1.216g, 8.80 mmol) were taken in a pressure tube and dissolved in (9:1) mixture of Dioxane & water (15 mL). Solution was purged with Argon gas for 30 minutes. Then catalyst $Pd(PPh_3)_4$ (0.508g, 0.44 mmol) was added and the reaction was performed according to the **general procedure B**. Pure compound was isolated by column chromatography (Silica gel, mesh size 230-400) eluting (5 % MeOH/ $CHCl_3$, V/V) to afford **4e** as yellow solid (yield 70%). HRMS (ESI) *m/z* $(M + H)^+$ calculated for $C_{13}H_{11}N_6$: 251.1045; found: 251.1046.

###### 4.5.12 Synthesis of 8-(4-chlorophenyl)-9-methyl-9H-purin-6-amine (4f)

Compound **3a** (1g, 4.40 mmol), 4-chlorophenylboronic acid (0.825g, 5.28 mmol), $K_2CO_3$ (1.216g, 8.80 mmol) were taken in a pressure tube and dissolved in (9:1) mixture of Dioxane & water (15 mL). Solution was purged with Argon gas for 30 minutes. Then catalyst $Pd(PPh_3)_4$ (0.508g, 0.44 mmol) was added and the reaction was performed according to the **general procedure B**. Pure compound was isolated by column chromatography (Silica gel, mesh size 230-400) eluting (5 % MeOH/ $CHCl_3$, V/V) to afford **4f** as yellow solid (yield 66%). $^1$H NMR (400 MHz, DMSO-$d_6$) δ in ppm 8.14 (s, 1H), 7.87-7.83 (m, 2H), 7.61-7.58 (m, 2H), 7.26 (s, 2H), 3.75 (s, 3H). $^{13}$C NMR (101 MHz, DMSO-$d_6$) δ in ppm 156.26, 153.09, 151.89, 149.22, 135.13, 131.23, 129.33, 129.24, 119.14, 30.87. HRMS (ESI) *m*/*z* (M + H)$^+$ calculated for $C_{12}H_{11}ClN_5$: 260.0703; found: 260.0696.

##### 4.5.13 Synthesis of N-(8-(4-methoxyphenyl)-9-methyl-9H-purin-6-yl)-4-methylbenzamide (AS1)

Compound **4a** (0.1g, 0.39 mmol) was dissolved in dry THF (5mL) and then LiHMDS (0.78mL, 0.78 mmol) was added dropwise at ice-cold condition keeping nitrogen atmosphere and the reaction mixture was allowed to stir at room temperature for 15 minutes. Then a solution of 4-methylbenzoyl chloride (0.062mL, 0.47 mmol) in dry THF (1 mL) was added dropwise at ice-cold condition and then stirred at room temperature for another 2 hrs. The progress of the reaction was monitored by checking TLC. After completion, reaction mass was washed with water and extracted with ethyl acetate. The organic layer was washed with brine solution, dried over $Na_2SO_4$ and was evaporated under vacuum. The resulting crude mixture was purified by flash column chromatography (Silica gel, mesh size 230-400) eluting with 80% EtOAc/ Pet ether to get compound **AS1** (yield 68%) as white solid. $^1$H NMR (400 MHz, $CDCl_3$) δ in ppm 9.27 (s, 1H), 8.79 (s, 1H), 7.94-7.88 (m, 2H), 7.77-7.71 (m, 2H), 7.29-7.23 (m, 2H), 7.06-6.99 (m, 2H), 3.91 (s, 3H), 3.86 (s, 3H), 2.40 (s, 3H). $^{13}$C NMR (101 MHz, $CDCl_3$) δ in ppm 161.66, 153.79, 152.21, 148.75, 143.33, 131.27, 130.84, 129.47, 128.00, 122.41, 121.21, 114.58, 55.57, 31.01, 21.66. HRMS (ESI) *m*/*z* (M + H)$^+$ calculated for $C_{21}H_{20}N_5O_2$: 371.1617; found: 371.1624.

##### 4.5.14 Synthesis of 1-(8-(4-methoxyphenyl)-9-methyl-9H-purin-6-yl)-3-(p-tolyl)urea (AS2)

Compound **4a** (0.1g, 0.39 mmol) was dissolved in dry THF (5mL) and then LiHMDS (0.78mL, 0.78 mmol) was added dropwise at ice-cold condition keeping nitrogen atmosphere and the reaction mixture was allowed to stir at room temperature for 15 minutes. Then a solution of required 4-methylphenylisocyanate (0.059mL, 0.47 mmol) in dry THF (1 mL) was added dropwise at ice-cold condition and then stirred at room temperature for another 2 hrs. The progress of the reaction was monitored by checking TLC. After completion, reaction mass was washed with water and extracted with ethyl acetate. The organic layer was washed with brine solution, dried over $Na_2SO_4$ and was evaporated under vacuum. The resulting crude mixture was purified by flash column chromatography (Silica gel, mesh size 230-400) eluting with 2% MeOH/ $CHCl_3$to get compound **AS2** (yield 72%) as white solid. $^1$H NMR (400 MHz, $CDCl_3$) δ in ppm 8.80 (s, 3H), 7.93-7.89 (m, 3H), 7.76-7.73 (m, 2H), 7.28-7.25 (m, 2H), 7.06-7.02 (m, 2H), 3.92 (s, 3H), 3.87 (s, 3H), 2.40 (s, 3H). $^{13}$C NMR (101 MHz, $CDCl_3$) δ in ppm 161.67, 153.79, 152.16, 148.78, 143.35, 131.28, 130.85, 130.63, 129.48, 128.03, 122.37, 121.16, 114.60, 55.58, 31.03, 21.68. HRMS (ESI) *m*/*z* (M + H)$^+$ calculated for $C_{21}H_{21}N_6O_2$: 389.1726

##### 4.5.15 Synthesis of N-(8-(4-methoxyphenyl)-9-methyl-9H-purin-6-yl)-4-methylbenzenesulfonamide (AS3)

Compound **4a** (0.1g, 0.39 mmol) was dissolved in dry DMF (5mL) and then NaH (0.018g, 0.78 mmol) was added portion wise at ice-cold condition keeping nitrogen atmosphere and the reaction mixture was allowed to stir at room temperature for 15 minutes. Then p-toluenesulfonyl chloride (0.089g, 0.47 mmol) were added portion wise at ice-cold condition and the reaction was performed according to the general procedure **C**. Pure compound was isolated by using column chromatography in 230-400 silica mesh size and with 80 % EtOAc in Pet ether (V/V) as eluent to obtain compound **AS3** as white solid (yield 80%). $^{1}H$ NMR (400 MHz, $CDCl_3$) δ in ppm 8.27 (s, 1H), 8.03-7.92 (m, 2H), 7.74-7.66 (m, 2H), 7.27-7.29 (m, 2H), 7.03-6.98 (m, 2H), 3.86 (m, 3H), 3.86 (m, 3H), 2.36 (s, 3H). $^{13}C$ NMR (101 MHz, $CDCl_3$) δ in ppm 161.31, 131.11, 129.81, 121.72, 114.76, 100.00, 55.93, 31.45, 21.49. HRMS (ESI) *m/z* $(M + H)^+$ calculated for $C_{20}H_{20}N_5O_3S$: 410.1287; found: 410.1274.

##### 4.5.16 Synthesis of N-(9-isopropyl-8-(4-methoxyphenyl)-9H-purin-6-yl)-4-methylbenzenesulfonamide (AS4)

Compound **4b** (0.1g, 0.35 mmol) was dissolved in dry DMF (5mL) and then NaH (0.017g, 0.70 mmol) was added portion wise at ice-cold condition keeping nitrogen atmosphere and the reaction mixture was allowed to stir at room temperature for 15 minutes. Then p-toluenesulfonyl chloride (0.080g, 0.42 mmol) were added portion wise at ice-cold condition and the reaction was performed according to the general procedure **C**. Pure compound was isolated by using column chromatography in 230-400 silica mesh size and with 67 % EtOAc in Pet ether (V/V) as eluent to obtain compound **AS4** as white solid (yield 76%). $^{1}H$ NMR (400 MHz, $CDCl_3$) δ in ppm 8.25 (br.s), 8.06-7.89 (m, 2H), 7.54-7.49 (m, 2H), 7.24-7.21 (m, 2H), 7.02-6.98 (m, 2H), 4.74-4.67 (m, 1H), 3.85 (s, 3H), 2.36 (s, 3H), 1.65 (d, *J*= 7.2 Hz, 6H). $^{13}C$ NMR (101 MHz, $CDCl_3$) δ in ppm 161.38, 148.08, 131.02, 129.34, 127.34, 121.48, 114.40, 55.53, 50.22, 21.62, 21.42, 1.10. HRMS (ESI) *m/z* $(M + H)^+$ calculated for $C_{22}H_{24}N_5O_3S$: 438.1600; found: 438.1611.

##### 4.5.17 Synthesis of N-(9-cyclopentyl-8-(4-methoxyphenyl)-9H-purin-6-yl)-4-methylbenzenesulfonamide (AS5)

Compound **4c** (0.1g, 0.32 mmol) was dissolved in dry DMF (5mL) and then NaH (0.015g, 0.64 mmol) was added portion wise at ice-cold condition keeping nitrogen atmosphere and the reaction mixture was allowed to stir at room temperature for 15 minutes. Then p-toluenesulfonyl chloride (0.074g, 0.38 mmol) were added portion wise at ice-cold condition and the reaction was performed according to the general procedure **C**. Pure compound was isolated by using column chromatography in 230-400 silica mesh size and with 58 % EtOAc in Pet ether (V/V) as eluent to obtain compound **AS5** as white solid (yield 68%). $^{1}$H NMR (400 MHz, $CDCl_3$) δ in ppm 88.06-7.89 (m, 2H), 7.55-7.52 (m, 2H), 7.24-7.22 (m, 2H), 7.03-6.98 (m, 2H), 4.74 (p, 1H), 3.86 (s, 3H), 2.47-2.39 (m, 2H), 2.36 (s, 3H), 2.10-1.97 (m, 4H), 1.66-1.59 (m, 2H). $^{13}$C NMR (101 MHz, $CDCl_3$) δ in ppm 161.35, 131.08, 129.35, 114.38, 58.35, 55.53, 31.36, 24.95, 21.62. HRMS (ESI) *m/z* $(M + H)^+$ calculated for $C_{24}H_{26}N_5O_3S$: 464.1756; found: 464.1747.

###### 4.5.18 Synthesis of 4-methoxy-N-(8-(4-methoxyphenyl)-9-methyl-9H-purin-6-yl)benzenesulfonamide (AS6)

Compound **4a** (0.1g, 0.39 mmol) was dissolved in dry DMF (5mL) and then NaH (0.018g, 0.78 mmol) was added portion wise at ice-cold condition keeping nitrogen atmosphere and the reaction mixture was allowed to stir at room temperature for 15 minutes. Then 4-Methoxybenzenesulfonyl chloride (0.097g, 0.47 mmol) were added portion wise at ice-cold condition and the reaction was performed according to the general procedure **C**. Pure compound was isolated by using column chromatography in 230-400 silica mesh size and with 75% EtOAc in Pet ether (V/V) as eluent to obtain compound **AS6** as white solid (yield 62%). $^{1}$H NMR (400 MHz, DMSO-$d_6$) δ in ppm 12.73 (s, 1H), 8.31 (s, 1H), 8.00-7.87 (m, 2H), 7.83-7.72 (m, 2H), 7.15-7.06 (m, 2H), 7.05-6.97 (m, 2H), 3.81 (s, 3H), 3.78 (s, 3H), 3.77 (s, 3H). $^{13}$C NMR 13C NMR (151 MHz, Chloroform-*d*) δ 161.64, 153.14, 147.86, 130.64, 120.83, 114.59, 113.88, 55.63, 55.53, 31.03. HRMS (ESI) *m/z* $(M + H)^+$ calculated for $C_{20}H_{20}N_5O_4S$: 426.1236; found: 426.1236.

###### 4.5.19 Synthesis of N-(8-(4-methoxyphenyl)-9-methyl-9H-purin-6-yl)-4-(trifluoromethyl)benzene sulfonamide (AS7)

Compound **4a** (0.1g, 0.39 mmol) was dissolved in dry DMF (5mL) and then NaH (0.018g, 0.78 mmol) was added portion wise at ice-cold condition keeping nitrogen atmosphere and the reaction mixture was allowed to stir at room temperature for 15 minutes. Then 4-(Trifluoromethyl)benzenesulfonyl chloride (0.115g, 0.47 mmol) were added portion wise at ice-cold condition and the reaction was performed according to the general procedure **C**. Pure compound was isolated by using column chromatography in 230-400 silica mesh size and with 75% EtOAc in Pet ether (V/V) as eluent to obtain compound **AS7** as white solid (yield 68%). $^{1}$H NMR (400 MHz, DMSO-$d_6$) δ in ppm 13.04 (s, 1H), 8.35 (s, 1H), 8.17-8.10 (m, 2H), 7.92-7.87 (m, 2H), 7.75-7.68 (m, 2H), 7.10-7.04 (m, 2H), 3.81 (s, 3H), 3.78 (s, 3H). $^{13}$C NMR (101 MHz, DMSO-$d_6$) δ in ppm 161.37, 148.49, 131.10, 126.62, 126.58, 125.51, 122.80, 121.56, 114.76, 100.00, 55.94, 31.53. HRMS (ESI) *m/z* $(M + H)^+$ calculated for $C_{20}H_{17}F_3N_5O_3S$: 464.1004; found: 464.0997.

##### 4.5.20 Synthesis of N-(8-(4-methoxyphenyl)-9-methyl-9H-purin-6-yl)cyclohexanesulfonamide (AS8)

Compound **4a** (0.1g, 0.39 mmol) was dissolved in dry DMF (5mL) and then NaH (0.018g, 0.78 mmol) was added portion wise at ice-cold condition keeping nitrogen atmosphere and the reaction mixture was allowed to stir at room temperature for 15 minutes. Then 4-Cyclopentylbenzenesulfonyl chloride (0.139g, 0.47 mmol) were added portion wise at ice-cold condition and the reaction was performed according to the general procedure **C**. Pure compound was isolated by using column chromatography in 230-400 silica mesh size and with 45 % EtOAc in Pet ether (V/V) as eluent to obtain compound **AS8** as white solid (yield 78%). $^1$H NMR (400 MHz, Chloroform-*d*) δ in ppm 7.75-7.70 (m, 2H), 7.05-7.00 (m, 2H), 3.89 (s, 3H), 3.87 (s, 3H), 2.31-2.23 (m, 2H), 1.90-1.82 (m, 2H), 1.71-1.57 (m, 3H), 1.40-1.07 (m, 4H). $^{13}$C NMR (101 MHz, Chloroform-*d*) δ in ppm 161.52, 130.80, 121.13, 114.45, 68.04, 65.92, 62.27, 60.46, 55.54, 31.15, 15.33, 0.06. HRMS (ESI) *m/z* $(M + H)^+$ calculated for $C_{19}H_{24}N_5O_3S$: 402.1600; found: 402.1593.

##### 4.5.21 Synthesis of N-(8-(2,4-dimethoxyphenyl)-9-methyl-9H-purin-6-yl)-4-methylbenzenesulfonamide (AS9)

Compound **4d** (0.1g, 0.35 mmol) was dissolved in dry DMF (5mL) and then NaH (0.017g, 0.70 mmol) was added portion wise at ice-cold condition keeping nitrogen atmosphere and the reaction mixture was allowed to stir at room temperature for 15 minutes. Then p-toluenesulfonyl chloride (0.080g, 0.42 mmol) were added portion wise at ice-cold condition and the reaction was performed according to the general procedure **C**. Pure compound was isolated by using column chromatography in 230-400 silica mesh size and with 80% EtOAc in Pet ether (V/V) as eluent to obtain compound **AS9** as white solid (yield 76%). $^1$H NMR (400 MHz, Chloroform-*d*) δ in ppm 8.30 (br.s, 1H), 8.08-7.92 (m, 2H), 7.46-7.42 (m,1H), 7.24-7.21 (m, 2H), 6.62-6.58 (m, 1H), 6.55-6.51 (m, 1H), 3.86 (s, 3H), 3.78 (s, 3H), 3.62 (s, 3H), 2.36 (s, 3H). $^{13}$C NMR (101 MHz, Chloroform-*d*) δ in ppm 176.07, 171.24, 163.34, 158.75, 147.83, 129.32, 110.79, 105.40, 100.00, 98.70, 60.47, 55.67, 30.16, 21.61, 14.27. HRMS (ESI) *m/z* $(M + H)^+$ calculated for $C_{21}H_{22}N_5O_4S$: 440.1392; found: 440.1390.

##### 4.5.22 Synthesis of N-(8-(4-cyanophenyl)-9-methyl-9H-purin-6-yl)-4-methylbenzenesulfonamide (AS10)

Compound **4e** (0.1g, 0.39 mmol) was dissolved in dry DMF (5mL) and then NaH (0.018g, 0.78 mmol) was added portion wise at ice-cold condition keeping nitrogen atmosphere and the reaction mixture was allowed to stir at room temperature for 15 minutes. Then p-toluenesulfonyl chloride (0.080g, 0.47 mmol) were added portion wise at ice-cold condition and the reaction was performed according to the general procedure **C**. Pure compound was isolated by using column chromatography in 230-400 silica mesh size and with 95 % EtOAc in Pet ether (V/V) as eluent to obtain compound **AS10** as white solid (yield 80%). $^1$H NMR (400 MHz, Chloroform-*d*) δ in ppm 8.27 (br.s, 1H), 7.97-7.87 (m, 2H), 7.87-7.84 (m, 2H), 7.81-7.75 (m, 2H), 7.22-7.18 (m, 2H), 3.84 (s, 3H), 2.31 (s, 3H). $^{13}$C NMR (101 MHz, Chloroform-*d*) δ in ppm 136.87, 136.72, 133.68, 133.33, 121.87, 118.10, 35.09, 25.37, 17.95. HRMS (ESI) *m*/*z* $(M + H)^+$ calculated for $C_{20}H_{17}N_6O_2S$: 405.1134; found:405.1126.

###### 4.5.23 Synthesis of N-(8-(4-chlorophenyl)-9-methyl-9H-purin-6-yl)-4-methylbenzenesulfonamide (AS11)

Compound **4f** (0.1g, 0.38 mmol) was dissolved in dry DMF (5mL) and then NaH (0.018g, 0.77 mmol) was added portion wise at ice-cold condition keeping nitrogen atmosphere and the reaction mixture was allowed to stir at room temperature for 15 minutes. Then p-toluenesulfonyl chloride (0.078g, 0.46 mmol) were added portion wise at ice-cold condition and the reaction was performed according to the general procedure **C**. Pure compound was isolated by using column chromatography in 230-400 silica mesh size and with 90 % EtOAc in Pet ether (V/V) as eluent to obtain compound **AS11** as white solid (yield 72%). $^1$H NMR (400 MHz, Chloroform-*d*) δ in ppm 8.27 (br.s, 1H), 8.03-7.94 (m, 2H), 7.73-7.68 (m, 2H), 7.52-7.47 (m, 2H), 7.28-7.25 (m, 2H), 3.87 (s, 3H), 2.37 (s, 3H). $^{13}$C NMR (101 MHz, DMSO-$d_6$) δ 135.78, 131.29, 129.86, 129.43, 128.22, 100.00, 31.42, 21.45. HRMS (ESI) *m*/*z* $(M + H)^+$ calculated for $C_{19}H_{17}ClN_5O_2S$: 414.0791; found: 414.0781.

### 5. NMR SPECTROSCOPIC DATA

#### 5.1 $^{1}$H NMR of compound 2a (400 MHz, DMSO-$d_6$):

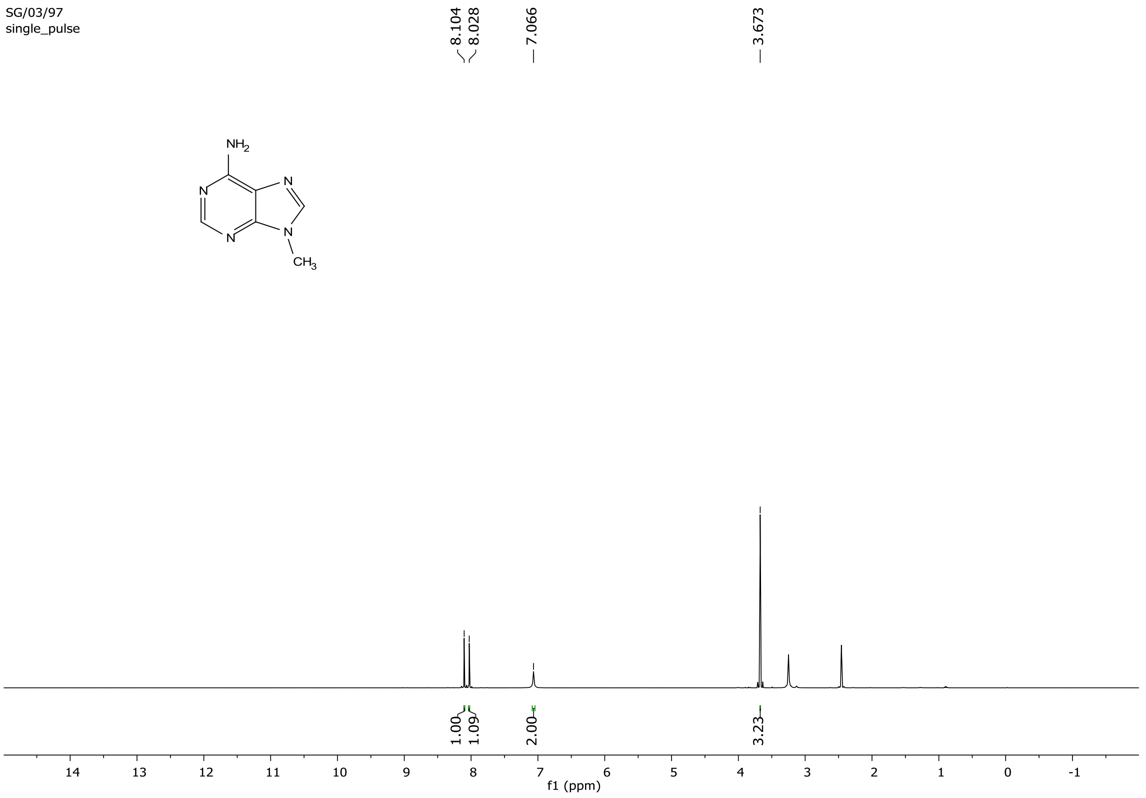

#### 5.2 $^{13}$C NMR of compound 2a (400 MHz, DMSO-$d_6$):

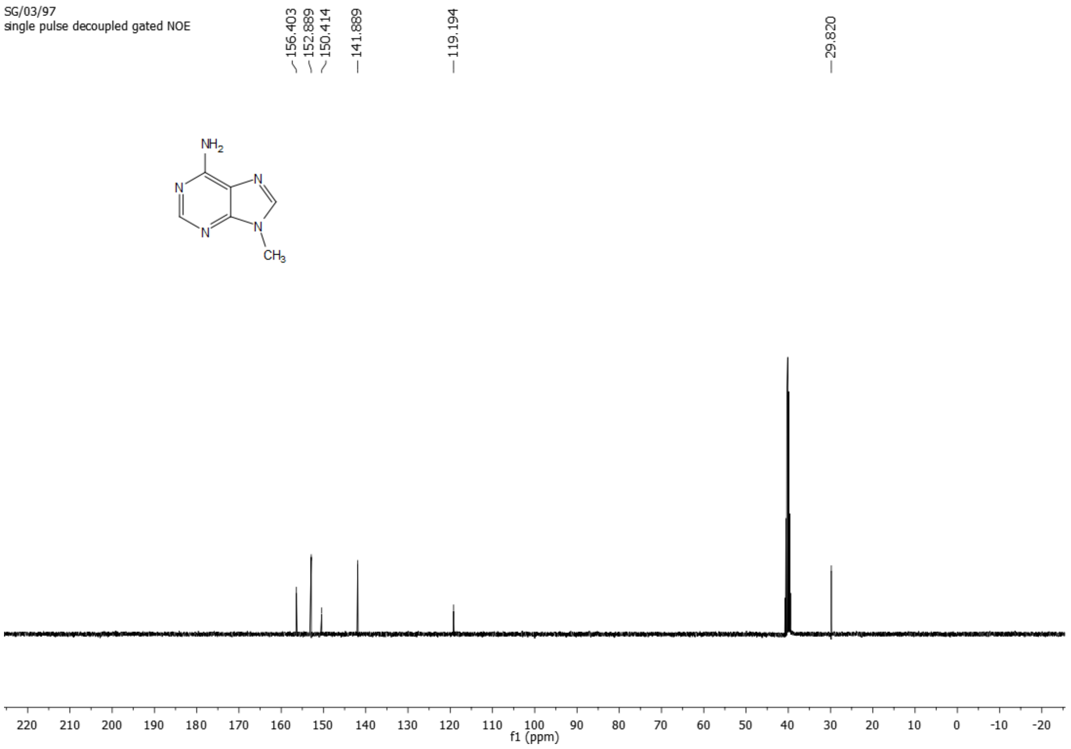

#### 5.3 $^{1}$H NMR of compound 2b (400 MHz, DMSO-$d_6$):

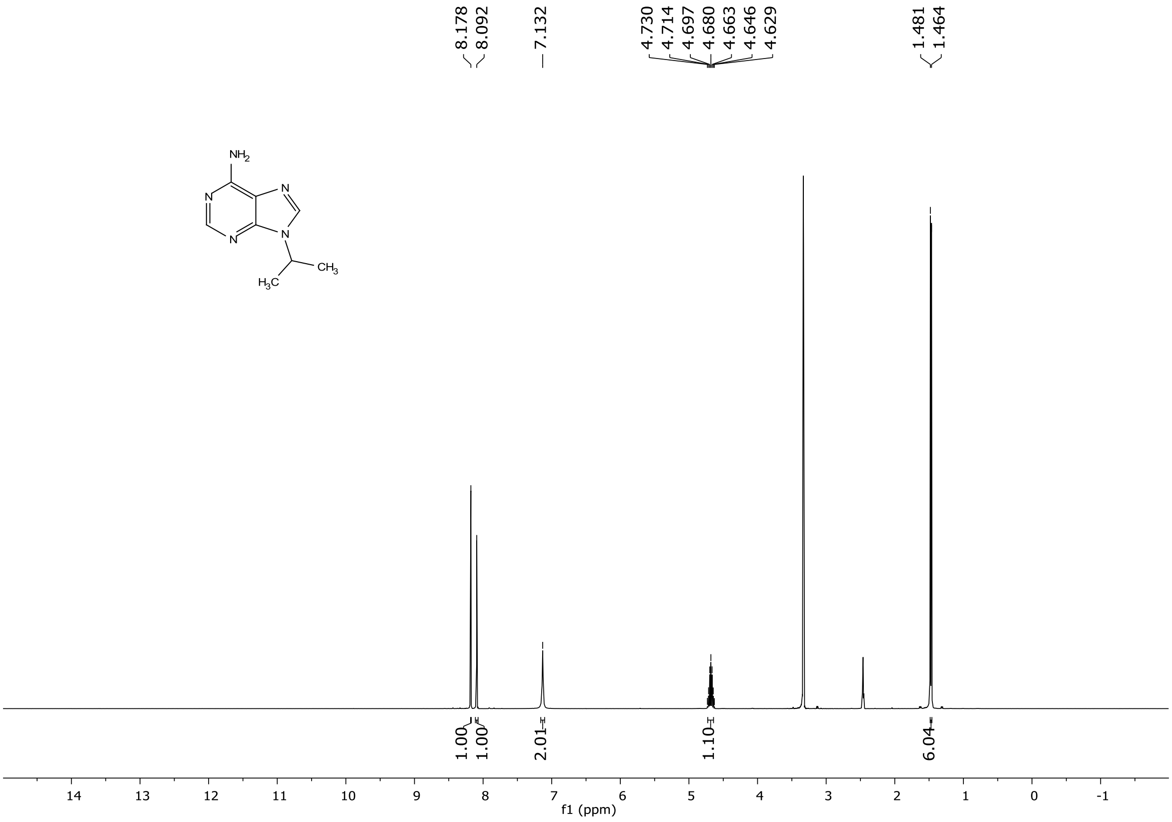

#### 5.4 $^{13}$C NMR of compound 2b (400 MHz, DMSO-$d_6$):

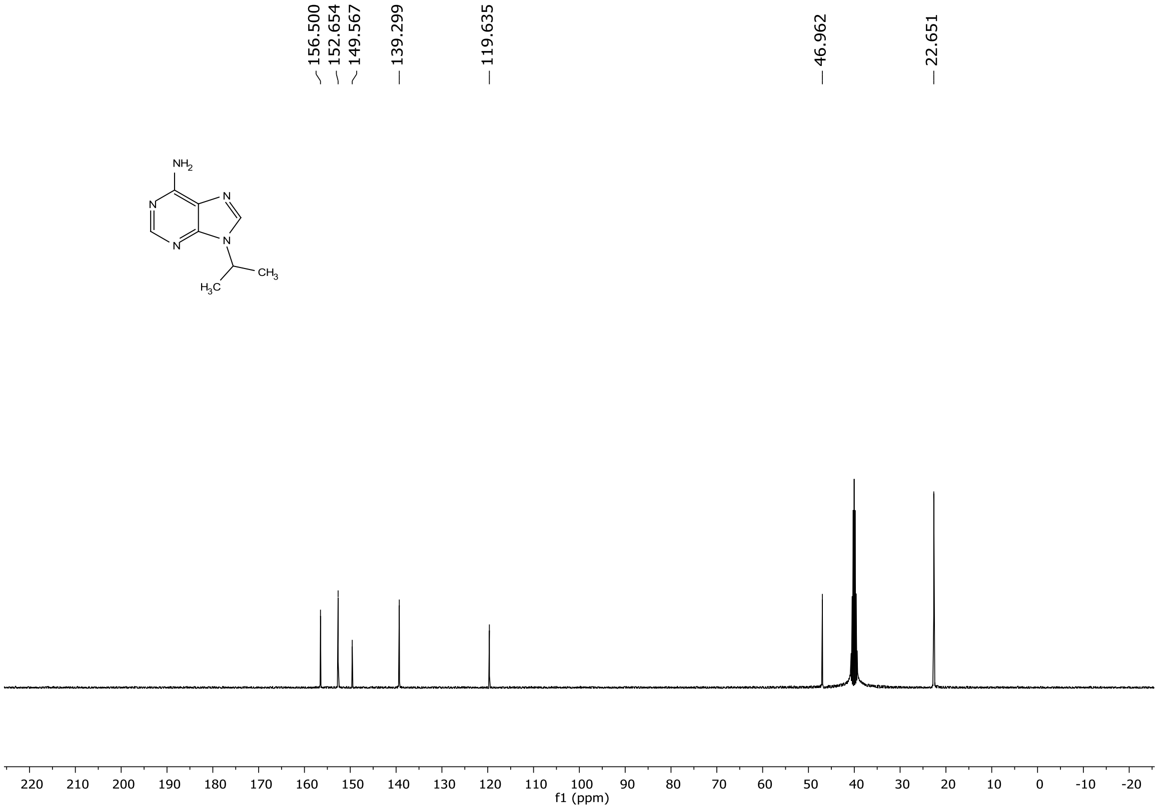

#### 5.5 $^{1}$H NMR of compound 2c (400 MHz, DMSO-$d_6$):

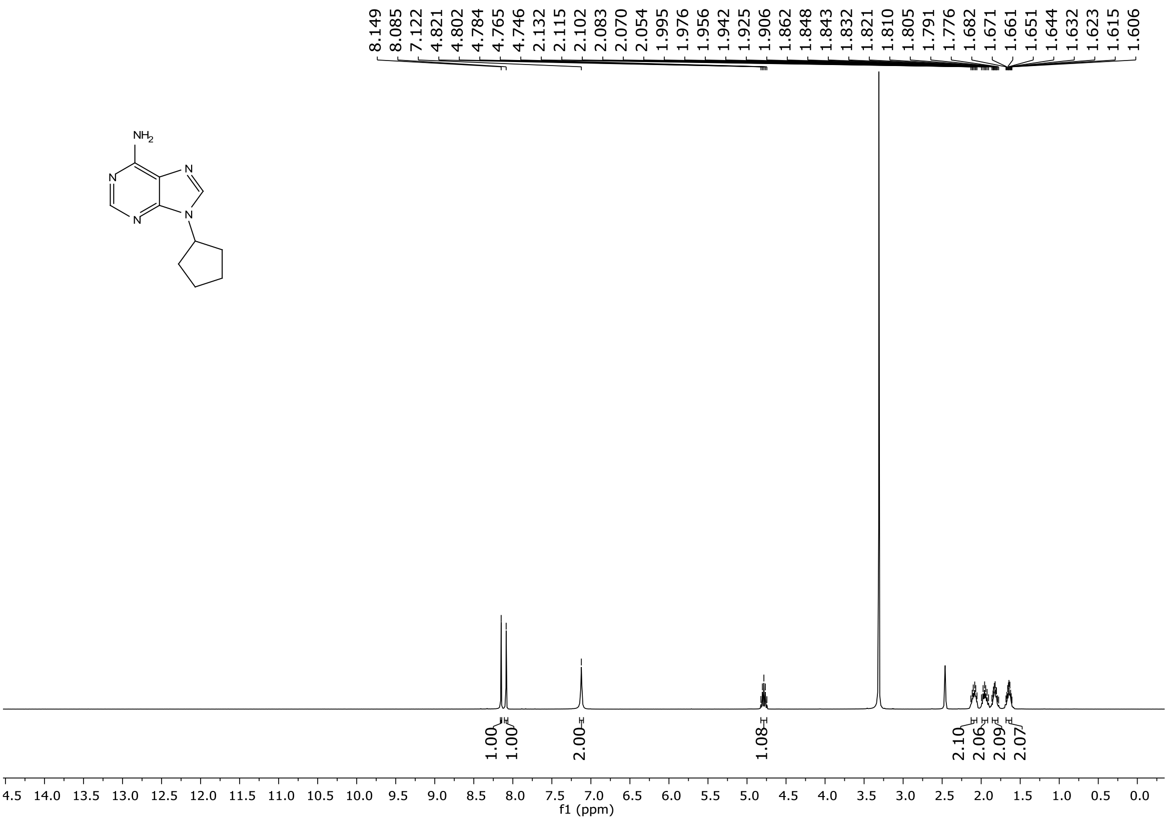

#### 5.6 $^{13}$C NMR of compound 2c (101 MHz, DMSO-$d_6$):

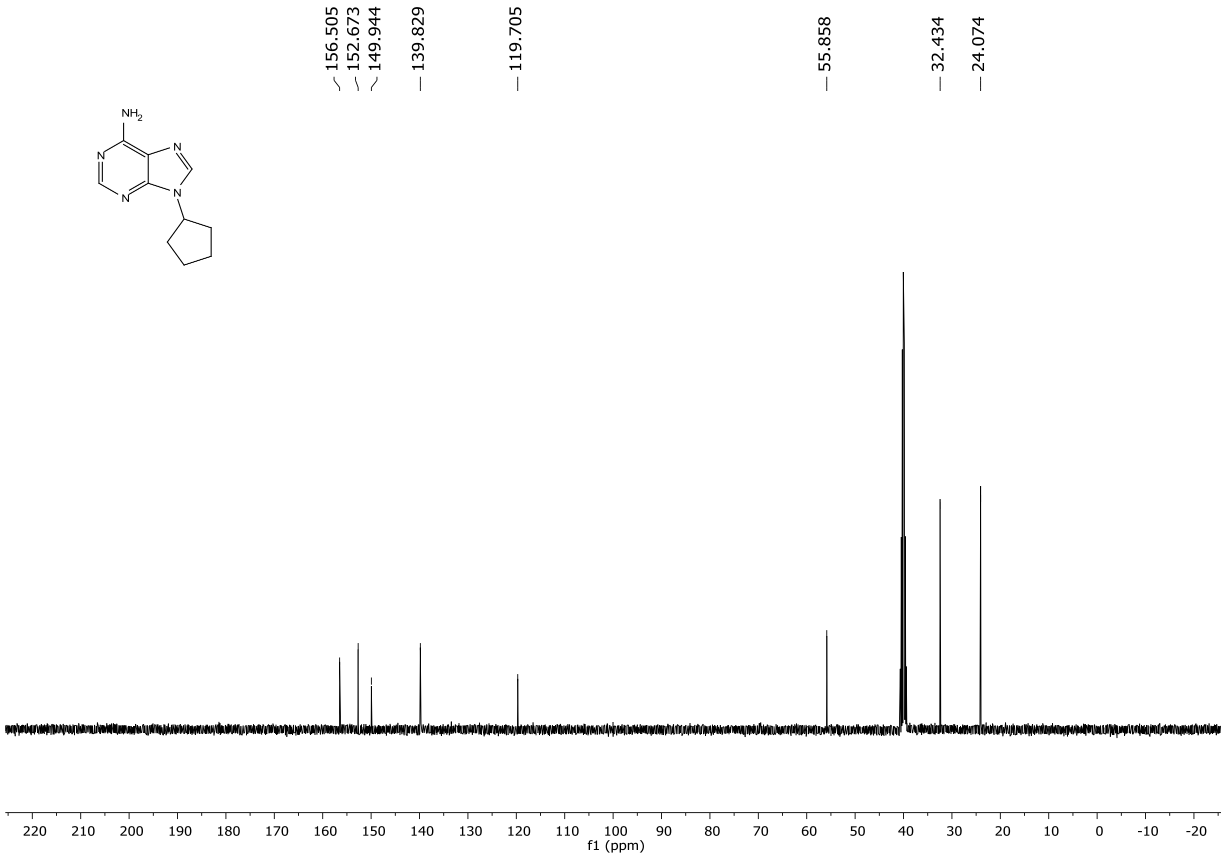

#### 5.7 $^{1}$H NMR of compound 3a (600 MHz, DMSO-$d_6$):

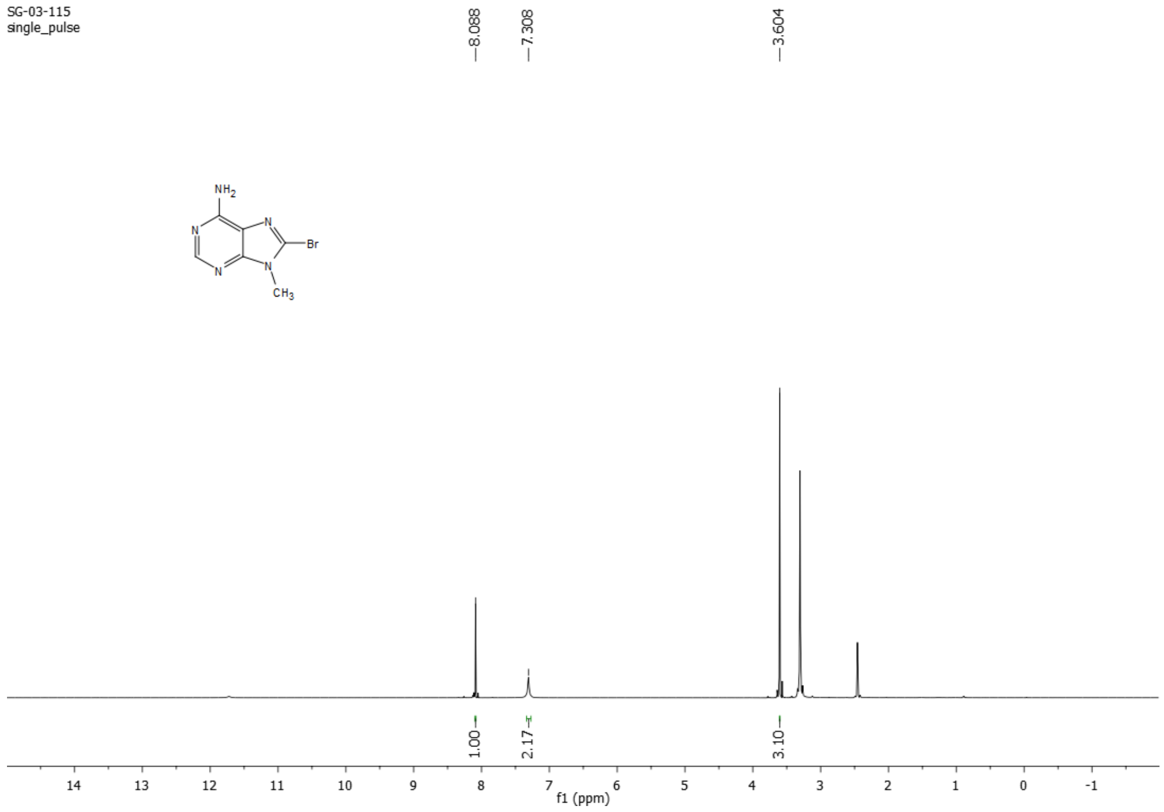

#### 5.8 $^{13}$C NMR of compound 3a (101 MHz, DMSO-$d_6$):

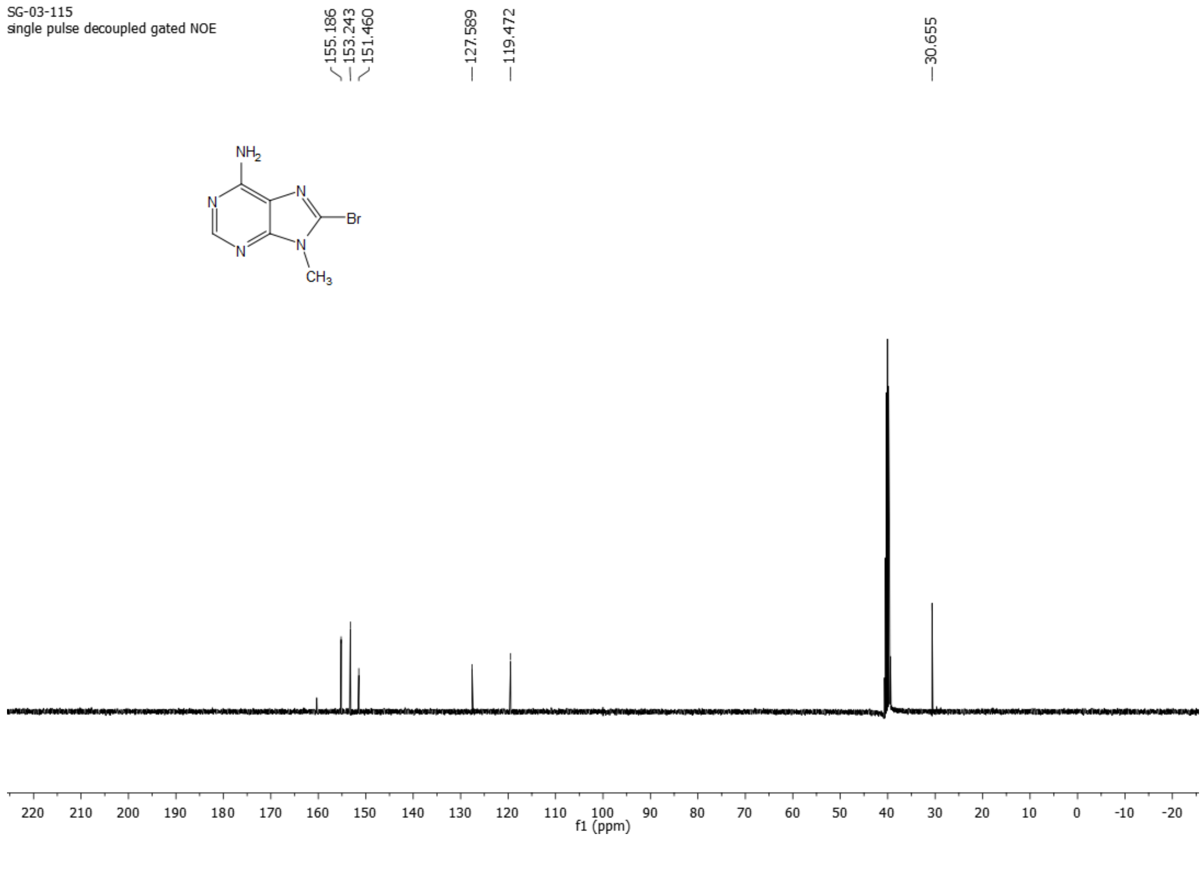

#### 5.9 [1]H NMR of compound 3b (400 MHz, DMSO-$d_6$):

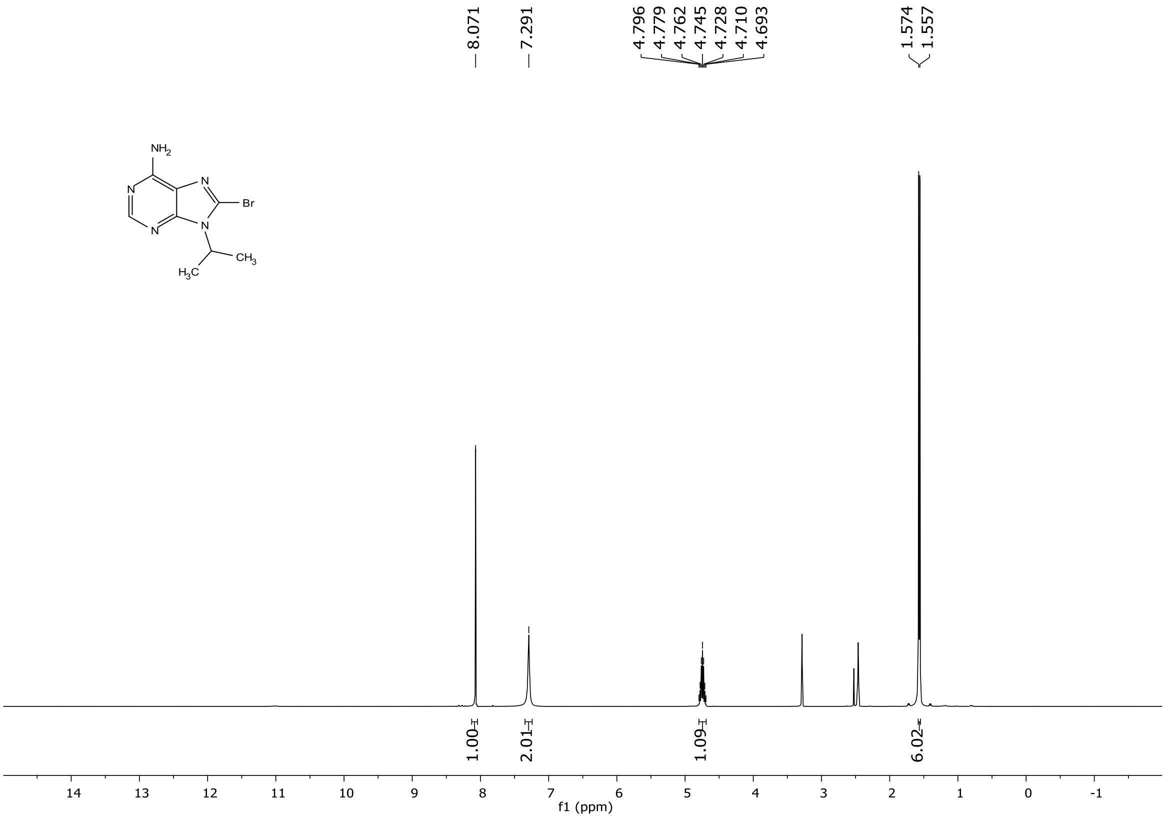

#### 5.10 [13]C NMR of compound 3b (101 MHz, DMSO-$d_6$):

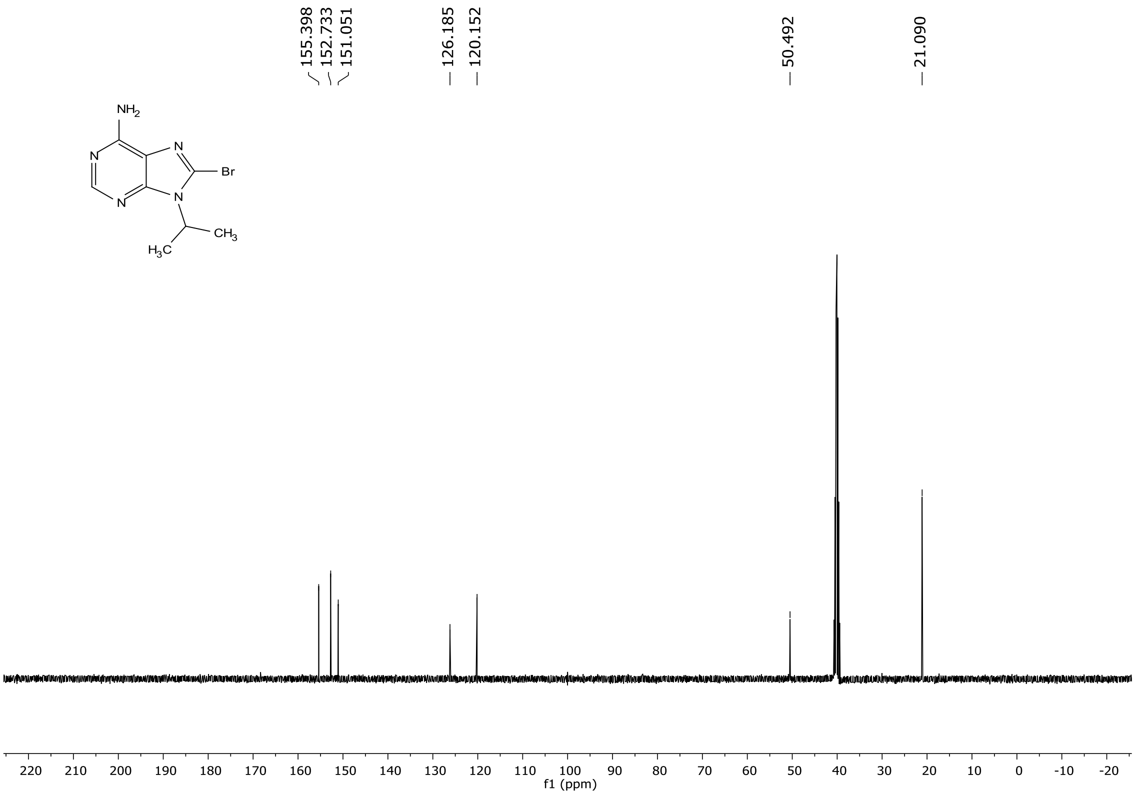

#### 5.11 $^{1}$H NMR of compound 3c (600 MHz, Chloroform-*d*):

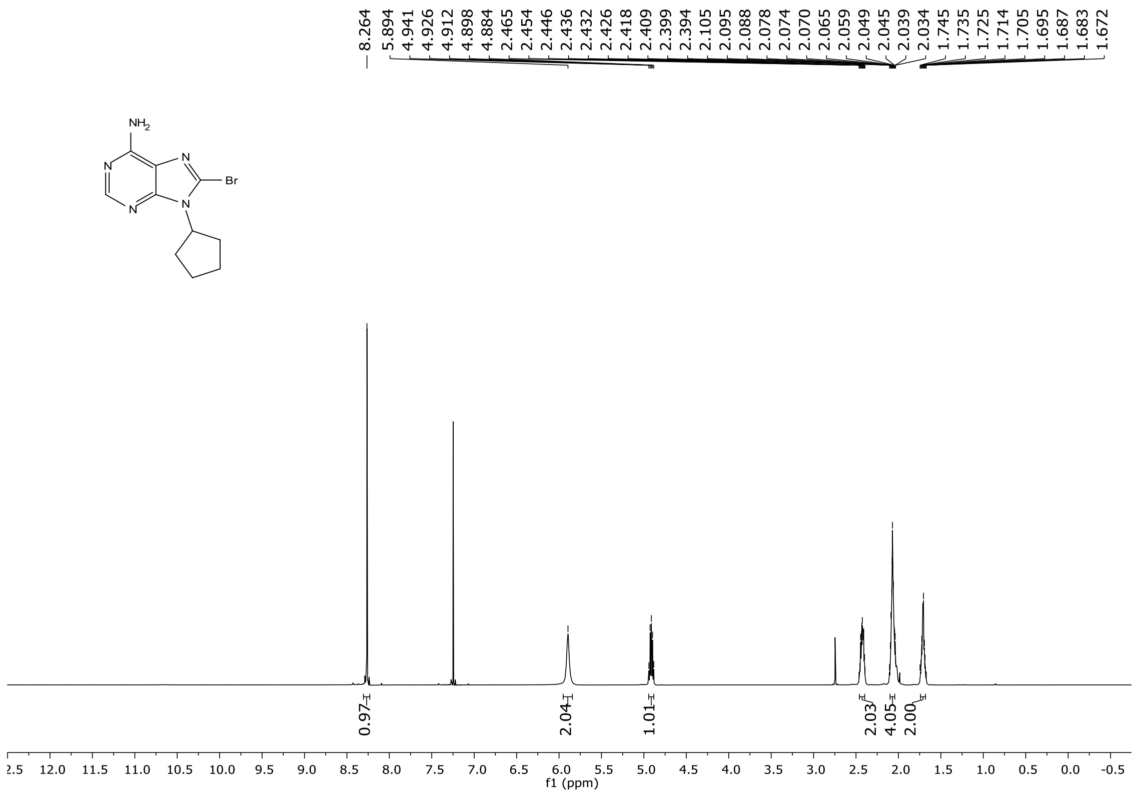

#### 5.12 $^{13}$C NMR of compound 3c (101 MHz, Chloroform-*d*):

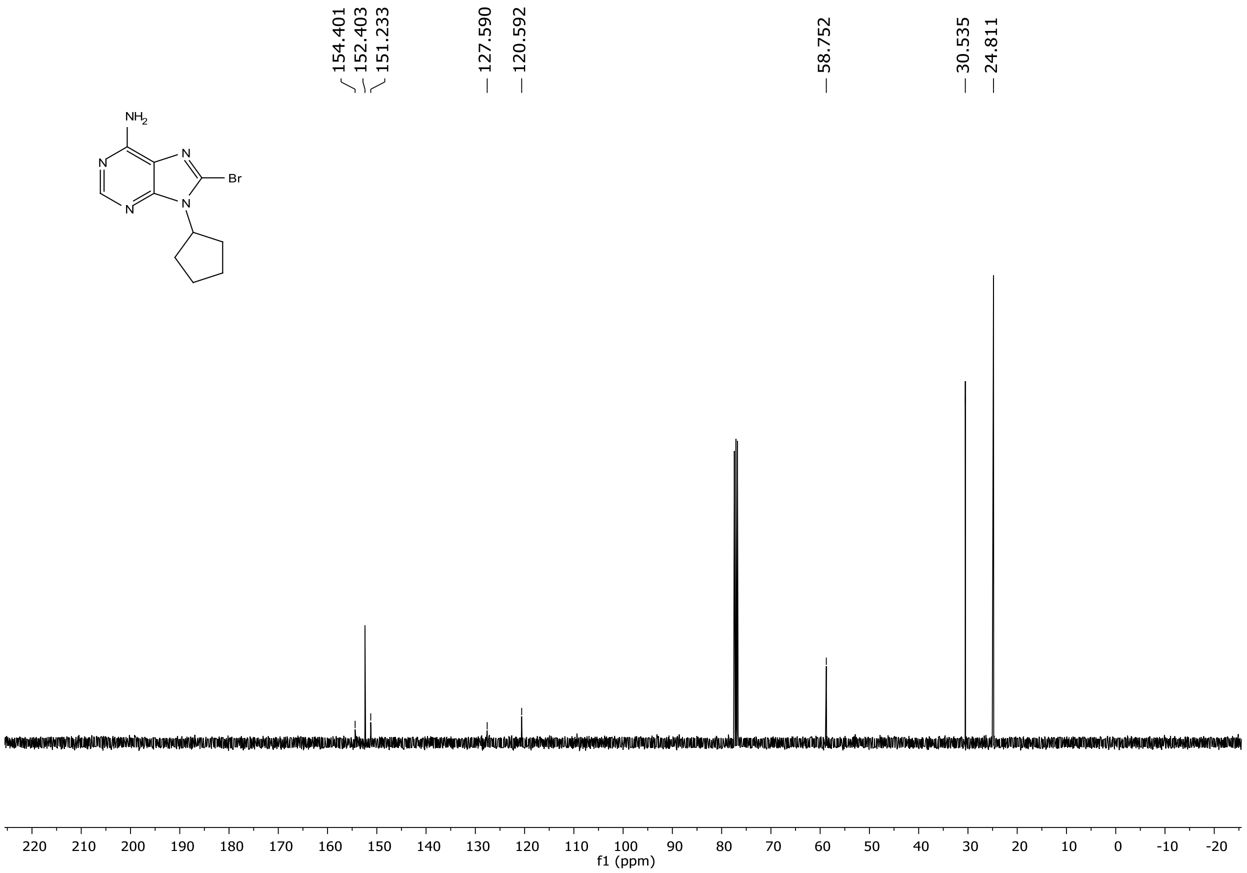

#### 5.13 $^{1}H$ NMR of compound 4a (400 MHz, DMSO-$d_6$):

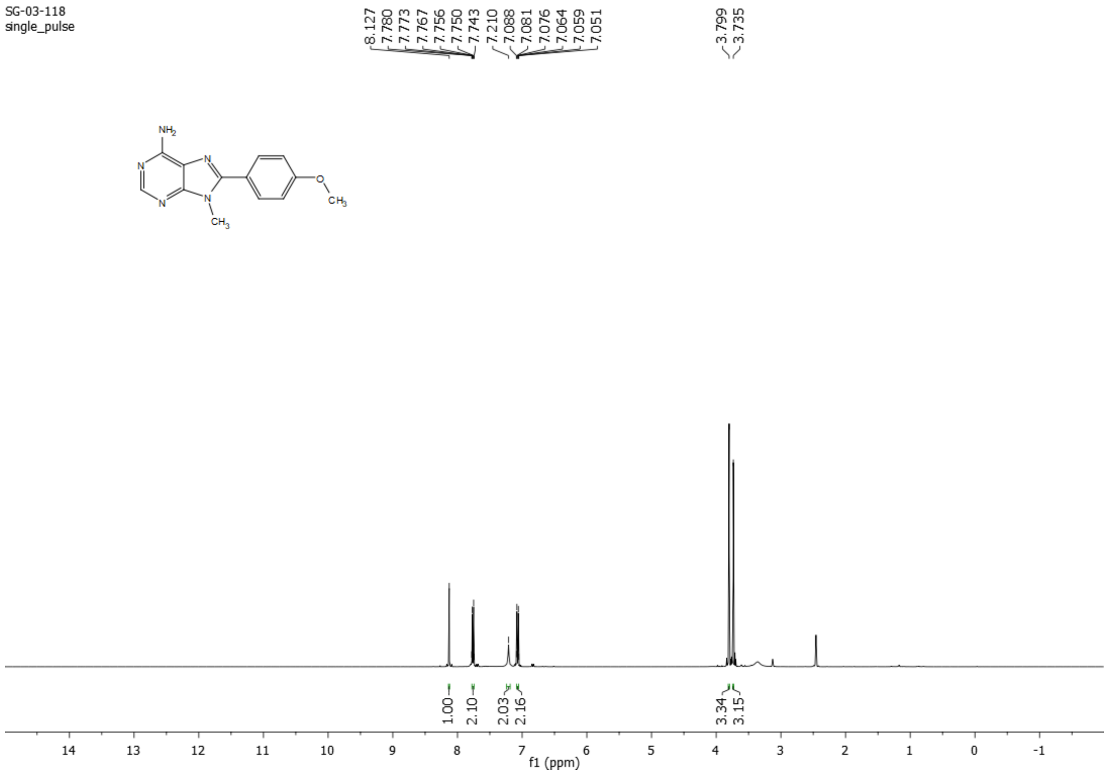

#### 5.14 $^{13}C$ NMR of compound 4a (101 MHz, DMSO-$d_6$):

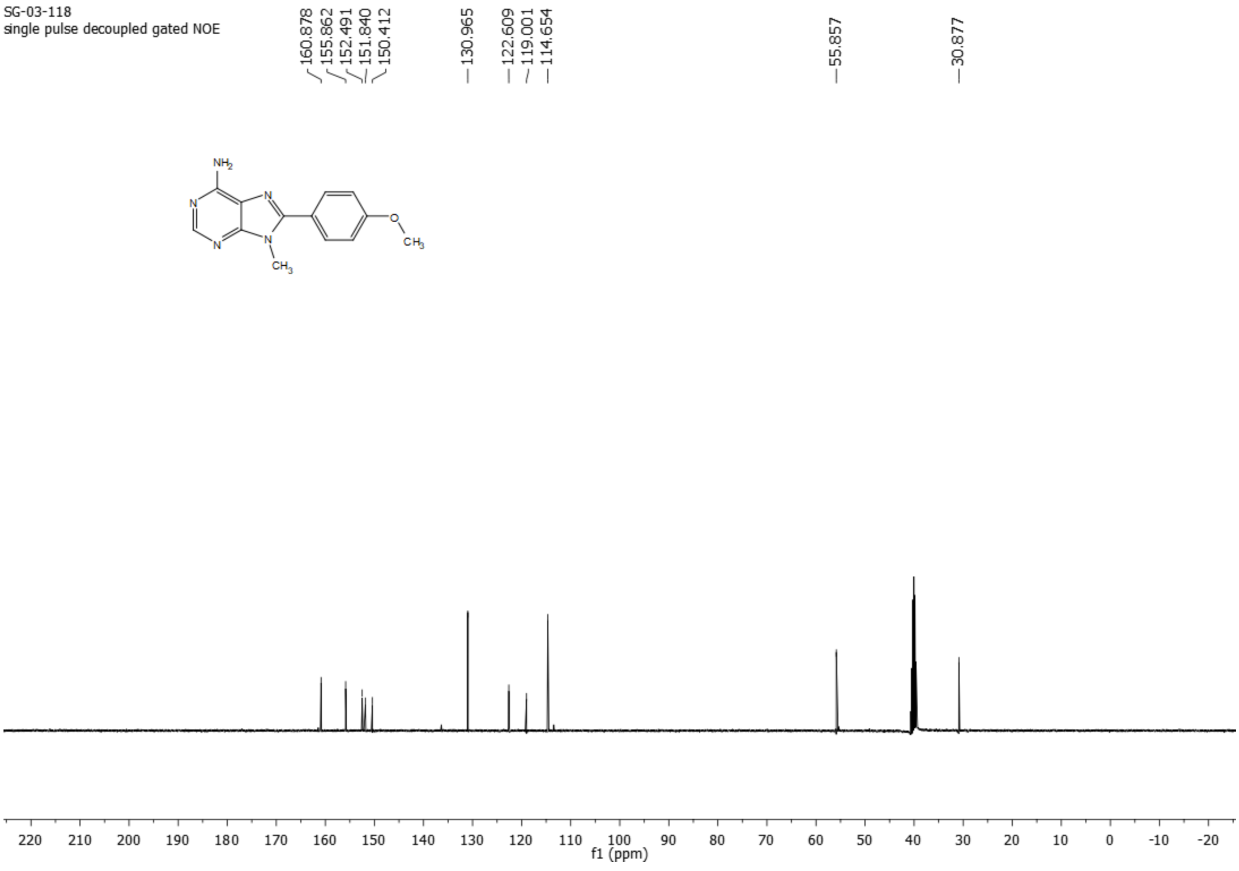

#### 5.15 $^{1}$H NMR of compound 4b (400 MHz, Chloroform-*d*):

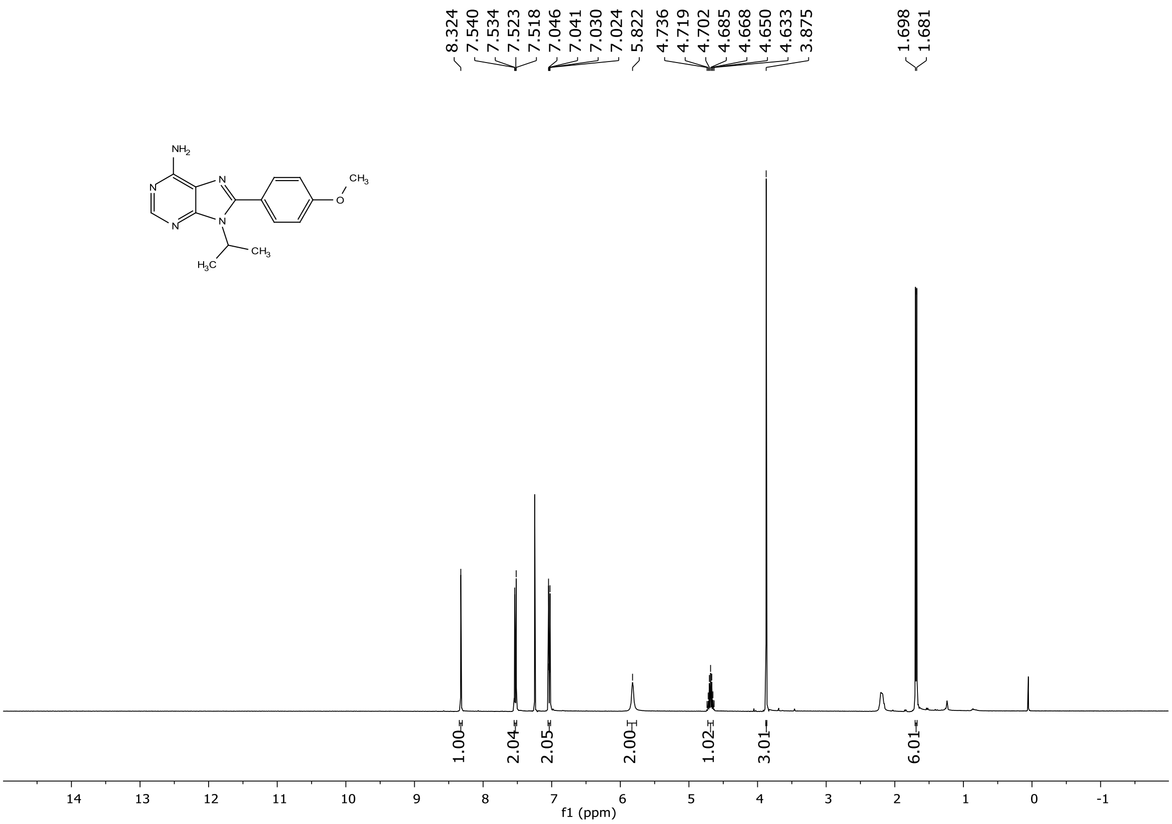

#### 5.16 $^{13}$C NMR of compound 4b (101 MHz, Chloroform-*d*):

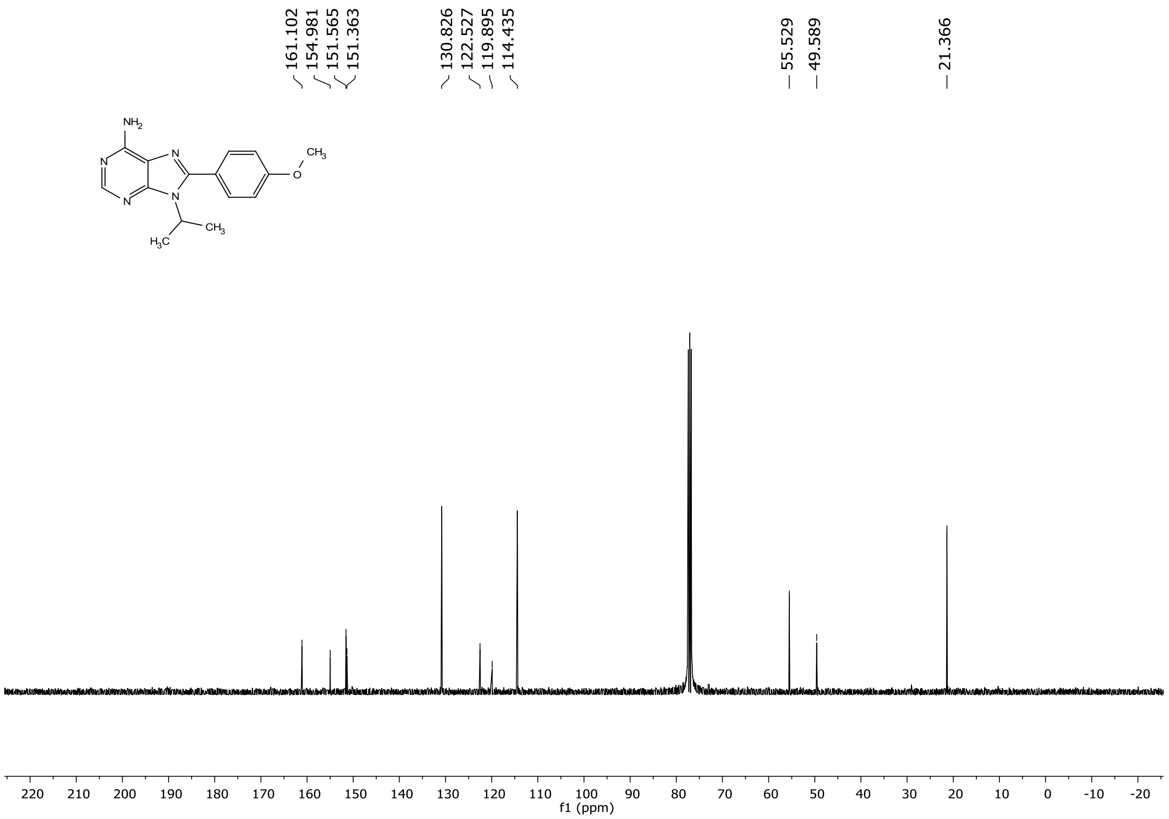

#### 5.17 $^{1}$H NMR of compound 4c (400 MHz, Chloroform-*d*):

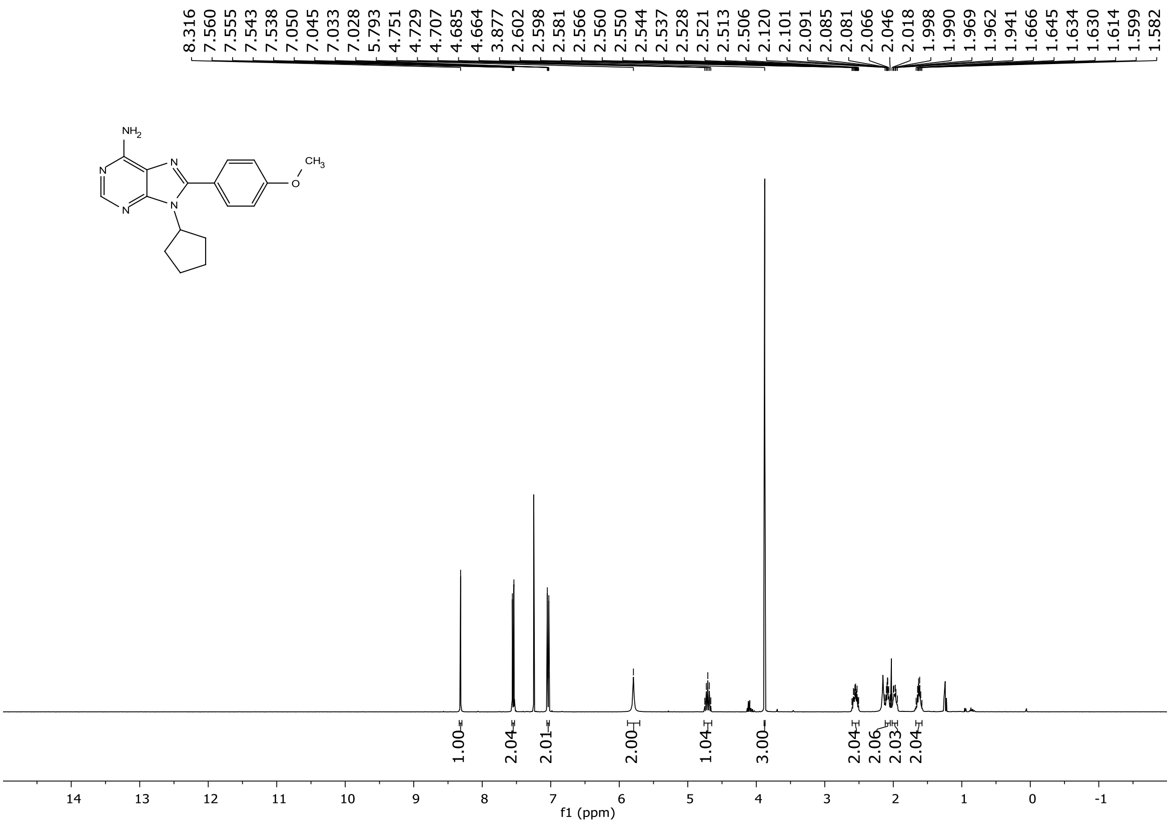

#### 5.18 $^{13}$C NMR of compound 4c (101 MHz, Chloroform-*d*):

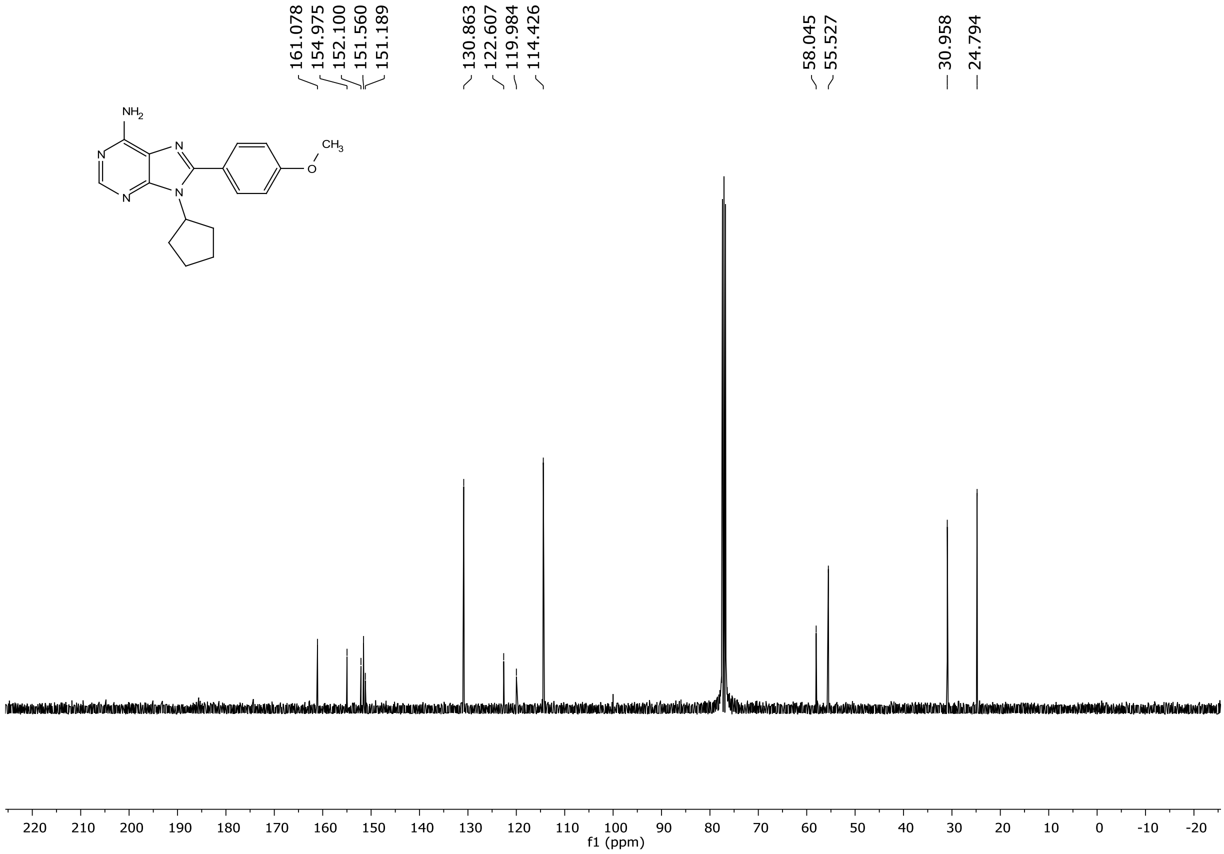

#### 5.19 [1]H NMR of compound 4d (400 MHz, Chloroform-*d*):

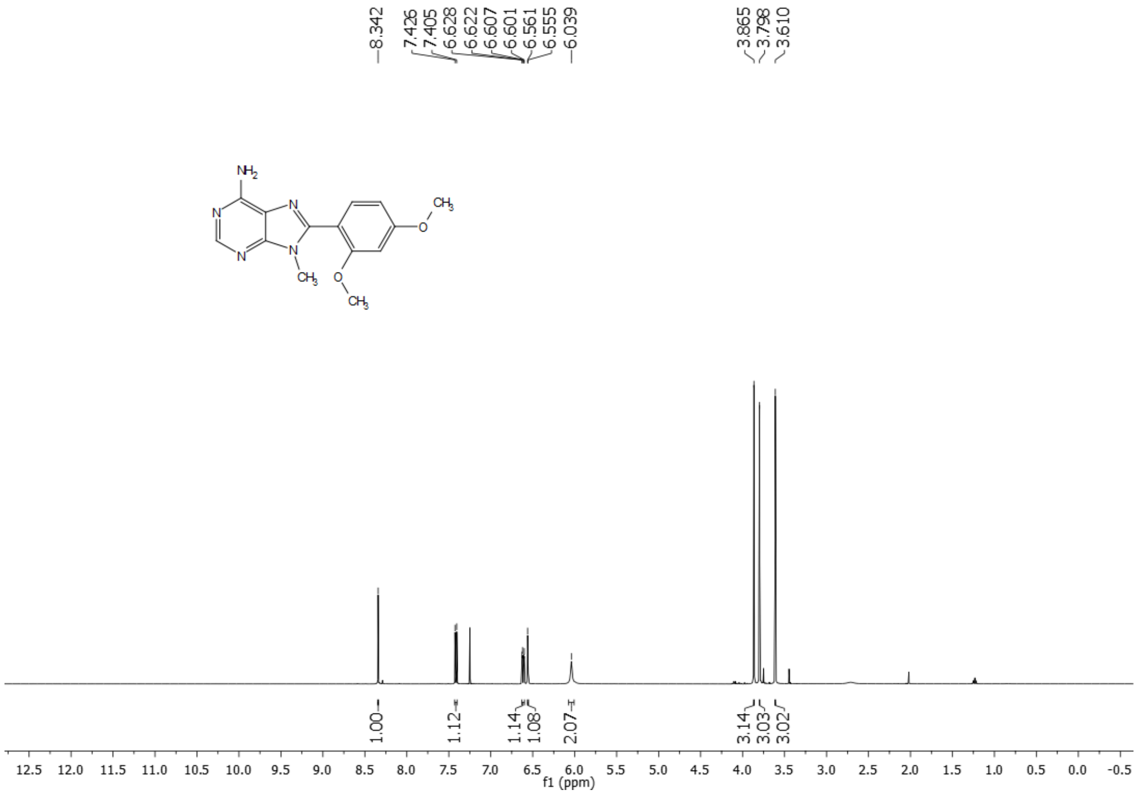

#### 5.20 [13]C NMR of compound 4d (101 MHz, Chloroform-*d*):

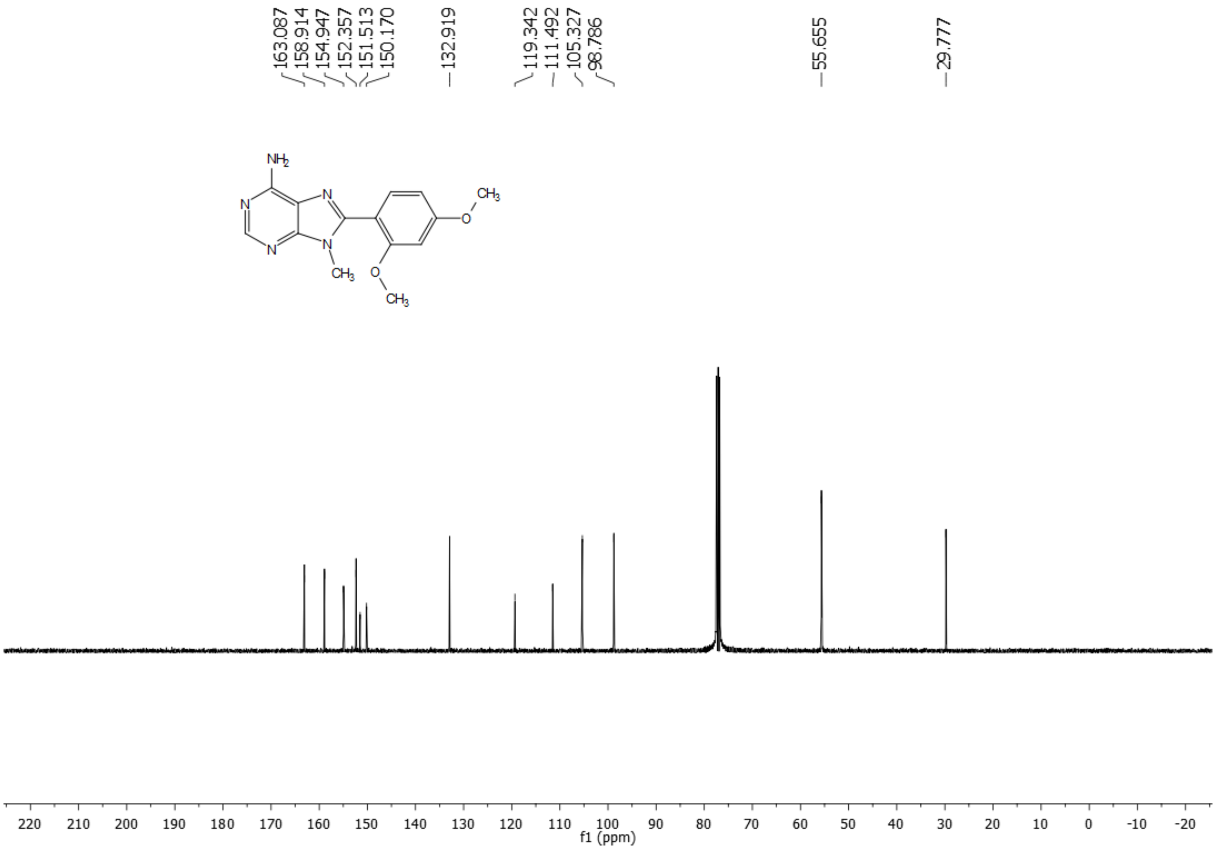

#### 5.21 $^{1}$H NMR of compound 4f (400 MHz, DMSO-$d_6$):

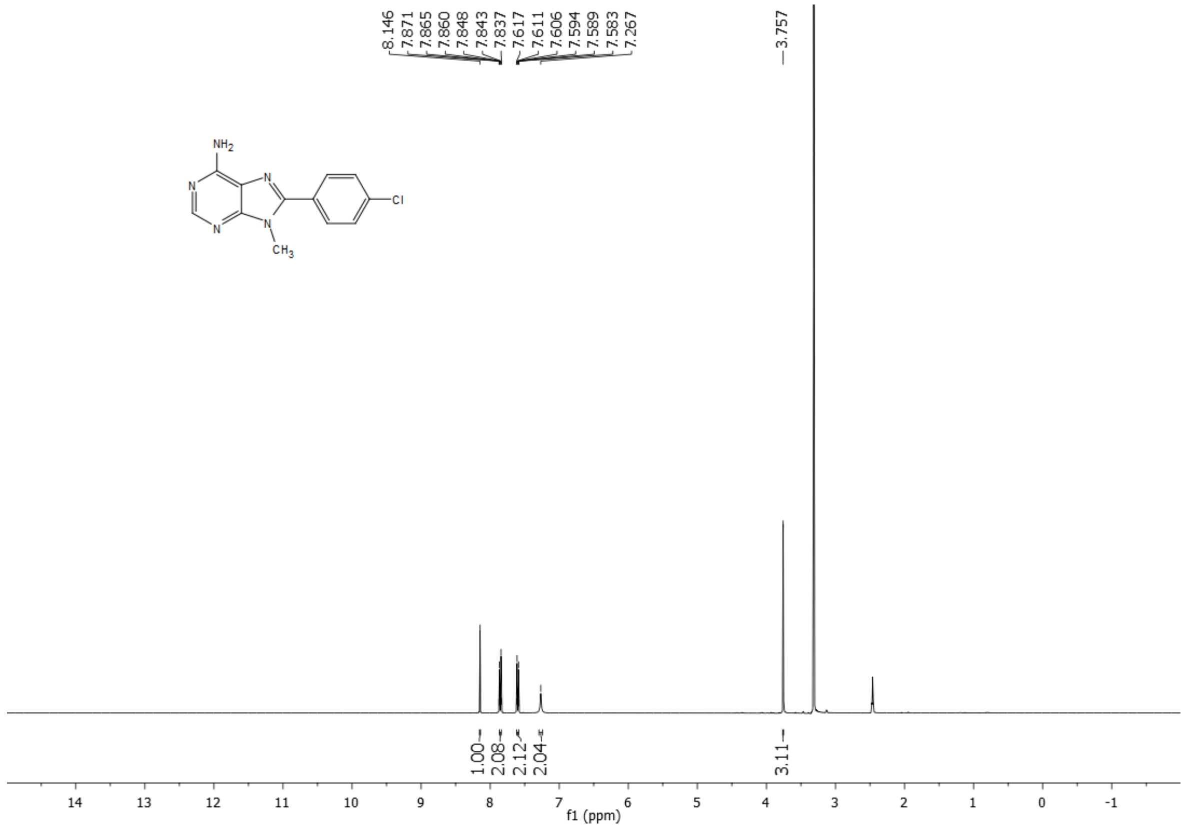

#### 5.22 $^{13}$C NMR of compound 4f (101 MHz, DMSO-$d_6$):

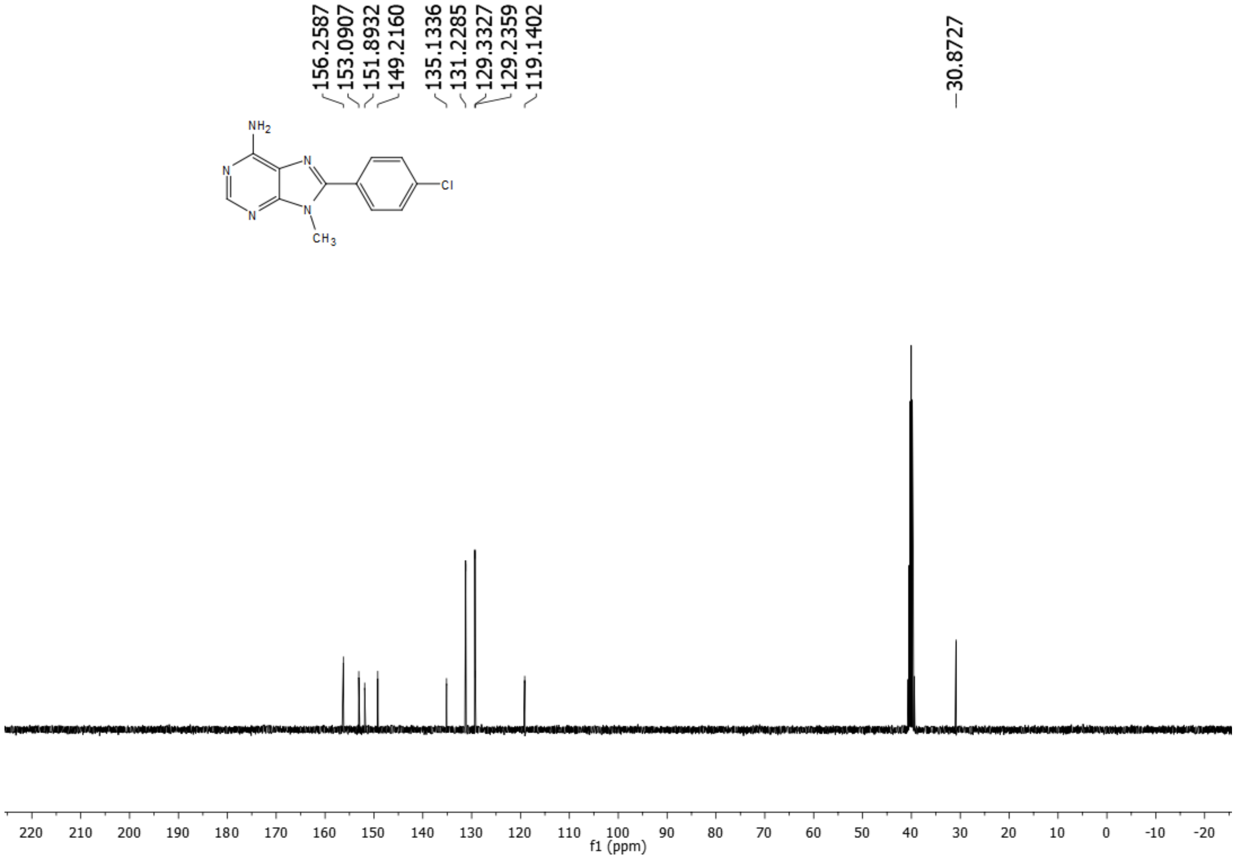

#### 5.23 $^{1}$H NMR of compound AS1 (400 MHz, Chloroform-*d*):

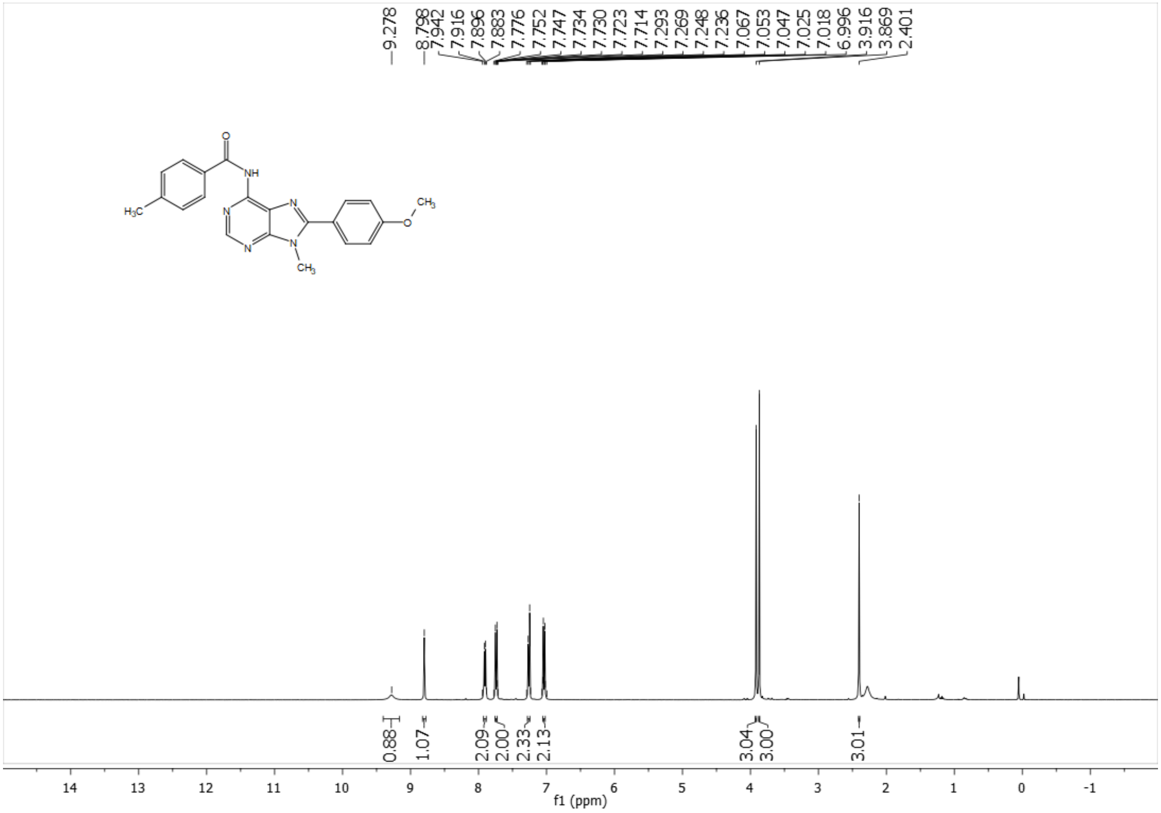

#### 5.24 $^{13}$C NMR of compound AS1 (101 MHz, Chloroform-*d*):

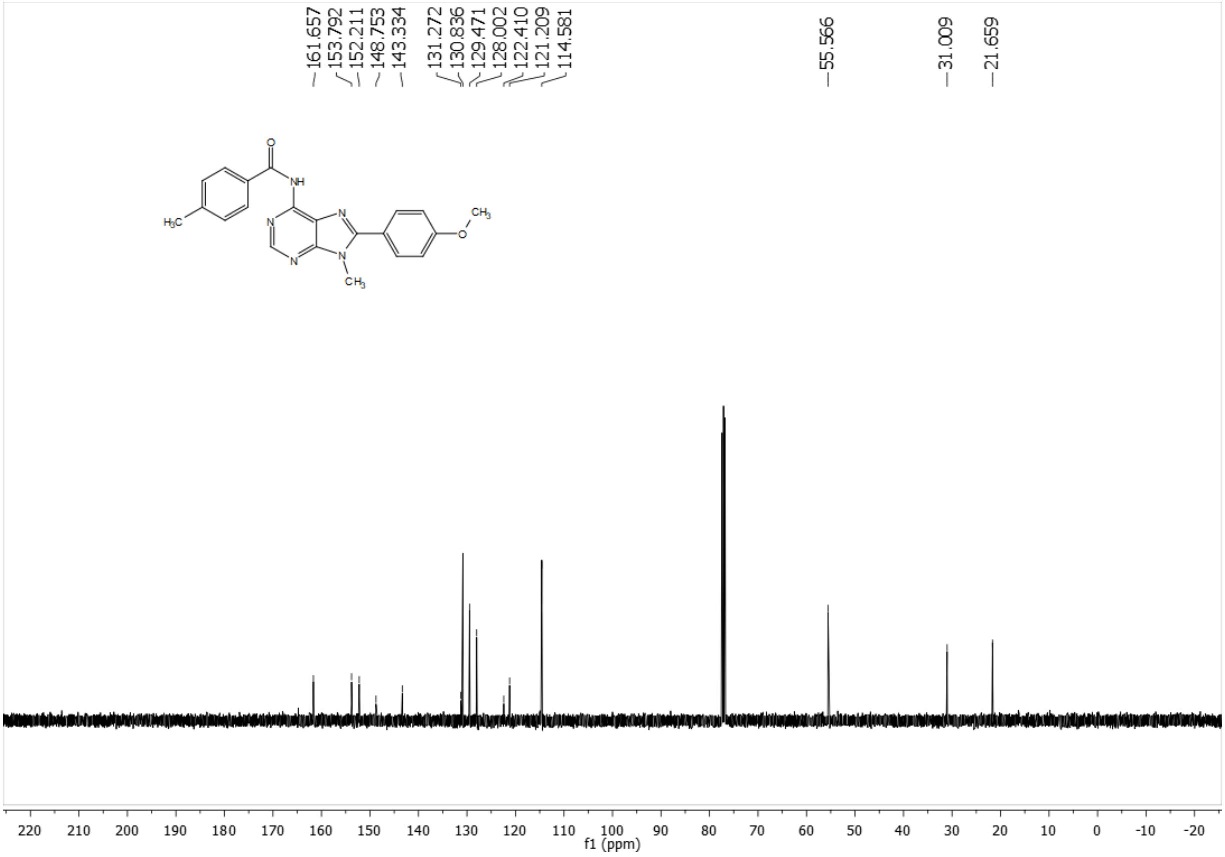

#### 5.25 $^{1}$H NMR of compound AS2 (400 MHz, Chloroform-*d*):

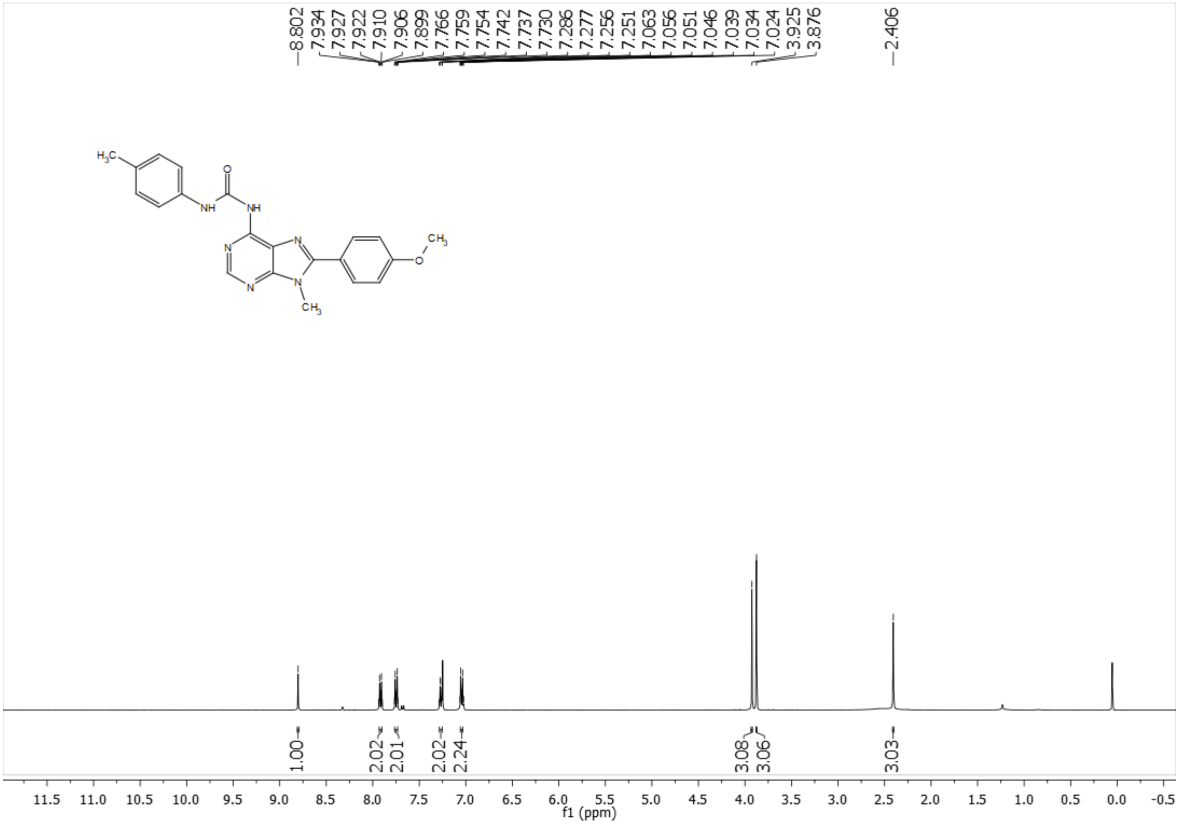

#### 5.26 $^{13}$C NMR of compound AS2 (101 MHz, Chloroform-*d*):

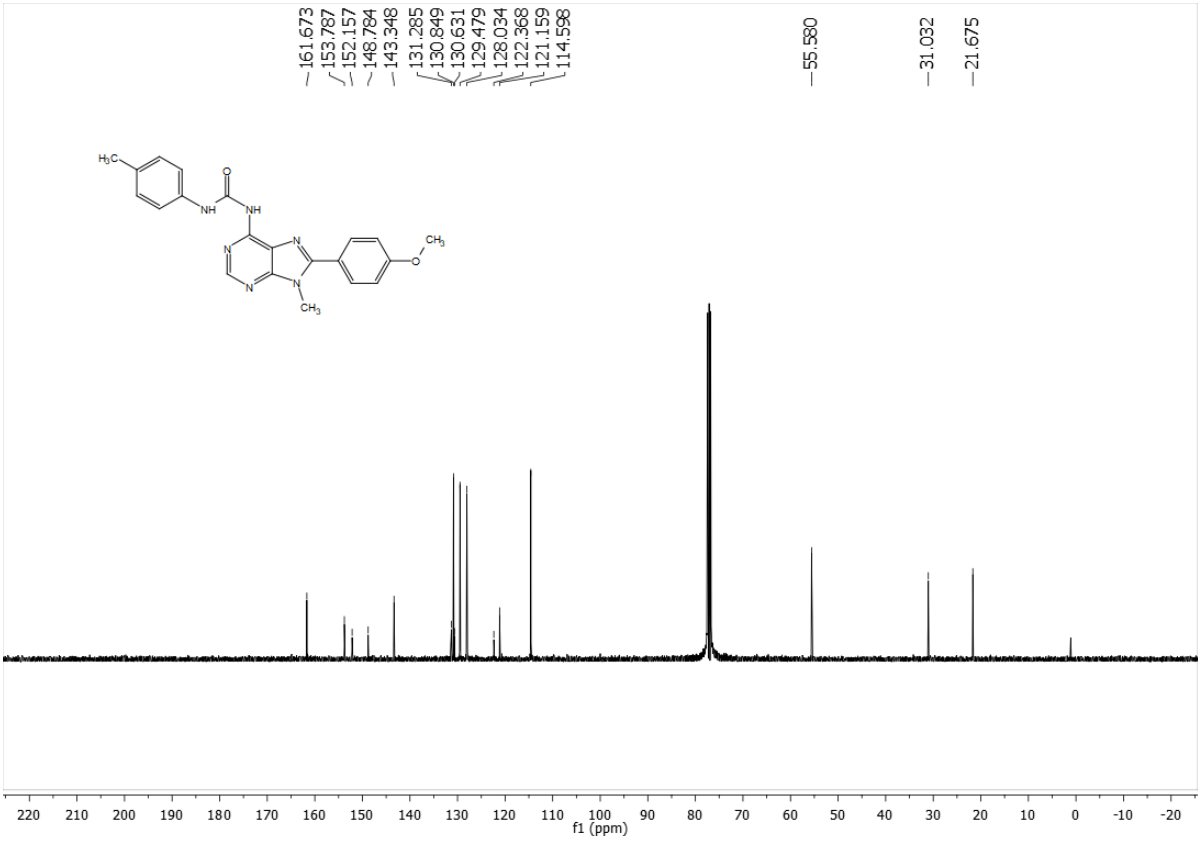

#### 5.27 $^{1}$H NMR of compound AS3 (400 MHz, Chloroform-*d*):

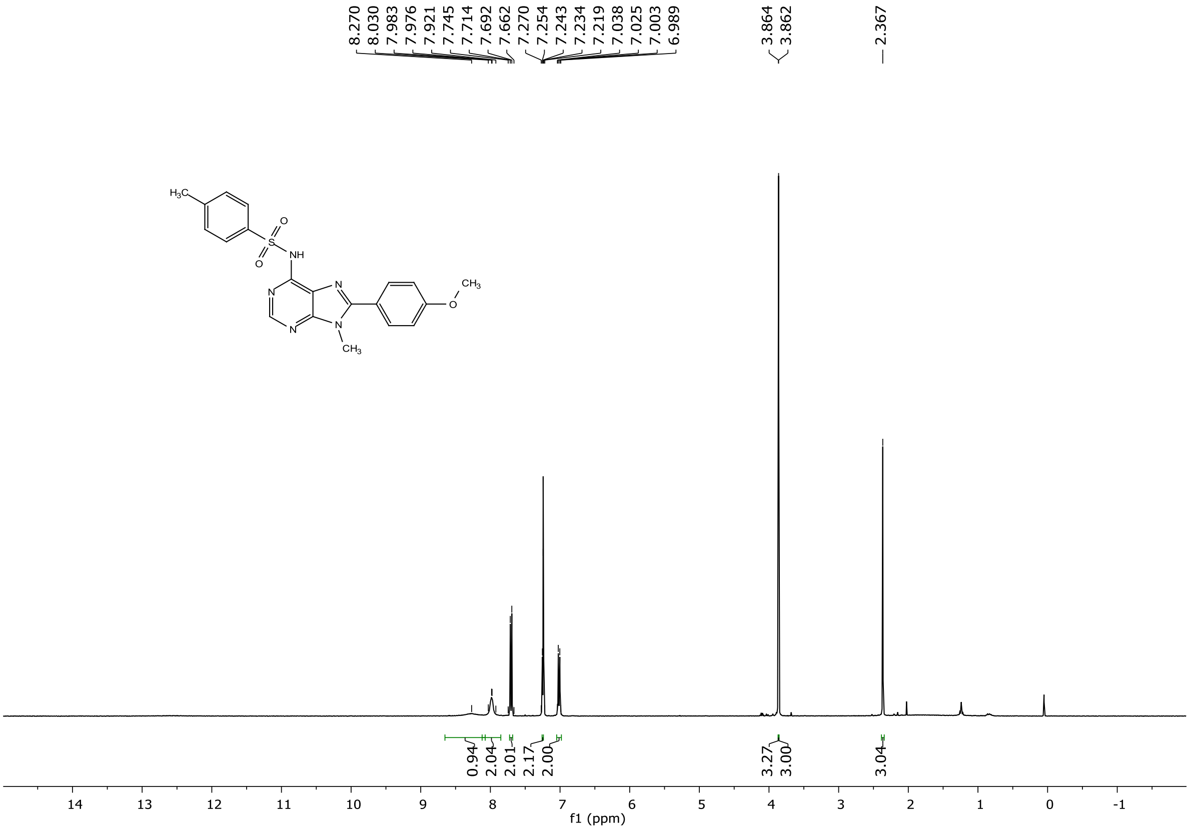

#### 5.28 $^{13}$C NMR of compound AS3 (101 MHz, Chloroform-*d*):

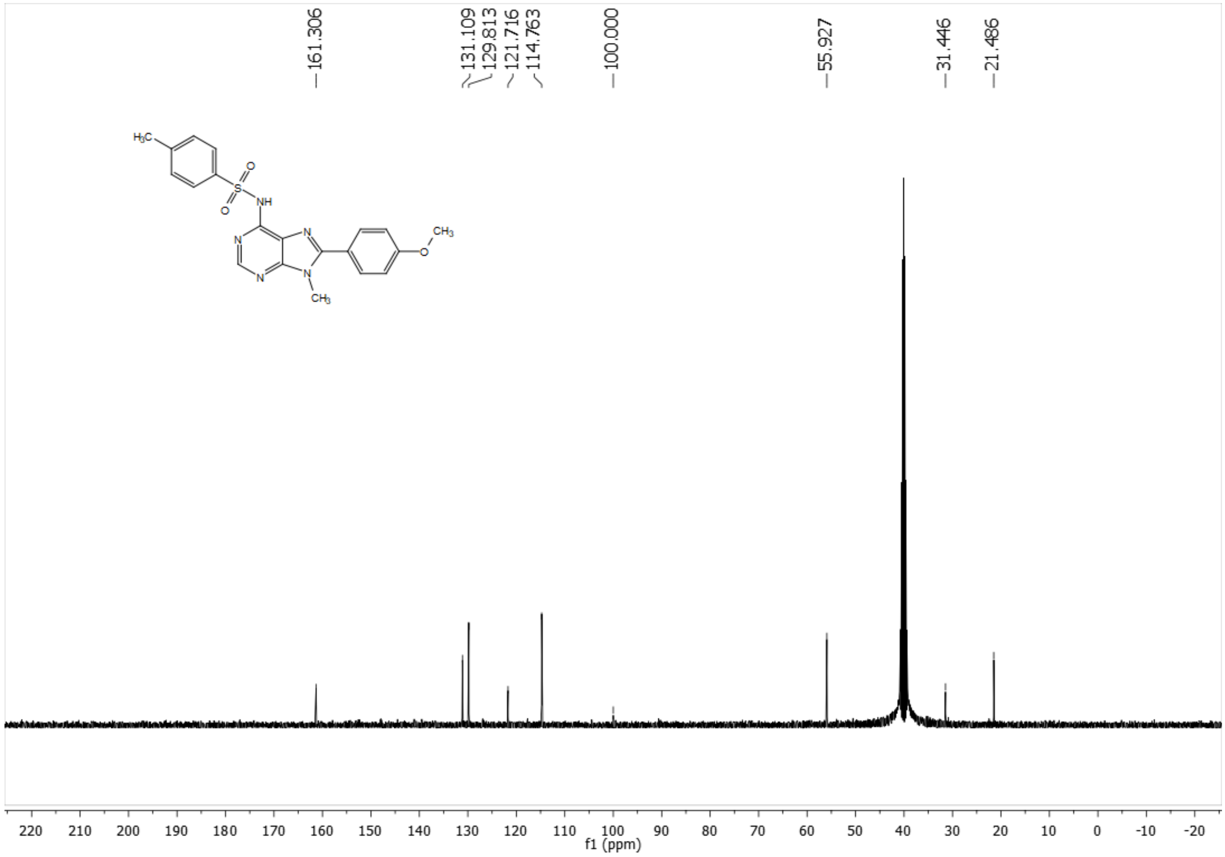

#### 5.29 $^{1}$H NMR of compound AS4 (400 MHz, Chloroform-*d*):

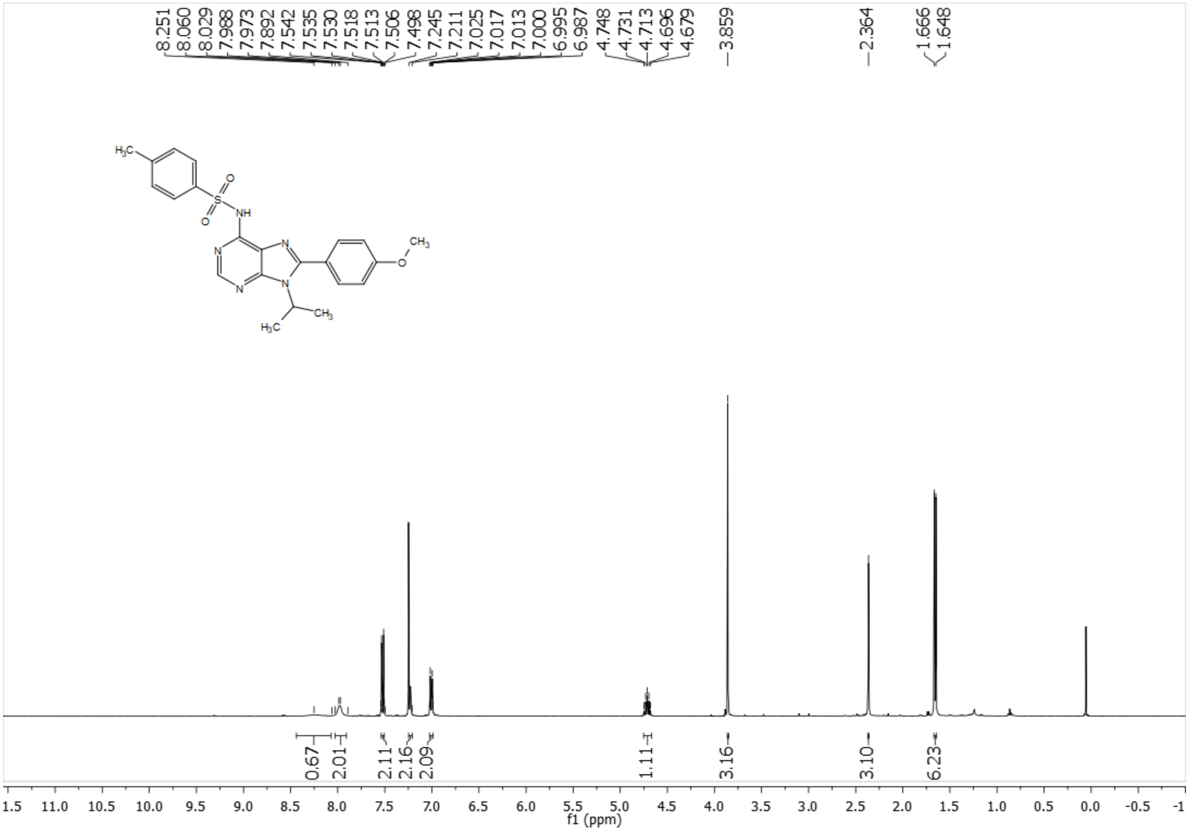

#### 5.30 $^{13}$C NMR of compound AS4 (101 MHz, Chloroform-*d*):

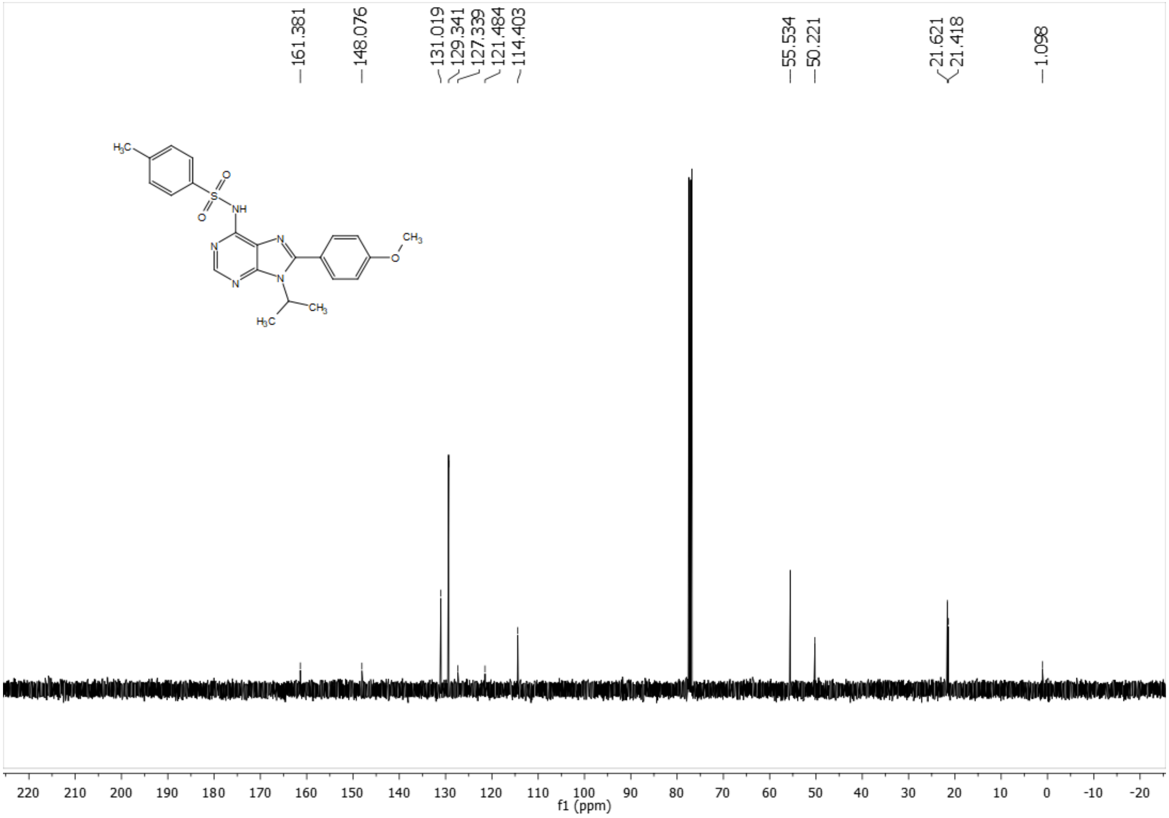

#### 5.31 $^{1}$H NMR of compound AS5 (400 MHz, Chloroform-*d*):

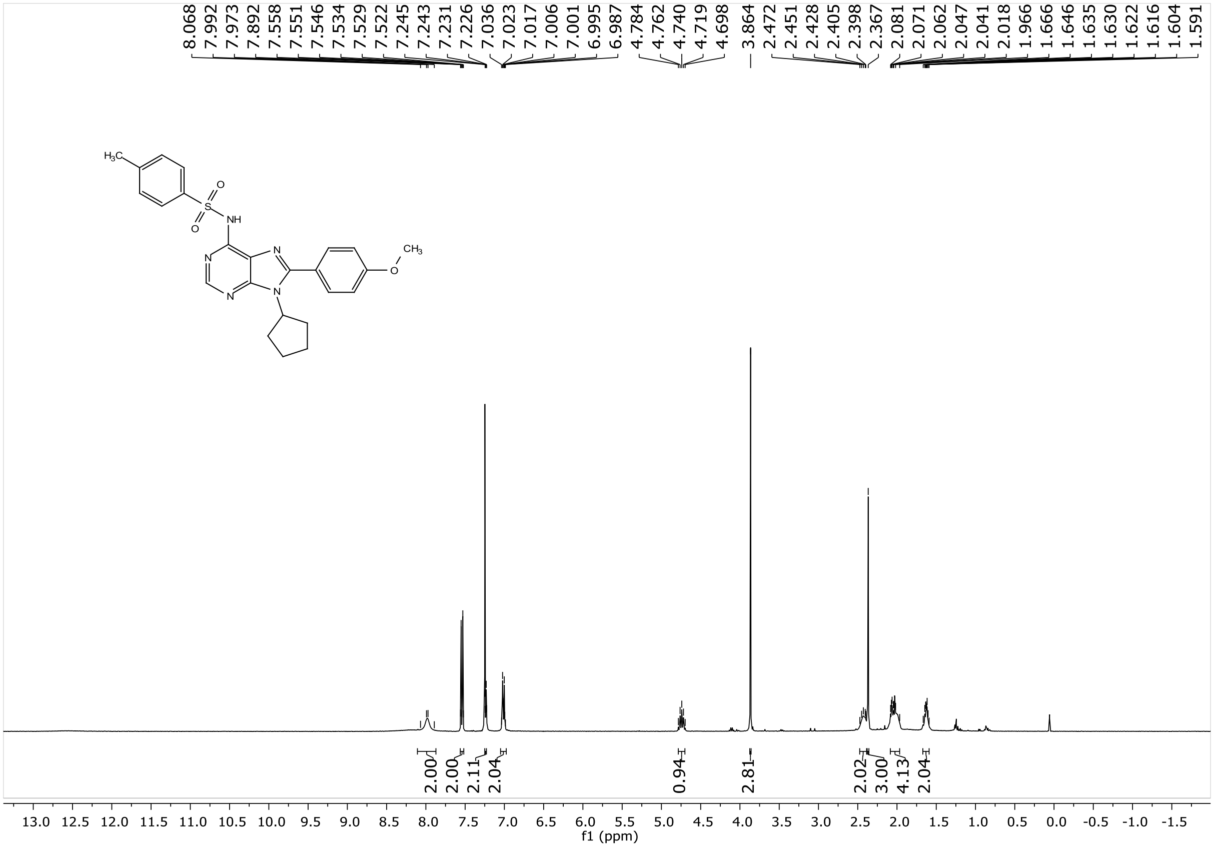

#### 5.32 $^{13}$C NMR of compound AS5 (101 MHz, Chloroform-*d*):

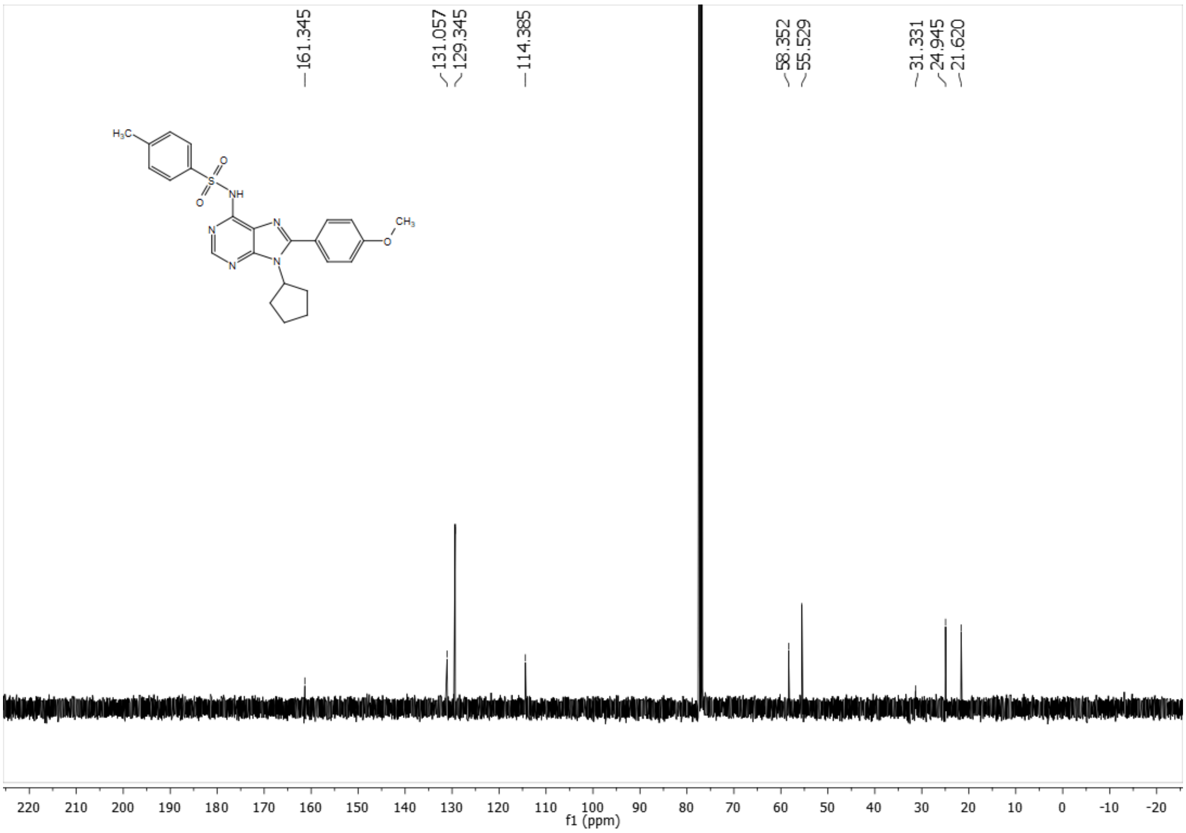

#### 5.33 [1]H NMR of compound AS6 (400 MHz, DMSO-$d_6$):

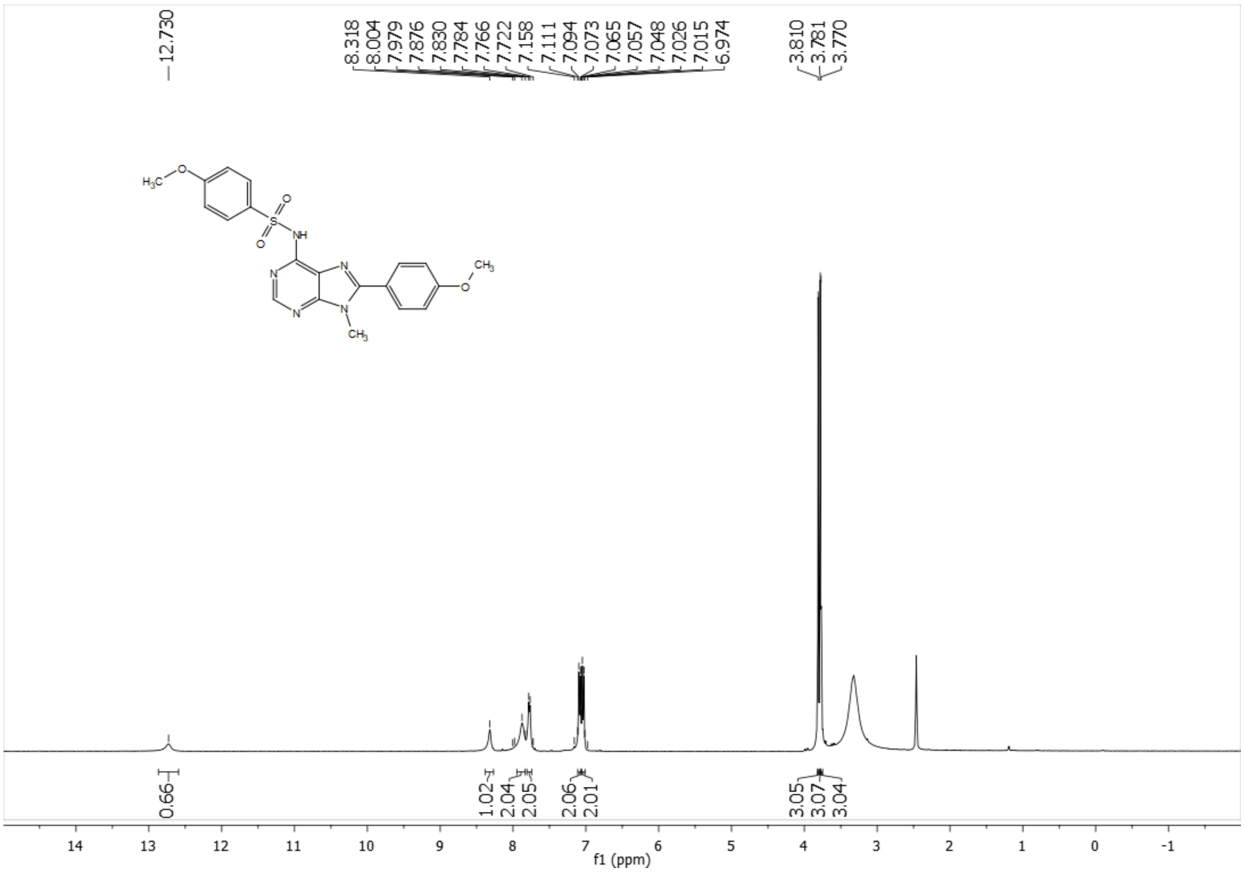

#### 5.34 [13]C NMR of compound AS6 (101 MHz, DMSO-$d_6$):

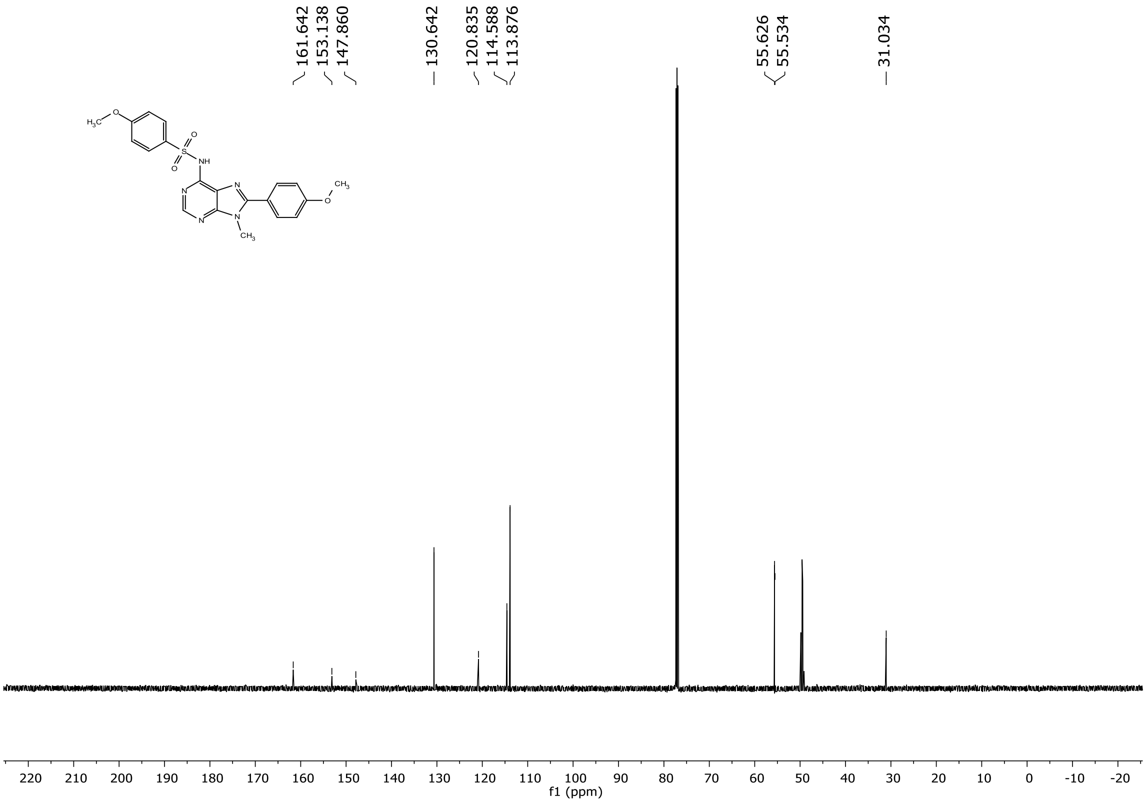

#### 5.35 $^{1}$H NMR of compound AS7 (400 MHz, DMSO-$d_6$):

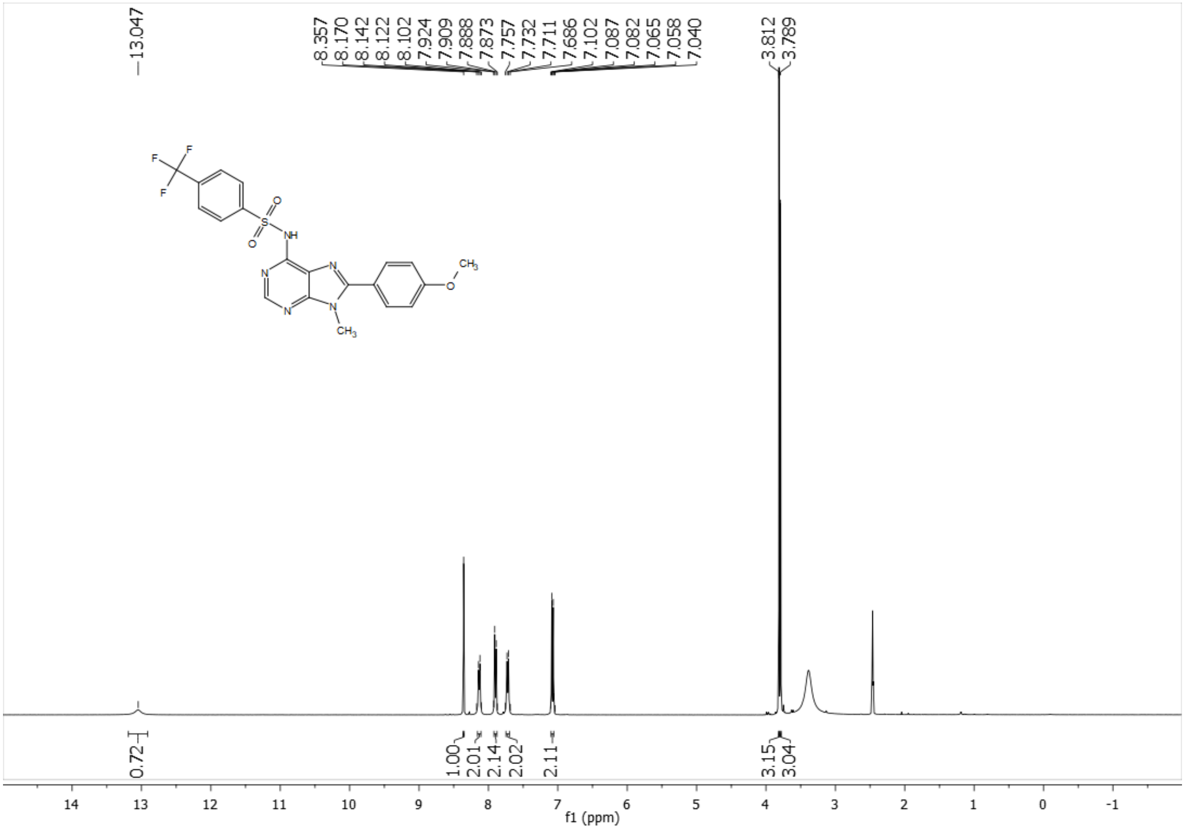

#### 5.36 $^{13}$C NMR of compound AS7 (101 MHz, DMSO-$d_6$):

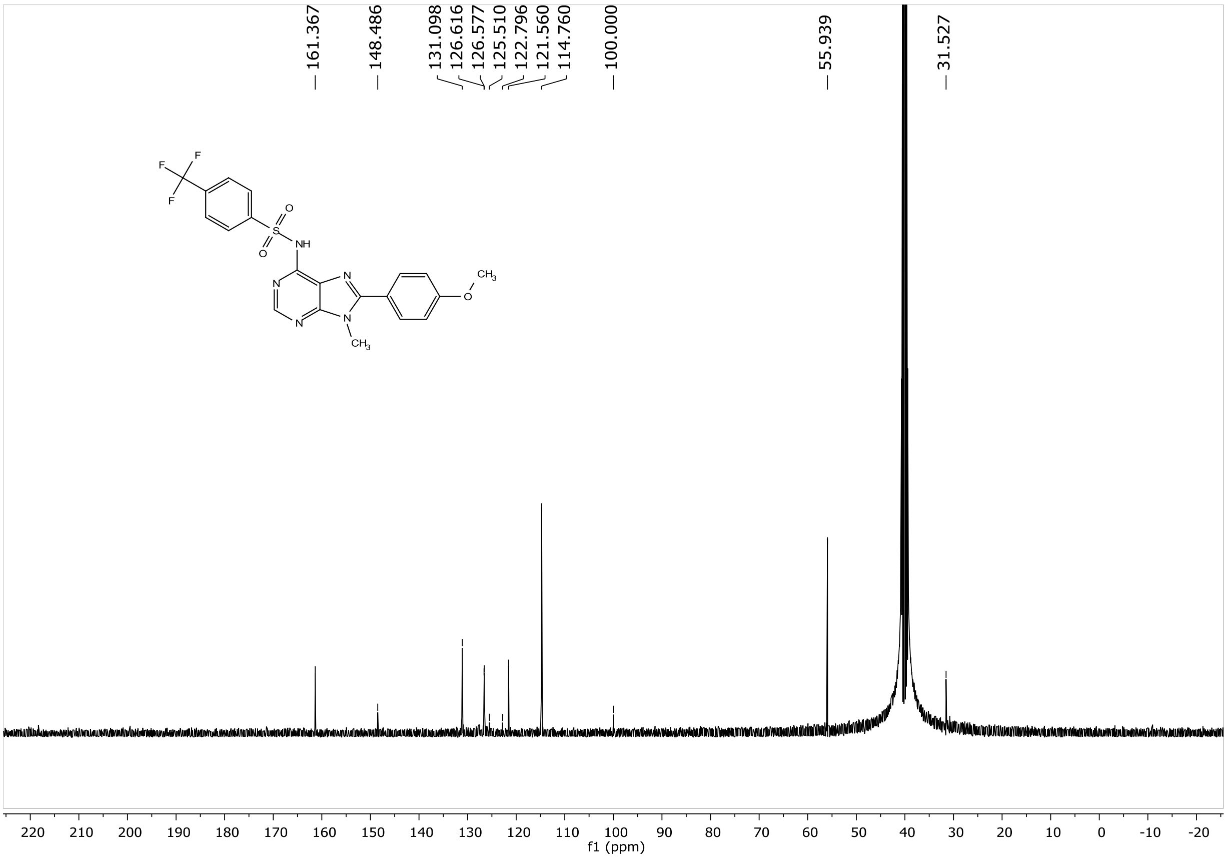

#### 5.37 [1]H NMR of compound AS8 (400 MHz, Chloroform-*d*):

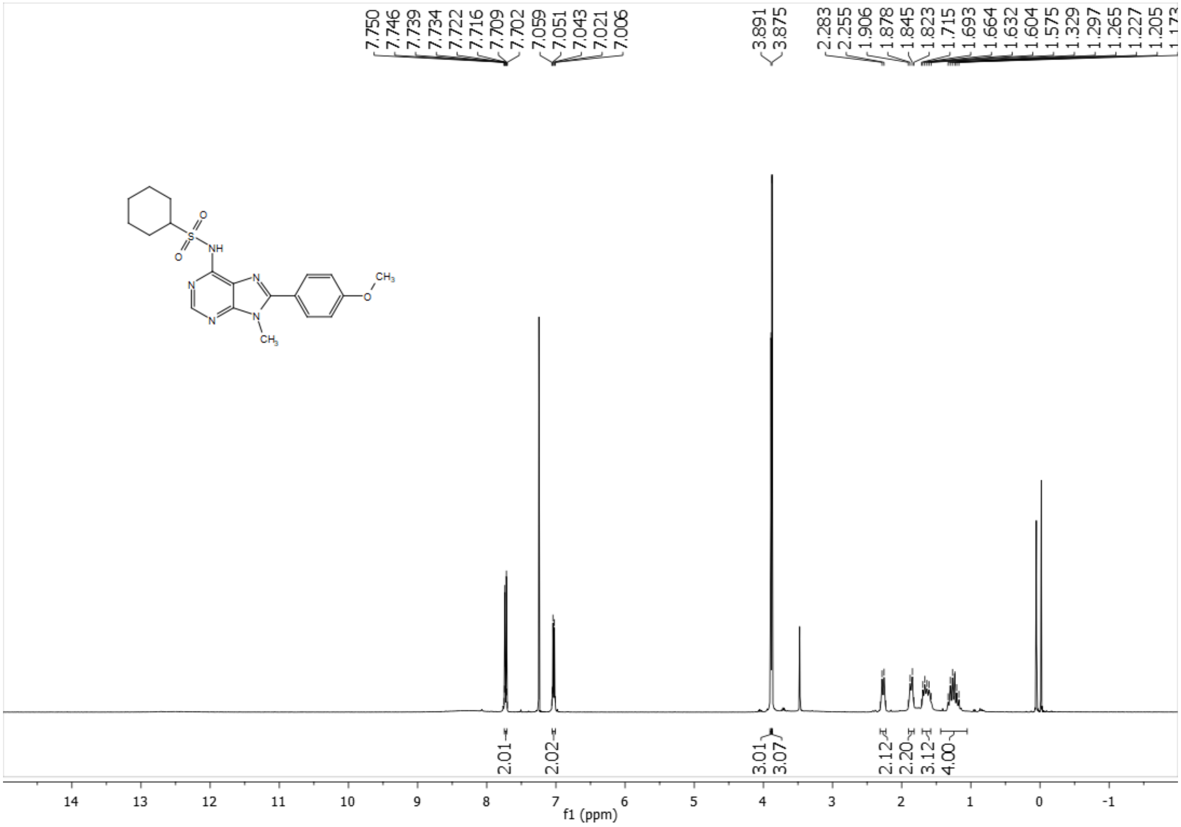

#### 5.38 [13]C NMR of compound AS8 (101 MHz, Chloroform-*d*):

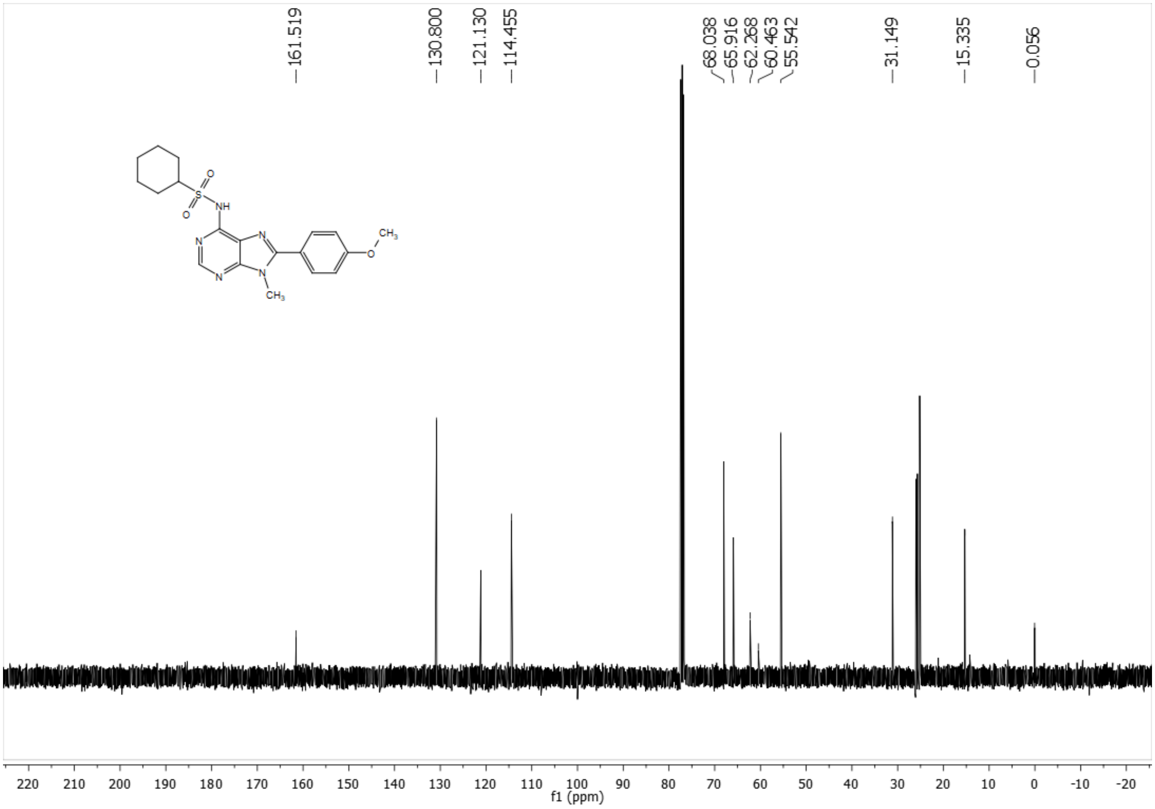

#### 5.39 $^{1}$H NMR of compound AS9 (400 MHz, Chloroform-*d*):

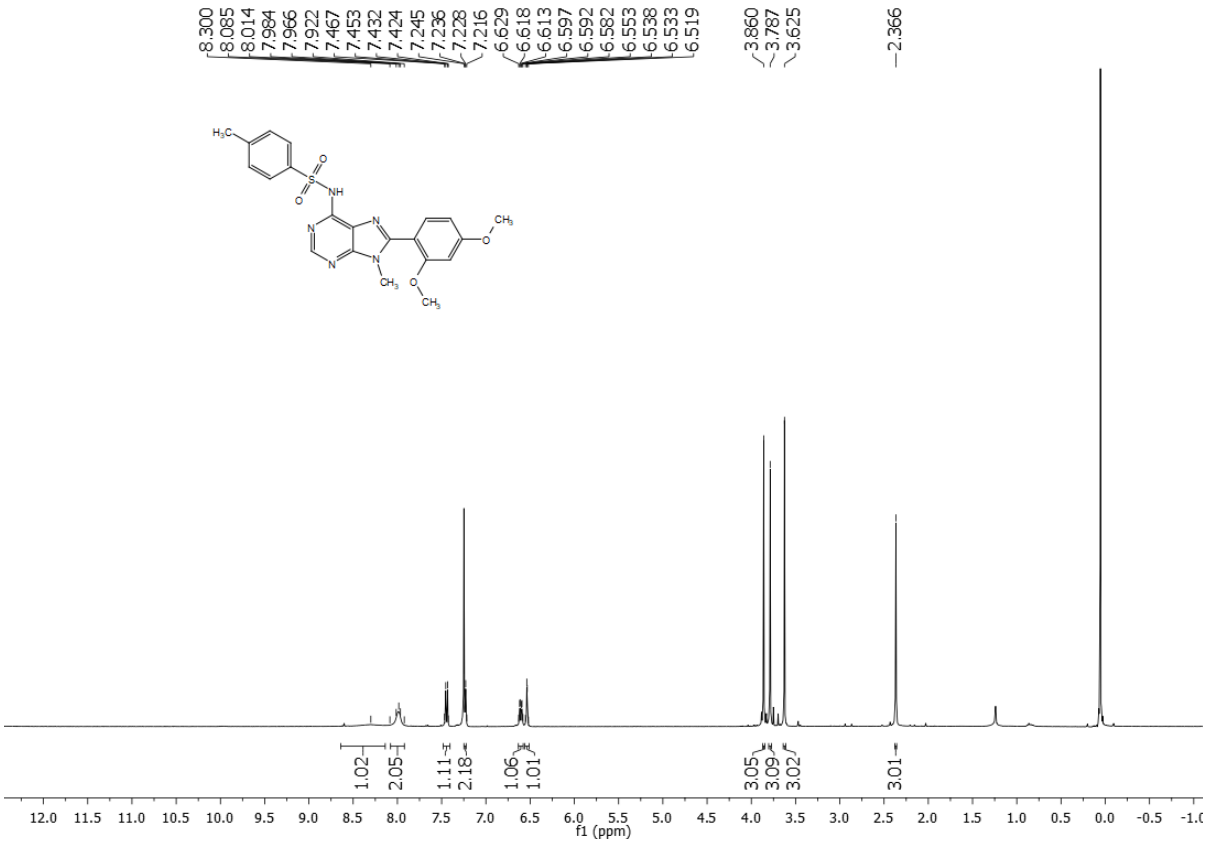

#### 5.40 $^{13}$C NMR of compound AS9 (101 MHz, Chloroform-*d*):

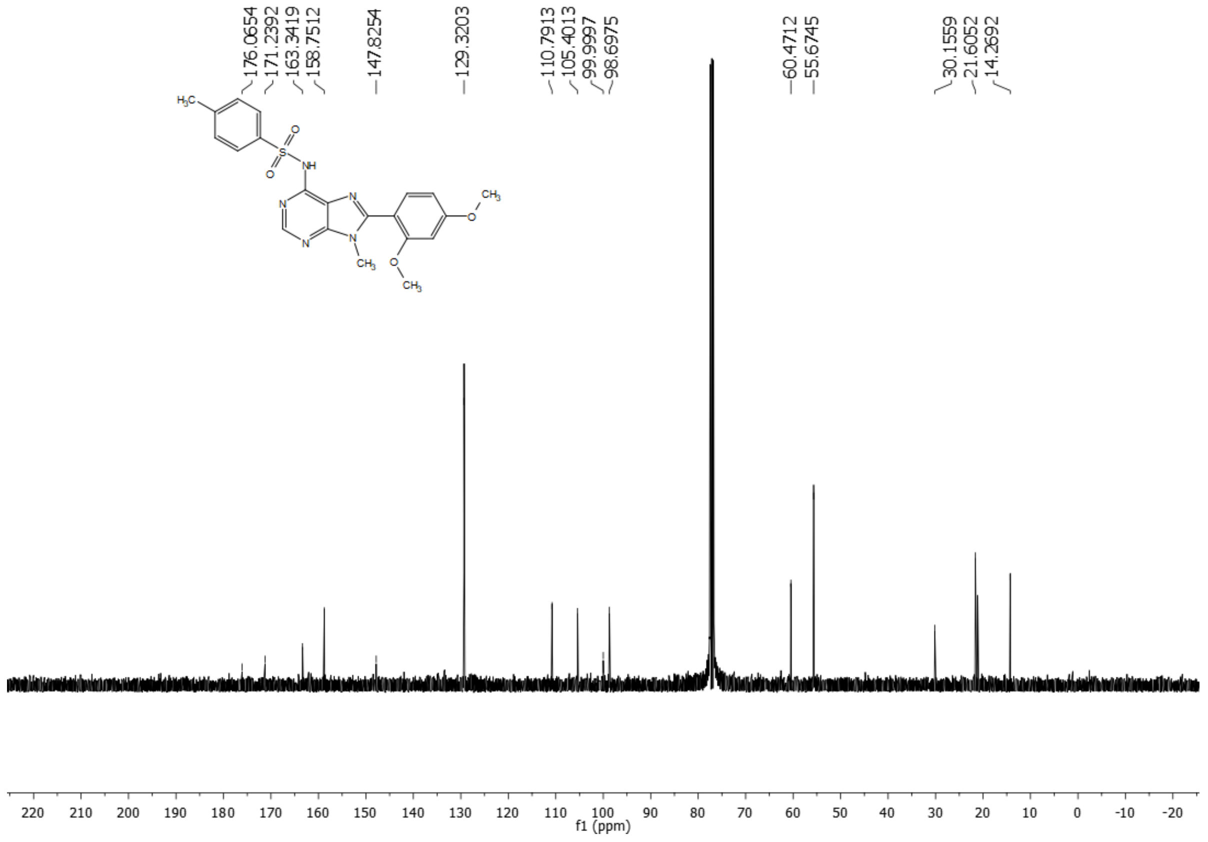

#### 5.41 $^{1}$H NMR of compound AS10 (600 MHz, Chloroform-*d*):

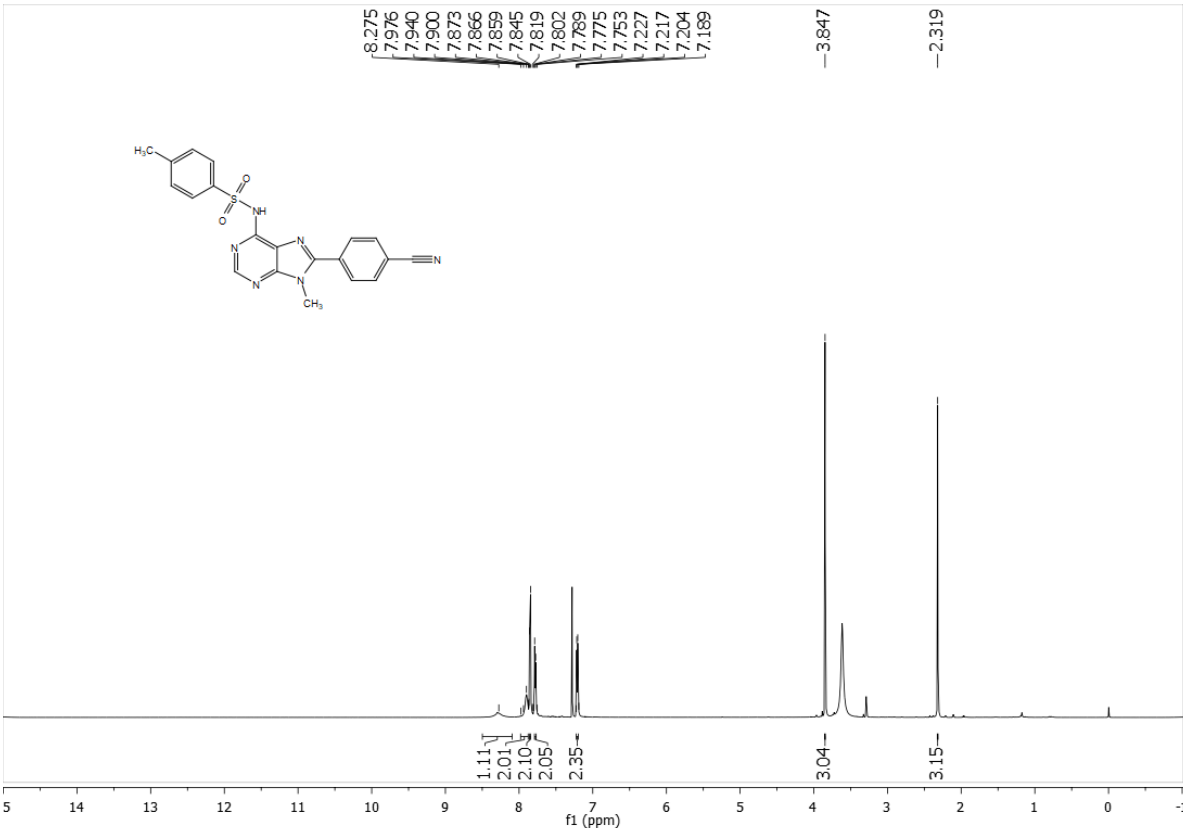

#### 5.42 $^{13}$C NMR of compound AS10 (101 MHz, Chloroform-*d*):

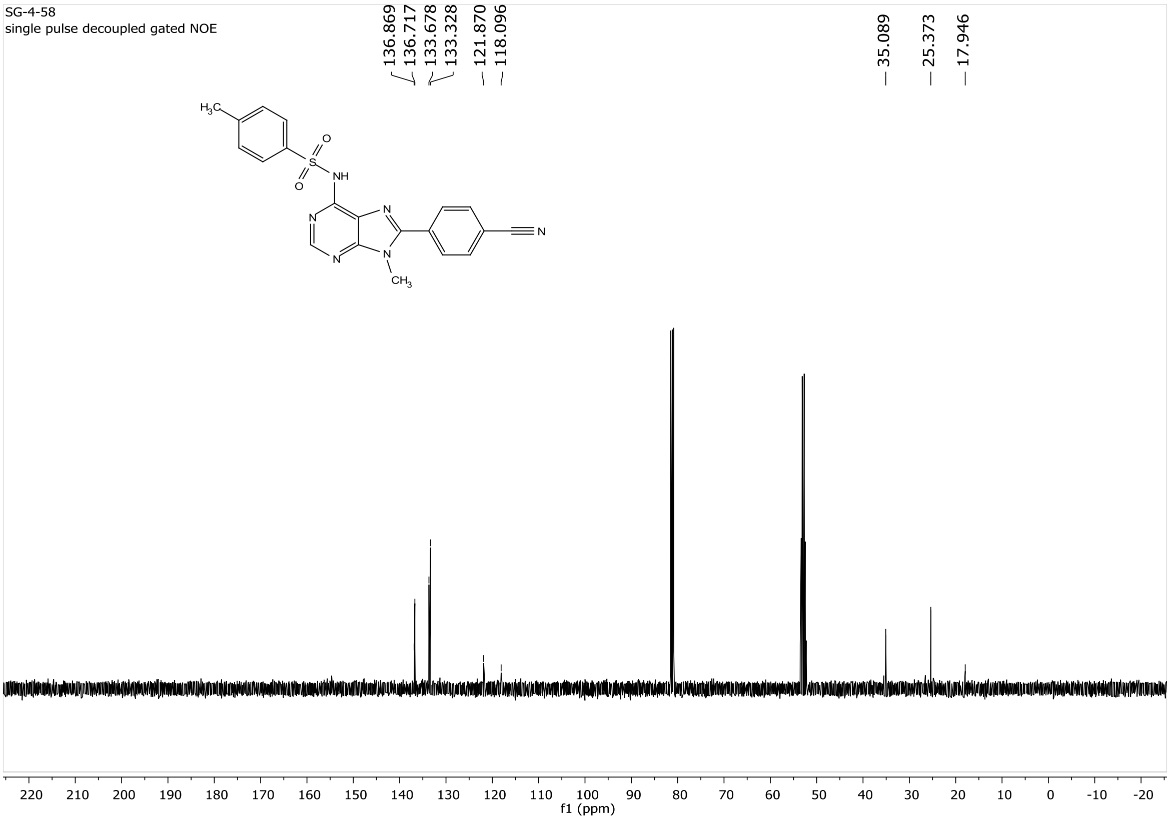

#### 5.43 $^{1}$H NMR of compound AS11 (400 MHz, Chloroform-*d*):

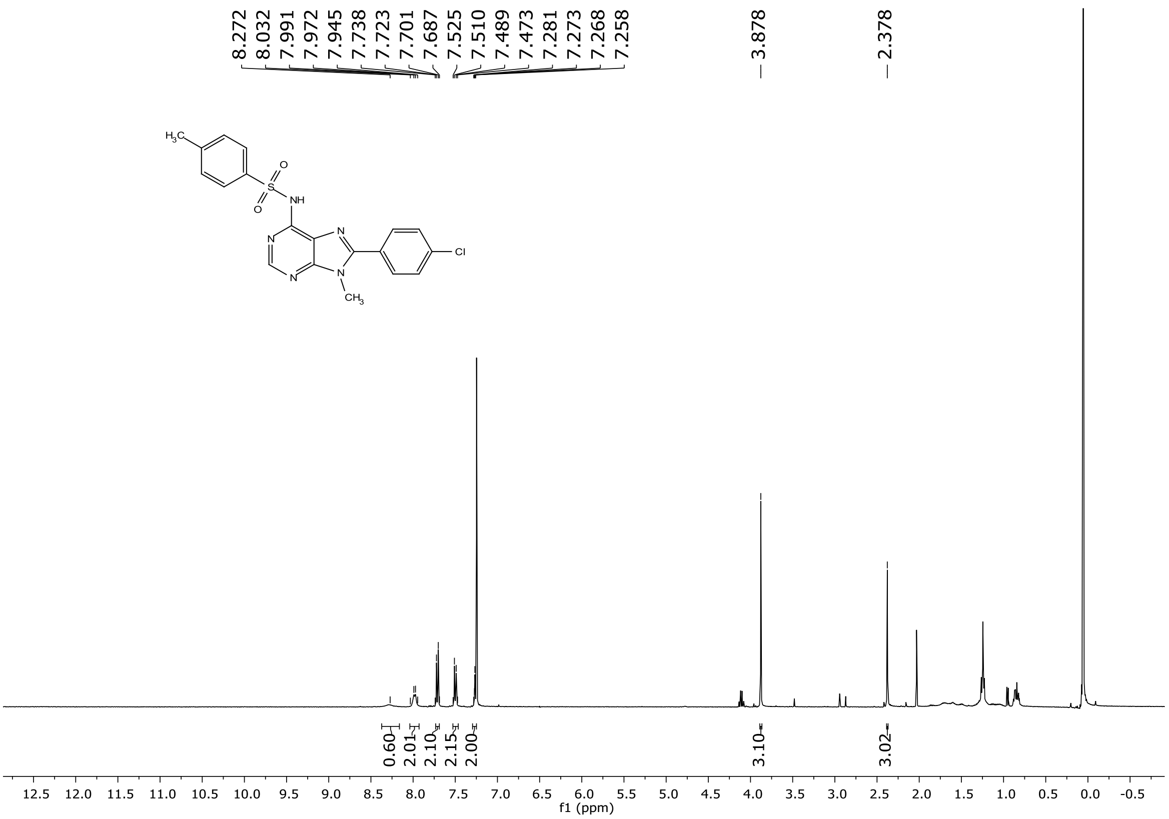

#### 5.44 $^{13}$C NMR of compound AS11 (101 MHz, DMSO-$d_6$):

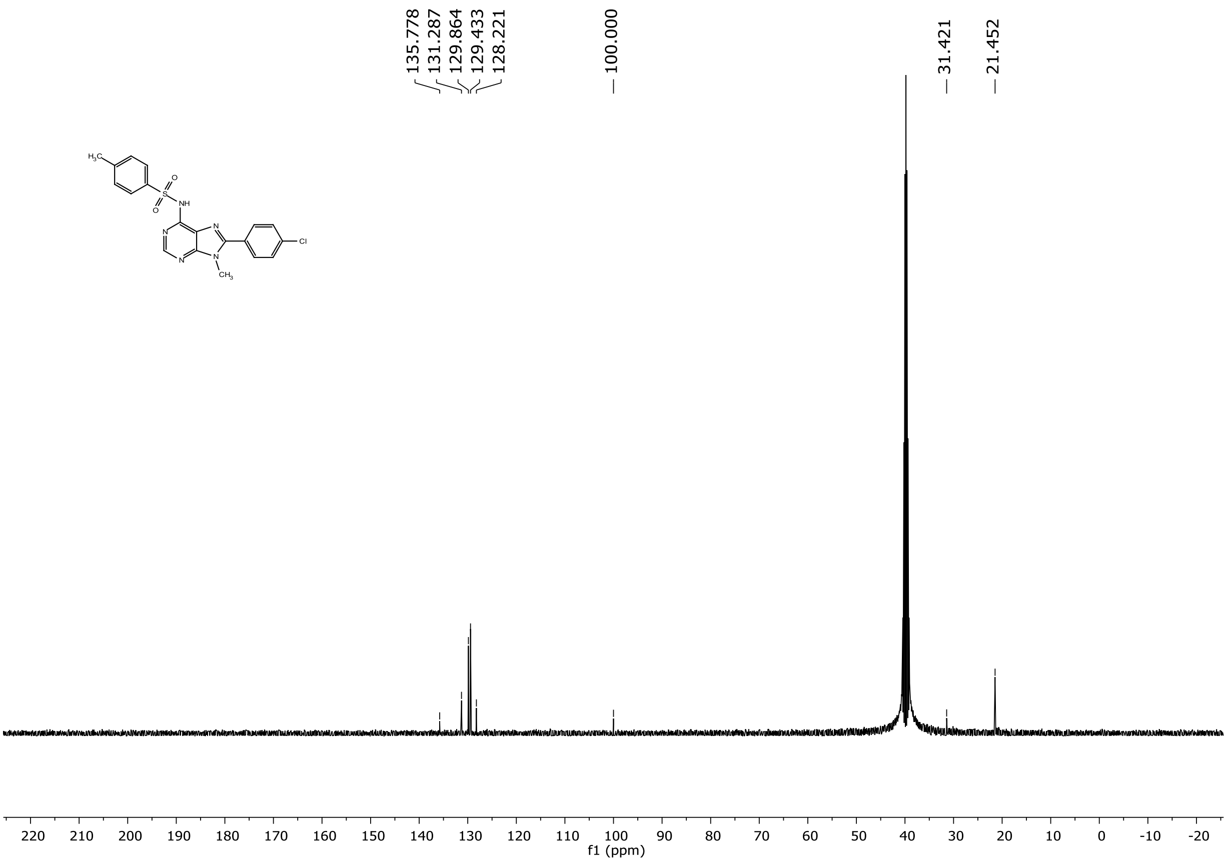
